# WWOX deficiency impairs neurogenesis and neuronal function in human organoids

**DOI:** 10.1101/2024.12.22.630016

**Authors:** Daniel J. Steinberg, Asia Zonca, Dania Abdellatif, Idan Rosh, Irina Kustanovich, Osama Hidmi, Kian Maroun, Shani Stern, Jose Davila-Velderrain, Rami I. Aqeilan

**Author notes:** Corresponding authors and lead contacts. These authors contributed equally to this work (Daniel J. Steinberg, Asia Zonca, Dania Abdellatif).

## Abstract

WOREE and SCAR12 syndromes are rare neurodevelopmental disorders caused by WWOX mutations, severely impairing brain development. The pleiotropic nature of WWOX complicates identifying specific mechanisms. Using neural organoids and single-cell transcriptomics, we identified radial glial cells (RGs) as preferentially affected, with disrupted cell cycle dynamics leading to an accumulation of cells in the G2/M and S phases, overexpression of the proto-oncogene MYC, and concomitant reduction in neuronal generation. Patient-derived organoids exhibited milder phenotypes compared to knockout organoids, showing functional neuronal impairments like hyperexcitability and delayed differentiation rather than RG dysfunction. Remarkably, gene therapy restored neuronal function, normalizing hyperexcitability and promoting maturation, without disturbing RG populations. We propose a model in which WWOX mutations impair neurogenesis via RG through cell-type specific dysregulation of the MYC and Wnt signaling pathways. These insights highlight potential therapeutic strategies for WWOX-related disorders and open avenues for interventions targeting these key molecular pathways.

**Teaser:** WWOX mutations disrupt radial glial function and neurogenesis via MYC dysregulation, with gene therapy offering targeted restoration.

## Introduction

WWOX-related epileptic encephalopathy (WOREE syndrome) and Spinocerebellar ataxia autosomal recessive type 12 (SCAR12 syndrome) are rare and devastating neurodevelopmental disorders linked to mutations in the WW-domain containing oxidoreductase *(WWOX)* gene (Abdel-Salam *et al*, 2014; Mallaret *et al*, 2014; Mignot *et al*, 2015). The typical symptoms for both disorders include early-onset epilepsy, cognitive impairments, global developmental delay, structural brain abnormalities and atrophy, and in some cases, premature death (Banne *et al*, 2021; Piard *et al*, 2019). These WWOX-related neurodevelopmental disorders display gradual severity in genotype-phenotype associations (Banne *et al*, 2021). First, SCAR12 syndrome has been diagnosed so far only in patients with missense mutations (Mallaret *et al*, 2014), while WOREE is commonly associated with nonsense mutations and copy number variations (Piard *et al*, 2019). Second, both survival and time-to-seizure onset analyses for 75 individuals with WOREE syndrome showed worst prognosis in individuals with null-mutations, whose severity correlated with the number of null-alleles (Oliver *et al*, 2023). These two diseases also differ clinically; SCAR12 typically manifests in early childhood with mild to moderate global developmental delay, cerebellar ataxia, and generalized seizures that are often responsive to treatment, allowing most individuals to survive into adulthood and attain basic motor and verbal milestones (Mallaret *et al*, 2014; Banne *et al*, 2021; Piard *et al*, 2019). Brain imaging in SCAR12 patients generally reveals mild hypoplasia of the cerebellar vermis and posterior white matter hyperintensities, with no other imaging abnormalities reported (Banne *et al*, 2021; Gribaa *et al*, 2007). In contrast, WOREE is a severe early-onset epileptic encephalopathy, presenting in the first months of life with drug-resistant seizures, profound developmental arrest, axial hypotonia, and early lethality (Banne *et al*, 2021; Piard *et al*, 2019; Oliver *et al*, 2023). Neuroimaging frequently shows diffuse brain atrophy, underdeveloped corpus callosum, cerebellar hypoplasia, and progressive white matter loss (Battaglia *et al*, 2023). The *WWOX* gene, mostly studied for its tumor suppressor effect, is known to influence a wide array of biological processes, often by functioning as an adaptor protein and transcriptional co-repressor (Abu-Remaileh *et al*, 2015), making its cellular effects context-dependent, varying according to the available binding partners. *WWOX* expression levels also vary during development. In human brain tissue samples WWOX transcription levels are relatively high during embryonic life and progressively decrease during fetal development until birth, after which expression increases gradually and remains high from adolescence onwards (Aldaz & Hussain, 2020). *In-vitro* study of brain development further demonstrated a similar pattern of differential *WWOX* expression during development and highlighted its connection to epilepsy (Gordon *et al*, 2021). However, whether similar developmental patterns of WWOX expression are recapitulated in experimental models of human neurodevelopment and how they relate to changes in cellular composition remains unclear.

Human neurogenesis is a highly dynamic process, involving diverse transitions between cellular populations. During embryonic development, neural induction gives rise to columnar neuroepithelial cells (NE), often referred to as the neuroectoderm (Schoenwolf *et al*, 2020). These cells line the ventricle creating a germinal zone, referred to as the ventricular zone (VZ). Around gestational week 7 (GW7), the NE cells acquire astroglia characteristics and transition into radial glia (RG) cells (Andrews *et al*, 2022; Götz & Huttner, 2005; Kriegstein *et al*, 2020). Around GW11 (Lui *et al*, 2011; Zarzor *et al*, 2023) certain RG cells transform, disconnecting their apical processes, becoming unipolar, and migrating away from the VZ (Hansen *et al*, 2010; LaMonica *et al*, 2013; Pollen *et al*, 2015). Accordingly, RGs subdivide into those that line the ventricle (ventricular RGs or vRGs, also known as apical RGs or aRGs), and those that migrate (outer or oRGs, also basal RGs or bRGs) (Andrews *et al*, 2022; Taverna *et al*, 2014). In this transformation oRGs inherit the basal process, which eventually leads to the entirety of vRGs disconnecting from the meninges and becoming truncated RG (tRG) cells midway into neurogenesis (around GW16.5) (LaMonica *et al*, 2013; Nowakowski *et al*, 2016). All neural stem cell (NSC) subtypes and their progenies are thus derived from the VZ. WWOX was shown to be most expressed in the VZ in neural organoids (Steinberg *et al*, 2021). This pattern, together with the neurodevelopmental implications of WWOX loss on patients, highlights the importance of WWOX in proper CNS development and neurogenesis. However, the specific molecular pathways affected by *WWOX* deficiency and how they contribute to disease pathogenesis is poorly understood.

In recent years, neural organoids have emerged as a revolutionary tool to model human brain development and disease in vitro (Arlotta, 2018). Neural organoids have been particularly useful in experimentally dissecting cellular heterogeneity in neurogenesis (Yang *et al*, 2022).

Our previous studies of WWOX deficient organoids revealed that multiple disease-relevant phenotypes like hyperexcitability and cellular composition abnormalities can be recovered and rescued by restoration of WWOX expression *in-vitro* (Steinberg *et al*, 2021). Macroscopic examination to these organoids with immunostaining and bulk RNA sequencing also suggested apparent changes in cellular populations and hypomyelination (Repudi *et al*, 2021b; Steinberg *et al*, 2021). However, what specific cell types and neurodevelopmental processes were affected the most by WWOX deficiency and how to ameliorate these alterations with clinically relevant therapeutic strategies remains an open challenge. To address these problems, in the present study we investigate the molecular, cellular, and functional consequences of WWOX mutations at cellular resolution by combining neural organoids derived from *WWOX*-deficient iPSCs and patient cells with single-cell transcriptomic profiling, reanalysis of human fetal data, and electrophysiological recordings. We discovered preferential expression of WWOX in RG cells, where its deficiency leads to impaired cell cycle progression, increased MYC expression, and reduction of neuronal generation. Patient-derived organoids showed a less severe phenotype than that seen in KO organoids, with generation of functionally impaired neurons that can be rescued with gene therapy.

## Results

### Generation of WWOX-deficient neural organoids

While WWOX mutations cause severe neurodevelopmental disorders, the molecular mechanisms underlying its role in human brain development remain poorly understood. To experimentally dissect the role of WWOX deficiency in human neurodevelopment, we generated unguided neural organoids, often referred to as cerebral organoids (COs) (Pașca *et al*, 2022), for wildtype (WT), WWOX knocked-out (WWOX-KO), and patient-derived (WOREE or SCAR12) conditions (**Fig. 1A**). We established a WT iPSC line from a healthy adult male donor (JH-iPS11) (**Fig. EV1A**). WWOX was knocked out in this line using CRISPR/Cas9 **(Fig. EV1B)**, and two subclones (JH-iPS11 WKO-1C and WKO-2C) were established. Sanger sequencing confirmed exon 1 indels introducing a premature stop codon, with both clones retaining normal karyotypes **(Fig. EVB-D)**. We used previously reported WOREE and SCAR12 iPSCs (23) for patient-derived lines and acquired new fibroblasts or peripheral blood mononuclear cells (PBMCs) isolated from skin biopsies and blood samples of additional WOREE patients, summarized in a table **(Fig. EV1E**). These additional individuals have complex heterozygous WWOX mutations. We reprogrammed these fibroblasts to iPSCs (**Fig. EV1F**) and termed the resulting lines LM-iPS and WCH S (**Fig. 1A**). WWOX-expression assessment with immunoblot (**Fig. EV1G**) revealed a decrease in protein level in the affected individuals compared to heterozygous and wildtype samples. This very low WWOX expression was consistent with all the lines previously established (Steinberg *et al*, 2021), despite the different phenotypes observed in the source patients and the derived organoids (Mallaret *et al*, 2014; Piard *et al*, 2019; Steinberg *et al*, 2021).

**Fig. 1.**
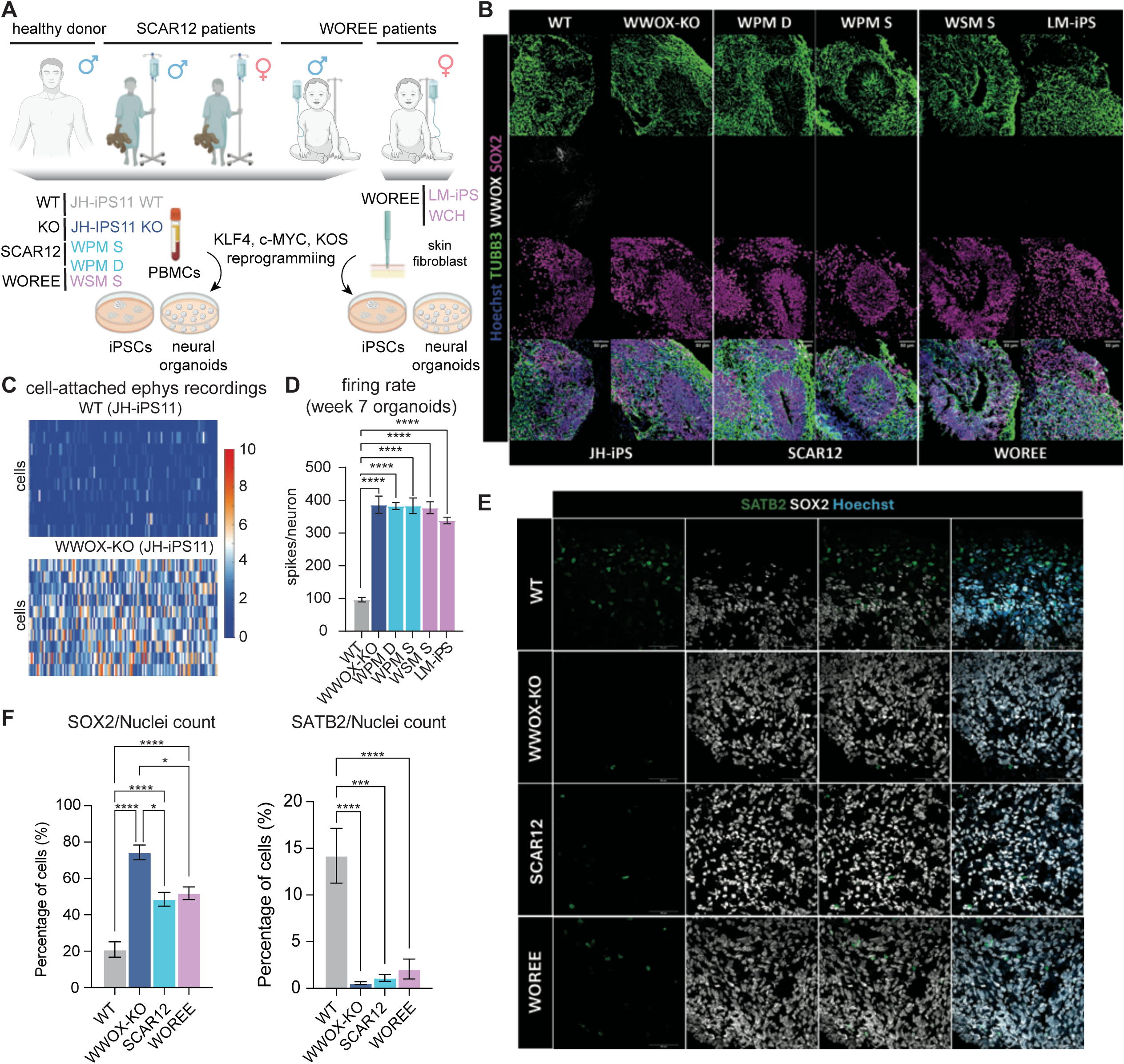
Generation of patient-derived neural organoids. **(A)** Schematic representation of the research approach employed in this study. This Fig. highlights the different stem cell lines and experimental groups used. KOS; KLF4-OCT3/4-SOX2 Sendai vector, iPSCs; induced pluripotent stem cells. **(B)** Representative Immunofluorescent staining of the unguided organoids (previously termed cerebral organoids; COs) used in this study, showing the pan-radial glia (RG) marker SOX2, the new-born neurons and RG marker β3-tubulin (TUBB3) and WWOX at week 7 of culture. (**C)** Raster plots demonstrating selected neuronal firing from WT and WWOX-KO week 7 organoids over 4 minutes of cell-attached recording. **(D)** Quantification of all the cell-attached recordings obtained from WT, WWOX-KO, WOREE and SCAR12 organoids, including those exemplified in (B). (WT: n= 30 neurons; WWOX-KO: n=30; WPM S: n=21; WPM D: n=21; WSM S: n=24; LM-iPS: n=14), collected from 1 batch. Statistical significance was determined by a one-way ANOVA test combined with Tukey’s correction for multiple comparisons. **(E)** Immunostaining for SATB2, an early expressed neuronal marker and superficial-layer neurons at week 7 COs, together with SOX2. **(F)** Quantification of the data shown in (E), n represents the number of organoids from two batches. (WT: n= 9, WWOX-KO: n=3; WOREE: n=15, SCAR12: n=7). See Table EV1 for details on organoid numbers, sections, and batches. Data represents biological and technical replicates and is presented as mean ± SEM. Statistical significance was determined using one-way ANOVA with Tukey’s multiple comparisons test. n.s (non-significant), *p ≤ 0.05, **p ≤ 0.01, ***p ≤ 0.001, ****p ≤ 0.0001.

Using this diverse iPSC cohort of WT, WWOX-KO, WOREE, and SCAR12 iPSCs (**Fig. 1A**) we established unguided organoids. Using immunofluorescence staining, we validated the presence of ventricular zone (VZ) and cortical plate (CP) structures at week 7 and assessed dorsoventral identity, with organoids showing predominantly dorsal characteristics (**Fig. 1B**), **(Fig. EVA-D)**. This analysis confirmed high WWOX expression in the VZ and reduced expression in all mutant lines. Cell-attached electrophysiological recordings of the organoids confirmed increased neuronal activity in WWOX-deficient organoids, consistent with the ability of *in-vitro* systems to recapitulate disease-relevant phenotypes (**Fig. 1C and Fig. EV1K**, **Fig. 1D**). Notably, SCAR12 patient-derived organoid recordings also showed hyperexcitability, a novel finding consistent with the epileptic seizures seen in patients (Mallaret *et al*, 2014) and with findings from a rodent model (Hussain *et al*, 2023).

Given the prominent expression of WWOX in the VZ, a critical neurogenic niche that orchestrates cortical architecture and cellular composition, we investigated alterations in early neurogenesis by analyzing the balance between RGs and nascent neurons during initial organoid development. At week 7, immunofluorescence analysis revealed a significant reduction in SATB2-positive cells concurrent with an expansion of the SOX2-positive population in WWOX-mutant organoids compared to controls (**Fig. 1E** and **Fig. 1F**), suggesting impaired transition from RGs to neurons. The severity of this neurogenic disruption demonstrated a genotype-dependent gradient, with WWOX knockout organoids exhibiting the most pronounced phenotype, followed by WOREE and SCAR12 variants, thus correlating with the overall phenotypic severity of human patients.

### scRNA-seq reveals impaired neurogenesis in WWOX-mutant organoids

To further examine the cellular and molecular composition of our organoids, we performed single-cell RNA sequencing (scRNA-seq) on organoids dissociated at week 16 **(Fig. EV1L-N)** using an established protocol (Velasco *et al*, 2019) (**Fig. 2A-B**, and **Fig. EV3A-B**). To identify the main cellular identities recapitulated by our experimental system, we grouped all cells jointly into 12 transcriptionally similar clusters following dimensional reduction and batch-correction analyses (Methods) (**Fig. EV3A**). By comparing these groups with cell type profiles from human fetal data (Polioudakis *et al*, 2019; Ramos *et al*, 2022; Trevino *et al*, 2021a; van Bruggen *et al*, 2022) and assessing the expression of known marker genes, we identified all major expected cell types, including RGs, neural progenitor cells (NP) (also referred to as intermediate progenitors), and neurons (Neu) (**Fig. 2A-D** and **Fig. EV3B-E**). Based on these comparisons, the total RGs population could be further interpreted in subsets of outer RGs (oRGs), cycling RGs (cRGs), and a mixed population of vRGs, neuroectodermal cells, and glial progenitor cells (RG) (**Fig. 1A** and **Fig. EV3C-D**).

**Fig. 2.**
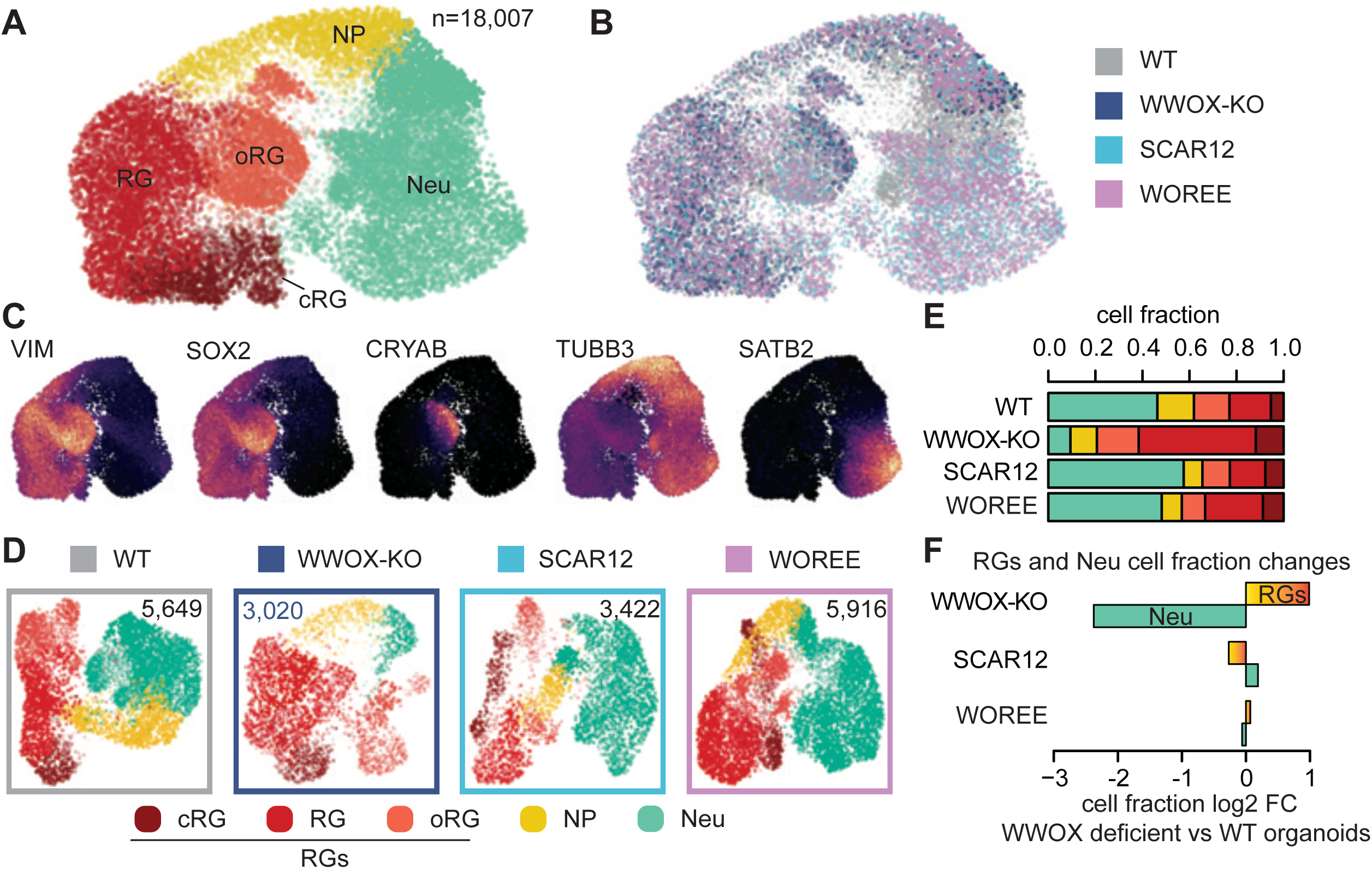
scRNA-seq reveals population changes in WWOX-deficient organoids. **(A)** Two-dimensional uniform manifold approximation and projection (UMAP) representation of cells colored by annotated cell types from all 4 cerebral organoid WWOX genotypes (total number of cells = 18,007). **(B)** The UMAP presented in (A), colored by condition. **(C)** Feature plots of renown marker genes for radial glia (VIM, SOX2), outer-radial glia (CRYAB) and neurons (TUBB3, SATB2). **(D)** Individual UMAP representation for each tested condition, with total cell numbers reported. **(E)** Stacked bar representing the cell fraction for the identified cell types for each condition. **(F)** Cell populations fraction changes in WWOX-mutated organoids, calculated relative to the WT fraction and presented as log2 fold change. RG = radial glia, cRG = cycling radial glia, oRG = outer radial glia, NP = neural progenitor, Neu = neurons.

Notably, WWOX-KO and WOREE organoids showed a prominent increase in the relative number of total recognized RG cells (RGs) (**Fig. 2D-F**), while SCAR12 organoids (modeling a relatively milder disease) exhibited an RG population similar to WT. This result was more pronounced in the WWOX-KO organoids, compared to the WOREE organoids. Concomitant to this cellular phenotype, WWOX-KO organoids also showed a dramatic decrease in identified neurons relative to WT organoids (**Fig. 2F**). Importantly, this was not the case in our patient-derived organoids. Although some changes could be observed in the cells interpreted as neural progenitors, oRGs or cRGs, due to the known heterogeneity of unguided organoids, we abstained from drawing conclusions at such finer resolution.

### Inverse relationship between MYC and WWOX expression through brain development and in neural organoids

Our current and previous data indicates that WWOX is expressed the highest in the ventricular-like zone (VZ) of human organoids at tissue-resolution (Steinberg *et al*, 2021) (**Fig. 1B**). Therefore, the impact of WWOX-deficiency on neurogenesis might primarily result from alterations to the cellular populations constituting this zone. To investigate developmental WWOX expression at higher resolution in this zone *in vitro* and *in vivo*, we complemented our organoids data with previously published transcriptomic data from human fetal samples at tissue-dissected (Miller *et al*, 2014), cell-sorted (Song *et al*, 2020), single-cell (Trevino *et al*, 2021a), and temporal resolutions (Cardoso-Moreira *et al*, 2019a). This multi-resolution analysis allowed us to further characterize WWOX expression patterns at cellular resolution across different developmental stages. Consistent with our observations in organoids, WWOX was preferentially expressed in RG cells *in vivo* at single-cell level at post-conceptional week 16 (**Fig. 3A-C**) and at bulk-level in SOX2^+^PAX6^+^ RG cells isolated from the germinal zone (GZ) of human cortical samples using fluorescence-activated cell sorting (FACS) (**Fig. 3D**), suggesting that the source of high WWOX expression in the fetal brain and in the VZ is the RGs and demonstrating that this pattern is not an artifact of the organoid system.

**Fig. 3.**
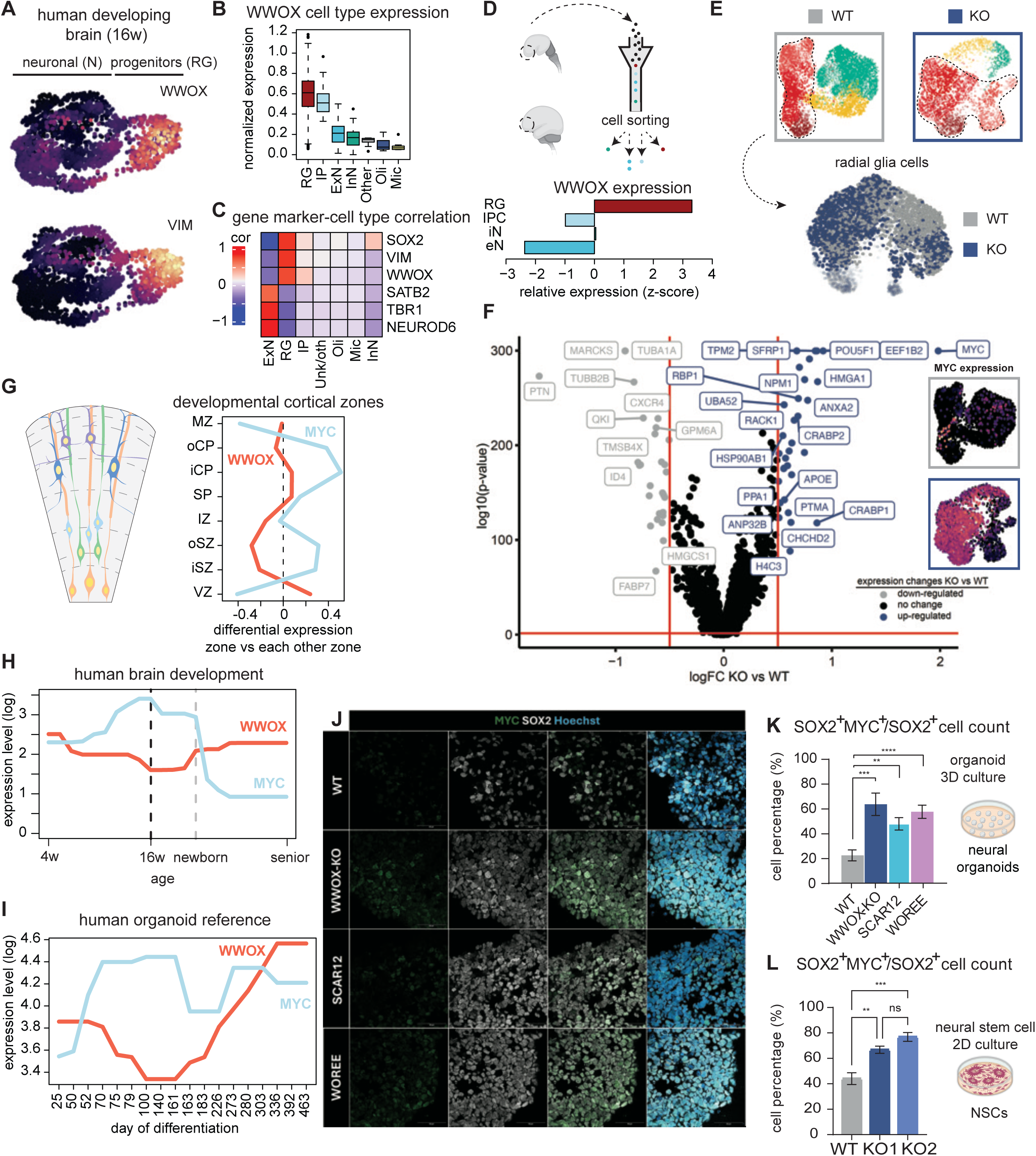
MYC negatively correlates with WWOX in primary brain samples and is activated in radial glia cells upon WWOX-KO. **(A)** WWOX and VIM expression plot in post conceptional week 16 human fetal single-cell data (10.1016/j.cell.2021.07.039). **(B)** Normalized WWOX expression across cell types of the single-cell fetal data in panel A. **(C)** Correlation of normalized marker expression to cell type membership in human fetal single-cell data. **(D)** WWOX normalized expression in human fetal cell sorted data (10.1038/s41586-020-2825-4). **(E)** Radial glia isolation from WT and WWOX-KO organoid (n=4,548). **(F)** Differential gene expression in radial glia from WWOX-KO and WT organoids. Upregulated genes in the WWOX-KO are shown in blue, downregulated genes in the WWOX-KO in grey, not significant genes are in black. Red bars show threshold of significance, -log10 p-value > 1.30 (p-value = 0.05) and log2 fold change of ± 0.5. On the left, MYC expression in WT and WWOX-KO organoids. **(G)** (Left) Schematic of developing cortical zones, (Right) pairwise signature of WWOX and MYC expression in cortical zones from w21 (10.1038/nature13185). **(H)** Normalized expression of WWOX and MYC in brain development from embryonic stages (w04) to adulthood (senior) (10.1038/s41586-019-1338-5). Top MYC expression around the peak of neurogenesis (w16). **(I)** Cortical organoid reference from Gordon et al. 2021 (10.1038/s41593-021-00802-y) showing the opposite trend between WWOX and MYC in time (day of differentiation). **(J)** Immunostaining of week 16 organoids showing high colocalization of MYC and SOX2, suggesting higher localization of MYC to the nucleus of WWOX-mutated (WWOX-KO, WOREE and SCAR12) organoids. **(K)** Quantification of the representative images shown in (J) (WT: n=7 organoids; WWOX-KO: n=5; SCAR12: n=8; WOREE: n=11). Statistical significance was determined using one-way ANOVA with Tukey’s multiple comparisons test. **(L)** 2D-cultured neural stem cells (NSCs) were immunostained for MYC, SOX2 and TUBB3 (see Fig. EV4C), and double-positive SOX2^+^MYC^+^ cells were quantified. The results are presented as a fraction of the total SOX2^+^ cells, data collected from two differentiation experiments. See Table EV1 for details on organoid numbers, sections, and batches Data represents biological and technical replicates and is presented as mean ± SEM. Statistical significance was determined using one-way ANOVA with Tukey’s multiple comparisons test. n.s (non-significant), *p ≤ 0.05, **p ≤ 0.01, ***p ≤ 0.001, ****p ≤ 0.0001. w = post conceptional week.

Because the most striking changes in the RGs population in our organoids were observed in the KO genotype, we focused on the comparison between RGs from the WT and WWOX-KO organoids to uncover potential molecular alterations mediating WWOX-deficient phenotypes (**Fig. 3E**). Differential gene expression analysis of WWOX-KO vs WT RG cells identified the proto-oncogene MYC (also known as c-Myc) as the top significantly upregulated gene in WWOX-deficient RGs (**Fig. 3F**). This result is consistent with a previous observation of MYC overexpression upon WWOX ablation in osteosarcoma (Akkawi *et al*, 2024). However, this is, to our knowledge, the first time that an inverse relationship between WWOX and MYC has been observed in non-neoplastic tissue. To investigate whether this inverse association is recapitulated *in vivo* and reproducible *in-vitro*, we analyzed MYC and WWOX expression patterns across cortical zones of the fetal human brain at mid-gestation (Miller *et al*, 2014) (**Fig. 3G**) and across developmental time in human forebrain samples (Cardoso-Moreira *et al*, 2019a) (**Fig. 3H**). In both cases a clear inverse correlation emerged in which increasing WWOX expression coincides with decreasing MYC expression and vice versa. This pattern was also reproduced in an independent transcriptomic dataset of human neural organoid maturation (Gordon *et al*, 2021) (**Fig. 3I**).

To validate these transcriptional patterns at the protein level, we immunostained cerebral organoids (weeks 10-16) and found a significant increase in double positive cells for SOX2 and MYC (**Fig. 3J** and **Fig. 3K**). This increase in MYC expression was observed also in the RGs from SCAR12 and WOREE organoids (**Fig. 3J-K** and **Fig. EV4A-D**). To further validate this observation, we modeled WWOX-deficient NE and RG cells in our previously described WWOX-KO WiBR3 human embryonic stem cell (hESC) system (Steinberg *et al*, 2021) using our unguided organoids protocol and assessed dorsal identities **(Fig. EV2E-F)**. Through immunofluorescence staining, we observed a marked increase in MYC-expressing SOX2+ cell populations **(Fig. EV2G-H)** at week 15 *in-vitro*, corroborated by a similar imbalance in RG and neuronal populations **(Fig. EV2I-J)**. Additionally, to substantiate our findings in an independent *in vitro* system, we modeled WWOX-deficient NE and RG cells in 2D by differentiating neural stem cells (NSCs) from the latter described hESCs-WWOX-KO system using an established protocol (Cohen-Carmon *et al*, 2020) (**Fig. EV5**). These cells expressed high levels of SOX2 and β3-tubulin, and WWOX in the WT NSCs (**Fig. EV5A**), confirming their identity. Immunoblot and immunostaining of these cells confirmed high MYC expression upon WWOX-KO (**Fig. 3L** and **Fig. EV5B-D**).

Altogether, our data uncovered a reproducible inverse relationship between WWOX and MYC, extending this association from cancer to neurodevelopment. This relationship seems to be robust across brain development, as it is evident not only in our in vitro models, but also in reference fetal and organoids datasets, and in both WOREE and SCAR12 organoids.

### Cell-cycle and potency abnormalities in WWOX-deficient MYC-overexpressing RGs

The MYC gene is a well-known regulator of cell proliferation and differentiation, and its dysregulation has been implicated in cell cycle control disruption in various contexts (Ahmadi *et al*, 2021; Felsher *et al*, 2000; García-Gutiérrez *et al*, 2019; Schuhmacher & Eick, 2013). To investigate potential cell cycle alterations in WWOX-mutated RGs, we estimated deviations in cell cycle progression between WT and WWOX-KO organoids by inferring cell cycle state from single-cell expression using a previously established computational tool *tricycle* (Zheng *et al*, 2022). Inference is based on estimating per cell a continuous cell-cycle pseudotime that represents progression through the cell-cycle phases (Methods). This analysis revealed an increase in the fraction of RGs with an estimated S (0.5π - 1π pseudotime) or G2M (1π - 0.5π) cell cycle state in WWOX-KO organoids compared to WT, suggesting cell cycle alterations in RG under WWOX deficiency (**Fig. 4A-C**). This is consistent with the known MYC effects on cell cycle, including acceleration of the transit through G1/S and prolongation of the G2/M phases (Deb-Basu *et al*, 2006; Felsher & Bishop, 1999; Schuhmacher & Eick, 2013). Radial glial cells estimated to be in G2/M or S phases also expressed higher levels of MYC compared to G1/G0 and M phase cells in WWOX-KO organoids but not in WT (**Fig. 4D**). To experimentally validate these findings, we co-stained week-16 organoids with phospho-histone H3 (pH3), a marker for cells in late G2-M phases, and SOX2, which confirmed an increased fraction of SOX2⁺ committed to mitosis in WWOX-KO and WOREE organoids, with a milder increase in SCAR12, consistent with a genotype-dependent gradient **(Fig. EV6A-B)**.

**Fig. 4.**
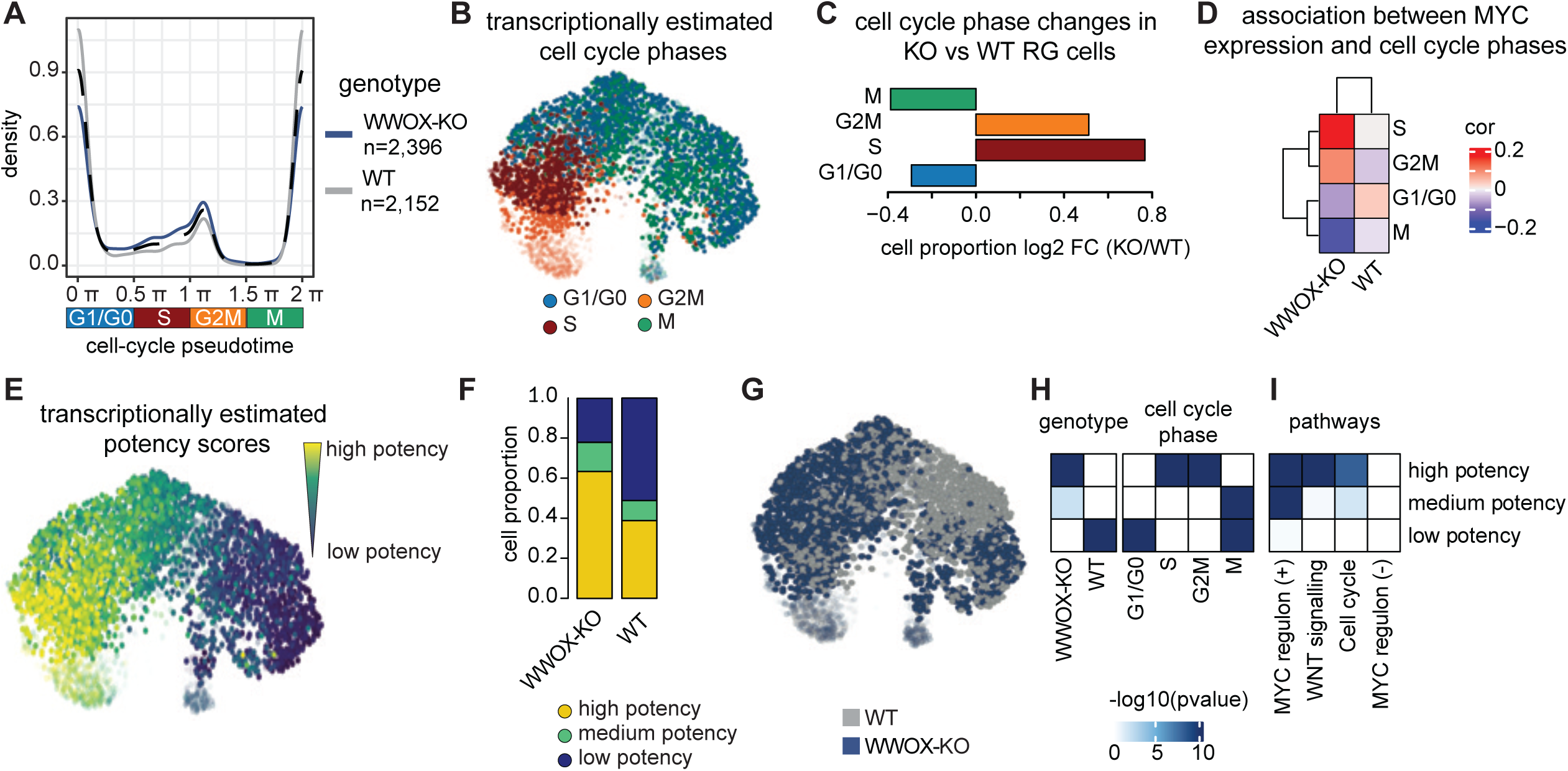
WWOX-KO leads to G2/M and S arrest and cellular metabolism alterations. **(A)** Cell cycle phase inference in radial glia cells. Time points define a continuous scale of cell cycle states, that can be discretized as 0π - 0.5π as G1/G0 phase, 0.5π - 1π as S phase, 1π - 1.5π as G2M phase and 1.5π - 2π as M phase. The lines define how many radial glia cells are in a specific phase of the cell cycle (n=2,396 for WWOX-KO, n=2,152 for WT). Cell landscape visualization of assigned cell cycle phase. **(C)** Log fold change of WWOX-KO on WT cell proportions across cell cycle phases, colored as in (B). **(D)** MYC normalized expression correlation to cell cycle phases showing high values for S and G2M phase in WWOX-KO organoids, and G1/G0 for WT. **(E)** UMAP colored by inferred potency scores. **(F)** Stacked bars quantifying potency states proportions in WWOX-KO and WT conditions. **(G)** Cell projection of organoid conditions. **(H)** Overrepresentation analysis of cells in each condition and cell cycle phase across potency states. High potency cells are overrepresented in WWOX-KO condition and in S - G2M phases, while Low potency in WT and G1/G0 - M phases. Medium potency cells are overrepresented in M phase, in WWOX-KO condition. **(I)** Enrichment analysis of pathways in transcriptional signatures of potency states. High potency cells are enriched in Wnt and cell cycle pathways, with targets regulated positively by MYC. Medium potency cells show similarly enrichment for MYC regulon and Wnt. Low potency cells show no specificity. -log10(p-value) were capped to have a maximum score of 10.

Next, since MYC is known to maintain the potency of PSCs (Chappell & Dalton, 2013) and was shown to block the differentiation of neural progenitors to postmitotic neurons in the mouse embryo (Wang *et al*, 2020), we hypothesized that it might have a similar effect in RG cells following loss of WWOX. To look for transcriptomic evidence of this, we classified RG cells in three groups from low to high commitment level based on “potency scores” estimated from single-cell expression using a previously established machine learning algorithm *CytoTrace2* (Kang *et al*, 2024). Most WWOX-KO RGs were estimated to have a high potency state, while WT RGs showed a low-potency state (**Fig. 4E-G** and **Fig. EV6C**), possibly supporting a notion of MYC-associated impaired differentiation. Pathway enrichment analysis of the cells in different pluripotency states revealed overrepresentation of genes involved in cell cycle, DNA replication, and cellular senescence in high potency cells (**Fig. EV6D**). This was further supported by high levels of p53 in WWOX-KO NSCs (**Fig. EV6E**). To test for associations between genotype, potency state, and cell cycle phase we performed statistical overrepresentation analysis. We found that high-potency RGs are overrepresented within WWOX-KO cells and cells in S - G2/M phases, while low-potency RGs are enriched within WT cells and cells in G1/G0 - M phases (**Fig. 4H**). To find additional evidence of MYC-mediated differentiation impairment of RG cells, we compiled from an integrative database target-genes with evidence of positive or negative regulation by MYC (i.e., regulons) and tested their association with RG potency states (Methods). WWOX-KO high-potency cells preferentially express target genes positively regulated by MYC, further suggesting a mediating role of MYC in impaired RG differentiation (**Fig. 4I**). Negatively regulated targets did not show an association. Further supporting this MYC-dependent mechanism, the Wnt pathway, which can induce expression of MYC and vice versa (He *et al*, 1998; Hao *et al*, 2019), has previously been shown to be activated in WWOX-mutated organoids (Steinberg *et al*, 2021). In line with these findings, our current enrichment analysis of Wnt signaling genes also showed evidence of association of this pathway with high-potency RGs (**Fig. 4I**). Medium-potency cells showed similar enrichment for MYC regulons and Wnt signaling, while low-potency cells did not show an association.

To validate these findings, we assessed Wnt pathway activity in WWOX-KO, WOREE, SCAR12, and rescued WWOX-KO organoids **(Fig. EV6F).** Western blot analysis revealed elevated levels of TCF4 (TCF7L2), a canonical Wnt effector that partners with β-catenin to drive transcription of Wnt target genes. This increase aligns with prior reports of TCF4 upregulation upon Wnt activation in human NPCs (Hennig *et al*, 2017). Thus, aberrant TCF4 induction in WWOX-deficient organoids is consistent with enhanced Wnt signaling observed in our WWOX-KO RGs.

To further substantiate MYC-mediated RG dysregulation, we leveraged our scRNA-seq dataset and explored convergent molecular networks shared by MYC and WWOX and perturbed under WWOX deficiency. We considered the intersection of the regulon and interactome of MYC, the interactome of WWOX, and differentially expressed genes to prioritize molecular candidates **(Fig. EV7A)**. We identified 12 candidates, including genes associated with cell cycle regulation (E2F1, TFAP2C, YEATS4) (Wu *et al*, 2001; Wong *et al*, 2012; Xian *et al*, 2023), stemness (NME2, CSDE1) (Qi *et al*, 2021; Ju Lee *et al*, 2017), RNA metabolism and ribosome biogenesis (DKC1) (Garus & Autexier, 2021), NPC proliferation (PQBP1) (Liu *et al*, 2024), and neuronal heterotopia (PAICS) (Agarwal, 2020; Yamada, 2020). Because reference interactomes and regulons do not consider cell type specificity, we assessed preferential expression in RG, IPC, or early neuronal (eN) cells using human fetal RNA-seq data (**Fig. EV7B**). Five of the candidates (*E2F1, YEATS4, PAICS, TFAP2C,* and *DKC1*) showed preferential expression in RG or IPC cells, further prioritizing their potential involvement in MYC-mediated RG dysregulation.

To provide additional independent evidence for a potential role of MYC involvement in the reduced differentiation potential of WWOX deficient neural progenitors, we performed MYC chromatin immunoprecipitation sequencing (ChIP-seq) in hESC-derived WWOX-KO and WT NSCs **(Fig. EV7D-F).** We found that MYC binding is enriched in the promoters of genes preferentially expressed by early postmitotic neurons in WT NCSs, including *BACH2*, *CACNA1E, LRRC7, CNTN3,* and *SOX11* **(Fig. EV7E-F).** Notably, this enrichment is lost in WWOX-KO NSCs, suggesting a reduced priming for neuronal differentiation. In contrast, MYC binding is stronger in KO versus WT in the candidate genes *DKC1* involved in DNA replication, and *CSDE1* involved in neuronal differentiation and predicted to be repressed by MYC **(Fig. EV7A).**

Together, these results suggest that WWOX deficiency sustains MYC–Wnt signaling, driving RG cell-cycle acceleration and high potency thereby impairing their neuronal differentiation potential.

### Early gene expression signature in WOREE- and SCAR12-derived neurons

After having established a role for WWOX-deficiency in neurogenesis through alterations in early progenitors, we next sought to analyze potential consequences of additional alterations in the generated neuronal populations in our organoids. Due to the low number of neurons generated in WWOX-KO organoids (**Fig. 2E**), the analysis focused on WOREE- and SCAR12-derived neurons (**Fig. 5A**). We isolated, combined, and re-clustered the cells transcriptionally identified as NP or neurons in the initial analysis of the scRNA-seq data (**Fig. 2A** and **Fig. EV8A**). We included NP cells to consider potential early alterations on identity specification. This resulted in 11 neuronal (ExN) clusters which were then compared to reference developmental neuronal cell types for interpretation (Wang *et al*, 2024) (**Fig. EV8B** We defined 7 subgroups of neurons (**Fig. 5B** and **Fig. EV8C**) with signatures corresponding to deep-layer excitatory neurons, as well as newborn and immature neurons. Based on transcriptional similarity with signatures of neuronal subpopulations sequentially observed in fetal neurogenesis, we further classified these 7 subgroups into two general groups – neurons with a gene signature resembling early or late developmental stages during neurogenesis. Cells interpreted as having an early expression signature correlate with *in-vivo* signatures of NP, immature, and newborn neurons, while those with late signatures primarily correlate with L5 and L6 excitatory neurons. A subset of late neurons (mixed) correlated with signatures of both excitatory and inhibitory neurons (**Fig. EV8B**). Early cells express regulators of the initial specification of neocortical cell fate (e.g., PAX6, EMX2, and FOXG1) and the immature marker DCX, and are overrepresented with pathways involved in biosynthesis, neuron differentiation, Wnt signaling, and cell proliferation. In contrast, late cells express regulators of the postmitotic specification of deep-layer neurons (e.g., FEZF2, and BCL11B/CTIP2), the calcium sensor SYT1, voltage-gated sodium and potassium channel subunits (e.g., SCN2A, SCN1A, KCNA3, KCNA4), and are overrepresented with pathways involved in carbohydrate metabolism, neurotransmitter secretion, synapses, and membrane depolarization (**Fig. 5C** and **Fig. 5D**). Although neurons of both groups appeared in all three organoids analyzed, so-called “early cells” were enriched in the WOREE- and SCAR12-derived organoids (**Fig. 5F** and **Fig. 5G**). Conversely, late neurons were enriched in the WT organoids. This observation was further highlighted by increased expression of neurogenesis-associated genes, such as NEUROD6 and NFIB, in both WOREE- and SCAR12-derived neurons as compared to WT (**Fig. EV9A and Fig. EV9B**). One possible interpretation of these results is that the observed differential maturity of transcriptional signatures might be a consequence of the earlier alterations in RG cells. Notably, only two genes were found to be differentially expressed between SCAR12 and WOREE neurons, suggesting that considerable differences in the manifestations of the diseases do not arise directly from the neuronal (**Fig. EV9C**).

**Fig. 5.**
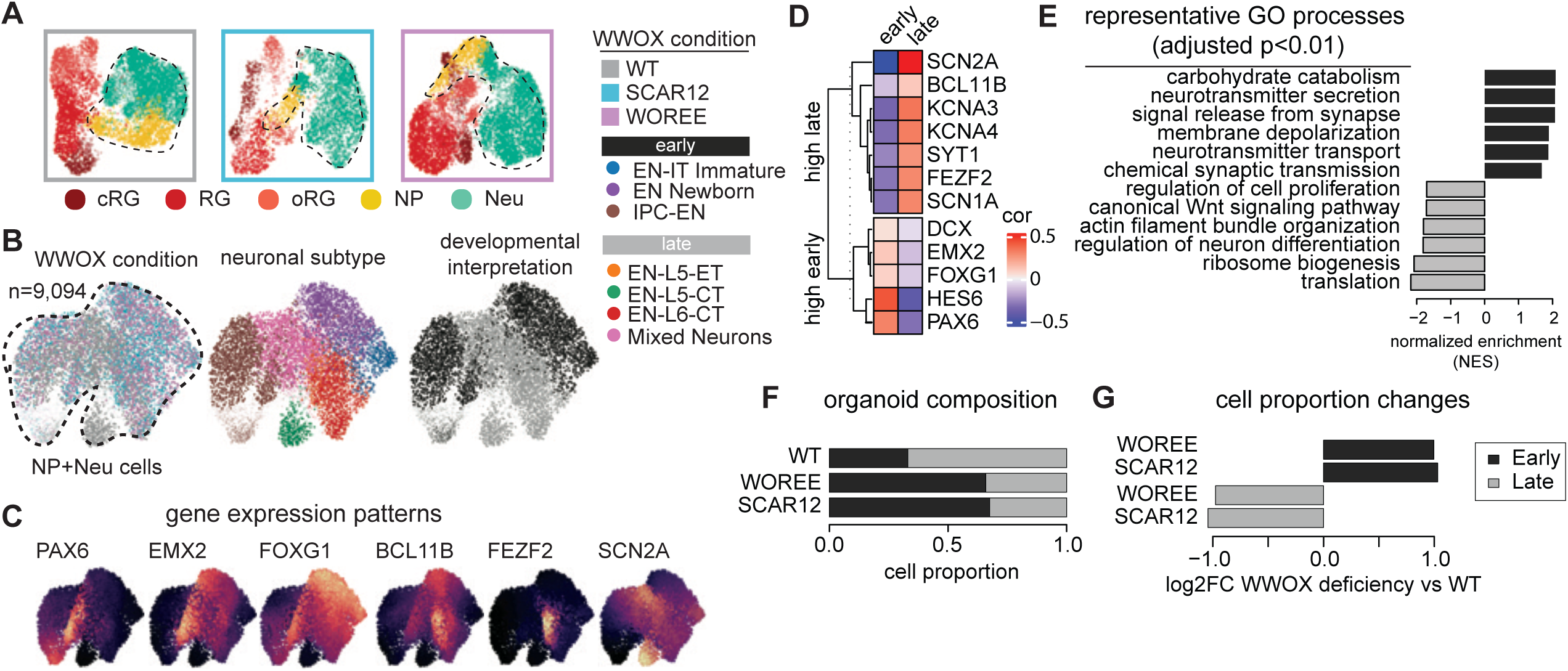
Neuronal population analysis in WT and patient-derived lines. **(A)** Isolation of neural progenitors (in yellow) and neurons (in green) from WT, SCAR12 and WOREE conditions. **(B)** UMAP visualization of neural progenitors (NP) neuronal (Neu) cells colored by condition (left), assigned neuronal subtype (middle) and developmental categories (right). Identified categories are divided in early stages of neuronal development (dark grey) and late stages (light grey). **(C)** Feature plots of renown marker genes for progenitor cells (PAX6, EMX1, FOXG1) and neurons (BCL11B, FEZF2, SCN2A). **(D)** Marker gene normalized expression correlation to developmental categories (“Late” and “Early” neurons). **(E)** Representative Gene Ontology (GO) terms enriched in “late” cells compared to “early” cells, plotted as normalized enrichment score (NES). **(F)** Quantification of log fold change of early and late stages of neuronal subtypes across conditions. **(G)** Comparison of neuronal subtype proportions in patient lines compared to WT early and late stages.

### Neuron-specific Gene Therapy Rescues Neuronal Hyperexcitability

We have previously shown that neuronal restoration of WWOX using an adeno-associated virus serotype 9 (AAV9), driven by Synapsin I promoter (*AAV9-SynapsinI-WWOX-2A-EGFP-WPRE*, referred to here as AAV9-WWOX) is able to rescue phenotypes seen in *Wwox*-null mice, including seizures, early lethality, and infertility (Repudi *et al*, 2021a). Given this remarkable success in *Wwox*-null mice, we set out to determine whether the molecular and electrophysiological phenotypes observed in the organoids could be rescued by neuronal AAV9-WWOX infection, irrespective of prior RG defects. To this end, we infected week 10 organoids with either AAV9-WWOX (see methods) or AAV9-EGFP (*AAV9-SynapsinI-EGFP-WPRE*), followed by 6 weeks of culture We confirmed the success of infection in week 16 organoids by immunostaining and quantification of fraction of NEUN+WWOX+ from all NEUN+cells (**Fig. 6A** and **Fig. EV10A-B**), and through western blot analysis of total protein content **(Fig. EV1L-N).** Neuronal activity was then measured through Ca^2+^ imaging, revealing that the firing rate of WOREE organoids was approximately 3-fold higher than that of SCAR12 organoids (**Fig. 6B** and **EV11**), in contrast to what we observed in an earlier time point, where both organoids showed similar neuronal hyperactivity (**Fig. 1B** and **Fig. 1C**). In both cases neuronal firing was significantly higher compared to the WT, with differences observed in event frequency, firing amplitude, and the proportion of active regions of interest (ROIs) in each organoid. Remarkably, neuron-specific restoration of WWOX was sufficient to reduce hyperexcitability to a level similar to the WT organoids (**Fig. 6B**).

**Fig. 6.**
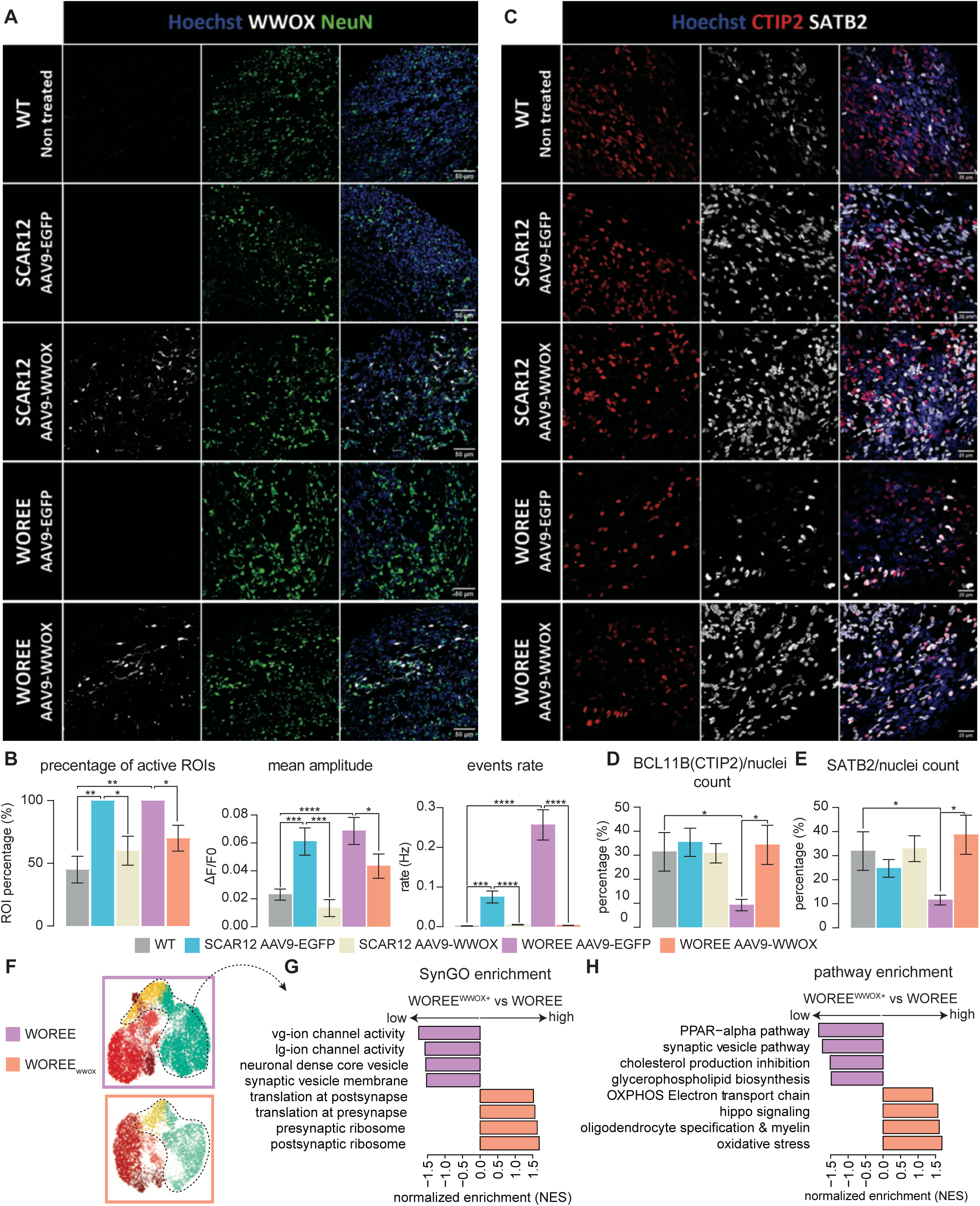
Neuronal restoration of WWOX using an AAV9 vector rescued hyperexcitability and neuronal abnormalities in WWOX-deficient organoids. Week 10 COs were infected with either AAV9-SynI-EGFP-WPRE (AAV9-EGFP) or AAV9-SynI-WWOX-2A-EGFP-WPRE (AAV9-WWOX) and grown for 6 weeks. **(A)** Validation of the successful infection of week 16 COs with AAV9-WWOX (WT: n=4 organoids, SCAR12 AAV9-EGFP: n=8, SCAR12 AAV9-WWOX: n=8, WOREE AAV9-EGFP: n=7, WOREE AAV9-WWOX: n=7). **(B)** Neuronal activity visualized using Ca2+ recording in week 16 COs, comparing AAV9-EGFP and AAV9-WWOX infection. (WT: n=3 organoids, SCAR12 AAV9-EGFP: n=6, SCAR12 AAV9-WWOX: n=8, WOREE AAV9-EGFP: n=6, WOREE AAV9-WWOX: n=6). Statistical significance was determined using multiple t-tests, correcting for multiple comparisons by controlling the False Discovery Rate (FDR) using the Benjamini–Hochberg procedure. (Left) Percentage of active regions of interest (ROIs) within the imaged organoids. (Middle) Mean event amplitude, reflecting number of spiking neurons. (Right) 3Calcium events rate (WT: n=20 ROI, SCAR12 AAV9-EGFP: n=30, SCAR12 AAV9-WWOX: n=30, WOREE AAV9-EGFP: n=40, WOREE AAV9-WWOX: n=40). **(C)** The expression of cortical layers marks in week 16 organoids following the infection with either AAV9-EGFP or AAV9-WWOX, as shown by immunofluorescence stainings. **(D)** Quantification of CTIP2+ nuclei, divided by the total number of nuclei in each field counted. (WT: n=4 organoids, SCAR12 AAV9-EGFP: n=8, SCAR12 AAV9-WWOX: n=8, WOREE AAV9-EGFP: n=7, WOREE AAV9-WWOX: n=7). Statistical significance was determined by a one-way ANOVA test, correcting for multiple comparisons by controlling the FDR using the Benjamini–Hochberg procedure. **(E)** Quantification of SATB2+ nuclei, divided by the total number of nuclei in each field counted. (WT: n=4 organoids, SCAR12 AAV9-EGFP: n=8, SCAR12 AAV9-WWOX: n=8, WOREE AAV9-EGFP: n=7, WOREE AAV9-WWOX: n=7). Statistical significance was determined by a one-way ANOVA test, correcting for multiple comparisons by controlling the FDR using the Benjamini–Hochberg procedure. **(F)** Analysis of gene sets enrichment in neuronal progenitors and neurons in WOREE and WOREE-WWOX organoids. Bars represent normalized enrichment scores that are high in the treated organoid (salmon), related to synaptic compartment and translation, and low (lilac), associated to ion channel activity, both voltage-gated (vg) and ligand-gated(lg). **(G)** Pathway enrichment showing an increase OXPHOS and oxidative stress terms in WOREE-WWOX compared to WOREE, while downregulating pathways related to lipid metabolism. See Table EV1 for details on organoid numbers, sections, and batches. Data represents biological and technical replicates, collected from one batch and is presented as mean ± SEM. Statistical significance was determined using one-way ANOVA with Tukey’s multiple comparisons test. n.s (non-significant), * p ≤ 0.05, ** p ≤ 0.01, *** p ≤ 0.001, **** p ≤ 0.0001.

Previously, we reported reduced SATB2 and CTIP2 expression (both by immunofluorescence and mRNA levels) in WWOX-KO and WOREE organoids (Steinberg *et al*, 2021). Therefore, we assessed if AAV9-WWOX infection could increase the expression of these markers. Immunostaining and qPCR analysis revealed again lower CTIP2 and SATB2 expression in WOREE organoids compared to the WT (**Fig. 6C-E** and **Fig. EV12**). WWOX restoration increased the expression of both markers in WOREE organoids but not in SCAR12, which showed CTIP2 and SATB2 expression levels similar to WT. Next, to assess whether transcriptomic changes reflect our electrophysiological observations, we compared expression levels of neuronal cells from AAV9-WWOX treated WOREE organoids vs non-treated (**Fig. 6F-H**). AAV9-treated organoids showed changes in expression of mediators of neuronal activity, including reduced voltage-gated and ligand-gated ion channel expression and increased synaptic translation, together with an increase in genes involved in pathways suggestive of neuronal maturation, such as oxidative phosphorylation (primary energy source for resting neuronal terminals and upregulated during cortical neuron differentiation) (Agostini *et al*, 2016; Zheng *et al*, 2016; Wei *et al*, 2023), promotion of oligodendrocyte differentiation and myelination, and synaptic function (**Fig. 6G** and **Fig. 6H**). Importantly, the infection with AAV9-WWOX of WOREE organoids did not affect the RGs, as appreciated by the absence of differentially expressed genes and cell-cycle abnormalities (**Fig. EV13**), potentially pointing to the safety profile of this approach.

## Discussion

In this study, we report insights into the cellular and molecular processes underlying WWOX-related neurological disorders. WOREE and SCAR12 syndromes are devastating diseases, yet due to their rarity, are rendered orphan. Our findings from the combination of iPSC-based models and omics analysis indicate that WWOX-deficient neural organoids exhibited reduced neurogenesis and an increased number of RGs, particularly in WWOX-deficient organoids. These RGs exhibited increased expression of MYC, verified in 2D-cultured NSCs, which could significantly influence brain development. Another major finding in our study was cell cycle alterations in WWOX-deficient RG cells, with cells accumulating in the G2/M and S phases. In addition, we provide evidence that neuronal WWOX restoration using an AAV9-based gene therapy could rescue hyperexcitability of patient-derived organoids.

Given the fundamental role of WWOX in neurodevelopment, we posited that its complete loss may impair neurogenesis. To directly address this, we established cerebral organoids devoid of WWOX expression using CRISPR-edited and patient-derived iPSCs. This model was shown previously to effectively study the development and disease of NSCs within a human context (Di Lullo & Kriegstein, 2017; Pollen *et al*, 2019; Eichmüller & Knoblich, 2022). Given the high expression of WWOX in vRG cells, we anticipated that its loss would influence neuronal differentiation. Our data demonstrates that WWOX deficiency leads to disrupted neurogenesis in brain organoids, with clear and statistically significant alterations in progenitor populations. Consistent with this, analyses of WiBR3 hESC-derived WT and KO organoids at week 15 (Steinberg *et al*, 2021), ) (Fig. EV2E–J) revealed significant increases in SOX2⁺ populations relative to wild type, albeit with inter-clonal heterogeneity, and an overall reduction in neurons during maturation, underscoring the reproducibility of this effect. Importantly, the incorporation of single-cell transcriptomics in the current study enables the identification of heterogeneous RG cells beyond SOX2-defined populations, providing a more detailed and comprehensive view of how WWOX loss perturbs corticogenesis. Indeed, unbiased scRNA-seq revealed striking alterations in cellular populations, characterized by a marked increase in RG cells, correlated with MYC upregulation and a decrease in mature neurons, further emphasizing the pleiotropic functions of the *WWOX* gene (Abu-Remaileh *et al*, 2015; Banne *et al*, 2021). Our conclusions are based on the establishment and analysis of multiple independent models, including the previously reported hESC-WWOX KO, WOREE and SCAR12 lines (23), newly generated wild-type and WWOX-KO JH-iPS11 lines, and compound heterozygous WOREE patient-derived lines, LM-iPS (6) and WCH S. Together, these complementary models enabled consistent detection of disease-specific phenotypes across WOREE and SCAR12, while minimizing patient-specific bias and enhancing translational relevance.

Our findings implicate MYC in the observed phenotype, but whether WWOX regulates MYC directly or indirectly remains unresolved. Notably, previous WWOX pull-down experiments and mass spectrometry analyses did not identify MYC as a direct binding partner (Abu-Odeh *et al*, 2014; Khawaled *et al*, 2020). However, WWOX is known to affect several MYC modulators, such as Dvl and GSK-3β (through Wnt pathway) (Wang *et al*, 2012; Abu-Odeh *et al*, 2014; Bouteille *et al*, 2009; Sze *et al*, 2004), p53 (Sachdeva *et al*, 2009; Chang *et al*, 2005, 53; Del Mare *et al*, 2016; Abdeen *et al*, 2018) and HIF1α (Kaidi *et al*, 2007; Abu-Remaileh & Aqeilan, 2014). The implications of this relationship could significantly affect neuronal differentiation.

Aberrant Wnt activation is evident in WWOX-deficient organoids, with scRNA-seq localizing this dysregulation to RGs that show impaired proliferation and differentiation. Wnt signaling regulates RG fate, with activation blocking cortical differentiation and inactivation promoting terminal neurogenesis (He *et al*, 1998; Harrison-Uy & Pleasure, 2012; Gan *et al*, 2014; Machon *et al*, 2007), WWOX normally modulates this pathway through GSK-3β inhibition, and its loss phenocopies GSK-3β deletion, leading to RG hyperproliferation and enhanced differentiation (Wang *et al*, 2012; Kim *et al*, 2009). Mechanistically, we observed induction of TCF4 (Fig. EV6F), a canonical Wnt effector, upregulated in NPCs upon Wnt activation (Hennig *et al*, 2017). TCF4 enhances MYC activity in neuroblastoma (Aljouda *et al*, 2025) and regulates MYC in NPCs via super-enhancer binding (Quevedo *et al*, 2019), while loss-of-function mutations cause Pitt– Hopkins syndrome (Papes *et al*, 2022). Thus, TCF4 induction in WWOX-deficient organoids not only reflects Wnt activation but may also amplify MYC-driven programs, sustaining RG expansion and disrupting neuronal differentiation,

Given these insights into MYC dysregulation in WWOX-deficient RGs, it is noteworthy that WWOX was initially identified as a putative tumor suppressor in breast cancer, with the potential to influence cell growth (Bednarek *et al*, 2000). Therefore, our findings align closely with the expected functions of WWOX; however, our observations in a non-tumor context were previously unforeseen. Our observed increase in Myc expression may influence cell cycle progression, a phenotype prominently observed in WWOX-mutant organoids. The implications of this relationship could significantly affect neuronal differentiation. Previous research using mouse embryonic cultures has demonstrated that elongation of the G1 phase during neurogenesis promotes neural progenitor differentiation, while a reduction in G1 duration inhibits neurogenesis and favors progenitor expansion (Di Lullo & Kriegstein, 2017; Pollen *et al*, 2019). The activation of MYC can also result in genomic instability (Meyer & Penn, 2008; Dhanasekaran *et al*, 2022), correlating with the increased DNA damage upon WWOX deficiency (Steinberg *et al*, 2021). Elevated p53 levels in this context could suggest a compensatory response. While WWOX-deficient organoids demonstrated a reduction in differentiated neurons, this effect was less pronounced in patient-derived organoids, indicating a potentially detrimental process in which damaged RGs differentiate into neurons. Both MYC and p53 are known to impact cellular metabolism (Goetzman & Prochownik, 2018; Liu *et al*, 2019), potentially affecting neuron maintenance and neurotransmitter production (Hertz, 2013; Díaz-García *et al*, 2017), although these outcomes were not studied here.

WWOX-deficient RGs fail to properly silence pluripotency programs, as evidenced by persistent upregulation of MYC and TFAP2C during the hiPSC-to-NSC transition (Luciani *et al*, 2024; Pastor *et al*, 2018) **(Fig. EV7A-C).** This maintenance of stemness factors likely drives RG expansion at the expense of neurogenic commitment, a phenotype validated across multiple WWOX models, including patient-derived organoids. Importantly, MYC dysregulation extended to ribosome biogenesis, with strong induction of the pseudouridine synthase DKC1, a direct MYC target that regulates rRNA processing and translation efficiency (Garus & Autexier, 2021; O’Brien, 2016; Miao, 2019). This aligns with our observation of “early neuron” states enriched for ribosome biogenesis in WOREE and SCAR12, implicating disrupted protein synthesis in compromised neuronal output.

MYC dysregulation in WWOX deficiency also impacts neuronal metabolism. During embryogenesis, MYC normally declines to enable the shift from glycolysis to oxidative phosphorylation, a key transition for neuroepithelial maturation (Abu-Remaileh & Aqeilan, 2014). In proliferating progenitors, MYC drives glycolytic gene expression (e.g., HK2, LDHA), whereas in neurons its downregulation allows transcription factors such as PGC-1α and ERRγ to promote oxidative phosphorylation (Wang *et al*, 2020). The persistence of MYC in WWOX-deficient RGs may therefore sustain a glycolytic program and interfere with this metabolic reprogramming, further limiting neuronal differentiation and maturation.

Beyond differentiation, MYC–WWOX interactions converge on pathways governing neuronal migration. Consistent with prior histopathological and transcriptomic studies of a WOREE fetus by Iacomino et al. (2020) (Iacomino *et al*, 2020), which reported disrupted cortical layering, including mislocalized external granular neurons, disorganized radial glia, and reduced vascularization, our analysis identified MYC-dependent upregulation of PAICS, an enzyme in de novo purine synthesis linked to progenitor maintenance and ventricular heterotopia (Agarwal, 2020; Yamada, 2020; Mizukoshi *et al*; Chan *et al*). Together, these findings highlight MYC dysregulation as a central driver of WWOX-deficient phenotypes, integrating stemness, ribosome biogenesis, metabolism, and migration into a unified framework of impaired corticogenesis in WOREE and SCAR12.

These results reveal a broad phenotypic spectrum of WWOX-related neurological disorders, encompassing a continuum from mild to severe developmental epileptic encephalopathies. Significantly, all neurons with WWOX mutations demonstrated hyperexcitability, providing the first evidence that SCAR12 organoids exhibit increased activity compared to baseline, although they are less active than neurons derived from WOREE patients. Furthermore, the neurons derived from patients displayed an “immature” gene expression profile—termed “early” due to the overall immaturity of the organoids—suggesting potential delayed differentiation. While both SCAR12 and WOREE shared excitability defects, only WWOX-KO and WOREE organoids exhibited significant reductions in SATB2^+^ and CTIP2^+^ neurons. Such disruptions in laminar identity could indicate a broader defect in neuronal maturation and corticogenesis beyond the excitability defects, correlating with the severity of WWOX mutation. It is important to note that due to the limitations of this model, we could not determine if given additional time, affected neurons can obtain a late-stage neuronal signature, leaving this question unanswered.

A possible explanation to the observed phenotypic differences between the mutants could relate to the presence of key players in neuronal development, downstream of MYC, that are differentially affected upon WWOX deficiency, in a genotype dependent pattern. An interesting future avenue would be to explore top hits identified in our MYC ChIP-seq data on WWOX-KO, Fig. EV7D. For instance, BACH2, which we determined to be enriched in MYC binding in WT but not WWOX-KO NSCs, has been established to promote p21 expression and consequent cell cycle exit in neuroblastoma cells, whereby it controls neuronal differentiation in a temporal pattern (Shim *et al*, 2006).

We finally evaluated a neuron-specific gene therapy previously developed by our team, which had shown efficacy in *Wwox*-null mice (Repudi *et al*, 2021a). Although neuronal WWOX expression was not detected in neural organoids at this point, the success of this therapy, together with evidence for the ability of WWOX to promote neuronal differentiation and neurite outgrowth in SH-SY5Y cells (Wang *et al*, 2012), this approach was speculated to be able to improve neuronal function also in earlier point. Importantly, in this study we applied an AAV9-hSynI–driven delivery system, offering a clinically relevant approach for targeting postmitotic neurons in human gene therapy applications. This therapy effectively normalized hyperexcitability in WOREE organoids and led to increased expression of neuronal markers such as CTIP2 and SATB2. Additionally, pathways associated with neuronal maturation, including oxidative phosphorylation and synaptic ribosomal biogenesis, were upregulated, along with downregulation of voltage-gated ion channels, suggesting consistency with decreased electrophysiological phenotypes and a potential shift away from premature gene-expression signatures. Remarkably, this restoration also influenced myelination signaling, a function previously associated with WWOX deficiency in both mice and oligocortical spheroids (Repudi *et al*, 2021b).

Another potential therapeutic approach arising from our findings could involve silencing *MYC* in RGs. However, the pervasive role of *MYC* in essential cellular processes—ranging from chromatin organization and transcriptional regulation to cell cycle progression and metabolic programming—poses significant challenges for its manipulation during neurogenesis. As highlighted earlier, *MYC* regulates key glycolytic genes critical for metabolic homeostasis during neurodevelopment, and its dysregulation has been linked to impaired differentiation and neuronal cell death (Zheng *et al*, 2016), thus modulating *MYC* could introduce considerable experimental variability and confounding effects. Considering this, an intriguing avenue for future research would be to identify specific downstream targets of *MYC* that can be selectively modulated in *WWOX*-deficient RGs, potentially offering a more precise and less disruptive therapeutic strategy.

Evidently, while our findings highlight RGs as principal affected populations of WWOX dysfunction, we acknowledge that other cell types may also be affected at later stages. Organoids are limited in their ability to fully model the maturation and contribution of glial and non-neural lineages, which may play important roles in disease onset and progression.

Collectively, our findings demonstrate that WWOX-mutations, associated with WOREE and SCAR12 syndromes, profoundly disrupt RG cell homeostasis and differentiation, and neuronal development, contributing to neurodevelopmental abnormalities. Our data suggests that these effects are driven, at least in part, by MYC overexpression and altered cell cycle dynamics, potentially regulated by Wnt signaling. In WWOX-deficient neural organoids, we observed hyperexcitability and delayed neuronal development. Notably, a neuron-specific gene therapy using AAV9-WWOX successfully rescued these deficits, normalizing hyperexcitability and promoting neuronal maturation. These results indicate that WWOX loss of function disrupts neurogenesis by dysregulating MYC, impairing neuronal differentiation and function. Our findings highlight the therapeutic potential of AAV9-WWOX in mitigating developmental deficits and highlight the important non-cell-autonomous role of WWOX in brain development.

## Materials and Methods

### Cell Culture and Maintenance

All iPSCs and hESCs (WiBR3) were maintained in 5% CO2 on irradiated DR4 mouse embryonic fibroblast (MEF) feeder layers. Cells were cultured in FGF-2/KOSR medium consisting of DMEM-F12 (Gibco; 21331-020 or Sartorius; 01-170-1A) supplemented with 15% Knockout Serum Replacement (KOSR, Gibco; 10828-028), 1% GlutaMax (Gibco; 35050-038), 1% MEM non-essential amino acids (NEAA, Sartorius; 01-340-1B), 1% Sodium-pyruvate (Sartorius; 03-042-1B), 1% Penicillin-Streptomycin (Sartorius; 03-031-113), and 10 ng/mL bFGF (Peprotech; 100-18B). Medium was changed daily, and cultures were passaged every 5-7 days by TrypLE™ (Gibco; 12604013). Rho-associated kinase inhibitor (ROCKi, Y27632) (Cayman; 10005583) was added at 10 µM for the first 24-48 hours post-passaging. For RNA or protein isolation, hPSCs were passaged onto Matrigel-coated plates (Corning; 356231) and cultured in NutriStem hPSC XF medium (Sartorius; 05-100-1A).

### Transfection and Gene Editing

For transfection, iPSCs were pre-treated with 10 µM ROCKi for 24 hours. Cells were detached using TrypLE, resuspended in PBS (with Ca2+ and Mg2+), and mixed with 100 μg total DNA constructs. Electroporation was performed using the Gene Pulser Xcell System (Bio-Rad) at 250 V, 500 μF, in 0.4 cm cuvettes. Transfected cells were plated on MEF feeder layers in FGF/KOSR medium supplemented with ROCKi. For WWOX Knockout, the px330 plasmid containing sgRNA targeting WWOX exon 1 was co-electroporated with pLKO.1 puro plasmid (1:10 ratio). After 48 hours, the cells were treated with puromycin at a concentration of (1 μg/ml) for 72 hr, after which fresh medium was added. The cells were grown until distinct colonies appeared and were isolated for further expansion, after approximately 7 days. Gene editing was validated by Western blot. sgRNA sequence (*hWWOX*): 5′-CACCGCATGGCAGCGCTGCGCTACG-3′

### Reprogramming of somatic cells

Skin biopsies and blood samples were obtained with informed consent under approval from the Kaplan Medical Center Helsinki committee.

Reprogramming was conducted using Sendai virus Cyto-Tune-iPS2.0 Kit according to manufacturer’s instructions.

#### Peripheral blood mononuclear cells (PBMCs)

PBMCs were isolated from a whole-blood sample obtained from a healthy adult male using ficoll gradient and cultured in StemPro-34™ medium supplemented with cytokines (SCF, FLT-3 ligand, IL-3, and IL-6). On day 0, cells were infected with reprogramming viruses. On day 3, cells were transferred to MEF-coated plates. By day 7, reprogramming cells were observed, and medium was gradually changed to mTeSR, supplemented with ROCKi for the first 48 hrs. On day 16, colonies were picked, expanded, validated for pluripotency markers, and sequenced for WWOX mutations. For further details, kindly refer to (*23*).

#### Skin fibroblasts

Primary fibroblasts obtained from donors were cultured in Fibroblast medium consisting of DMEM supplemented 15% Fetal Bovine Serum (FBS), 1% NEAA, 1% sodium pyruvate, 1% GlutaMax, 100 μM 2-mercaptoethanol without antibiotics. 2-4 days prior to reprogramming, the fibroblasts were seeded in 6-well plated coated with 2% gelatin solution and grown to 60-70% confluency. On day 0, cells were infected with Sendai viruses. The next day, the medium was replaced by fresh fibroblast medium. On days 2-6 the medium was changed daily. On day 7, fibroblasts were harvested using Trypsin type C solution (Sartorius; 03-053-1B) and seeded onto MEF-coated plates. On day 8, the medium was changed to FGF-2/KOSR medium as described above, with 10 µM ROCKi added for the first 3 days. The medium was replenished daily. By day 21, small colonies appeared, and by day 28 the cells for colony picking and further expansion. All lines were stained and validated for pluripotency markers and sequenced for WWOX mutations.

All experiments conformed to the principles set out in the WMA Declaration of Helsinki and the Department of Health and Human Services Belmont Report.

### Cerebral organoid generation and culture

Cerebral organoids were generated from iPSCs as previously described (*23*):

All iPSCs were maintained on mitotically inactivated MEFs. Prior to protocol initiation, cells were passaged onto 60mm plates coated with MEFs and grown to 70-80% confluency. Embryoid Body (EB) Formation: On day 0, hESC colonies were dissociated to single cells using 1 mg/mL Dispase II solution (Sigma; D4693) followed by TrypLE treatment. 4,500 cells were seeded per well in ultra-low attachment 96-well U-bottom plates (S-Bio Prime; MS-9096UZ) in hESC medium (DMEM/F12, 20% KOSR, 1% GlutaMax, 1% NEAA, 1% Penicillin-Streptomycin, 100 µM 2-mercaptoethanol, 5 ng/mL bFGF, and 10 µM ROCKi). EBs were fed every other day for 5 days.

#### Neural Induction

On day 6, medium was replaced with Neural Induction (NI) medium (DMEM/F12, 1% N2 supplement, 1% GlutaMax, 1% MEM-NEAA, 1 µg/mL Heparin). NI medium was changed every other day until neuroepithelium establishment (days 11-12). Matrigel Embedding and Differentiation: Well-developed EBs were embedded in Matrigel droplets and transferred to 90mm non-treated culture dishes with Cerebral Differentiation Medium (CDM: 1:1 DMEM/F12 and Neurobasal medium, 0.5% N2, 1% B27 without vitamin A, 1% GlutaMax, 1% penicillin/streptomycin, 0.5% NEAA, 50 µM 2-mercaptoethanol, 2.5 µg/mL insulin, and 3 µM CHIR-99021).

#### Maturation

From day 16, organoids were cultured on an orbital shaker (Thermo MaxQ 2000 CO2 Plus Orbital Shaker) at 70 rpm in Cerebral Maturation Medium (CMM: DMEM/F12 and Neurobasal medium, 0.5% N2, 1% B27 with vitamin A, 1% GlutaMax, 1% penicillin/streptomycin, 0.5% NEAA, 50 µM 2-mercaptoethanol, 2.5 µg/mL insulin, 400 µM vitamin C, and 12.5 mM HEPES buffer, without CHIR-99021). From week 6, 1% Matrigel was added to the medium. Medium was changed every ≤ 72 hr, and organoids were transferred to fresh sterile plates every 30 days. All of the described media were filtered through a 0.22µm filter and stored at 4°C until usage.

### AAV9 infection

AAV9 viruses were obtained as described previously (*59*).

For infection, AVV9-WWOX and AAV9-EGFP viruses were diluted in CMM without antibiotics to a final concentration of 3x10^10^ viral genomic copies (GC)/μL. The medium volume was calculated as 100 μL/organoid. Then, ∼25 organoids at week 11 were transferred to low-attachment 6-well plates, the spent medium was carefully removed and 1.5 ml of virus-containing CMM was added. The organoids were cultured stationary for 5 hr in a humidified incubator, followed by the addition of extra 1.5 ml of virus containing medium was added before over-night incubation. The following morning, the medium was diluted 1:1 with fresh CMM and the organoids were incubated for an additional 24 hr. After ∼48 hr after the original infection, a complete medium change was performed and the organoids were cultured as described in an earlier section, up until harvesting for analysis at week 16.

### NSCs differentiation

Neural stem cells (NSCs) were generated from hESCs using a modified dual-SMAD inhibition protocol (*45*). Briefly, WiBR3 hESCs cultured on a MEF-feeder layer were released from MEFs and dissociated to single cells as previously described for organoids, and seeded on Matrigel-coated plates at 50,000 cells/cm^2^ in MEF-conditioned media supplemented with 10 ng/mL FGF-2 and 10 μM ROCKi. Neural induction was initiated the following day using a NSCs medium containing DMEM/F12:Neurobasal (1:1), 2% B27 supplement, 1% N2 supplement, 1% Penicillin-Streptomycin, 1% NEAA and 100 μM β-mercaptoethanol. From day 1 to 10, dual SMAD inhibitors 100 nM LDN-193189 (Axon Medchem; 1527) and 20 µM SB-431542 (Sigma; S4317) were added along with 1 µM XAV-939 (Peprotech; 2848932). From day 10 to 20, SMAD inhibitors were withdrawn, and 50 nM SAG (Sigma; 566660) was added with 1 µM XAV-939. The medium was changed daily. By day 20-22, cells had become NSCs, were passaged and cryopreserved, and marker expression was validated using immunofluorescence. NSCs were maintined on Matrigel-coated plates in NSCs medium supplemented with N2 (1%), B27 (2%), 40 ng/ml EGF (Peprotech; AF-100-15), 40 ng/ml FGF2, and 1.5 ng/mL hLIF (Sigma; L5283). Medium was changed every 2-3 days, and cells were passaged at 60-70% confluence using TripLE at room temperature.

### Immunofluorescence

Organoids fixation and immunostaining were performed as previously described (*23*). Briefly, for histological analysis, organoids were rinsed thrice in PBS before being immersed in 4% ice-cold paraformaldehyde for 45 minutes on an orbital shaker for fixation. Following three cold PBS washes, the organoids were subjected to cryoprotection by overnight immersion in a 30% sucrose solution. Subsequently, the organoids were embedded in OCT compound, rapidly frozen on dry ice, and sectioned at 10 μm thickness using Leica CM1950 cryostats.

Immunofluorescent staining was performed on these sections following a standard procedure. First, they were brought to room temperature and rehydrated in PBS. The sections were then permeabilized using 0.1% Triton X in PBS (PBT) and blocked for 60 minutes in a solution containing 5% normal goat serum (NGS) and 0.5% BSA in PBT. Primary antibodies, diluted in the blocking solution, were applied to the sections and incubated overnight at 4°C. The following day, the sections were washed three times in PBS with 0.05% Tween-20 (PBST) under gentle agitation on an orbital shaker. Secondary antibodies and Hoechst33258, diluted in blocking buffer, were then applied for 90 minutes. After four PBST washes while place on an orbital shaker, coverslips were mounted using

Immunofluorescence Mounting Medium (Dako; s3023). Imaging was done using multiple systems, an Olympus FLUOVIEW FV1000 confocal laser scanning microscope, a ZEISS LSM 980 Airyscan2 confocal, and a 3DHistech Pannoramic scanner, with image processing carried out using the associated Olympus FLUOVIEW software, zen-lite for LSM980 or NIH ImageJ. Quantification of marker-positive cells was conducted using QuPath v0.5.1, followed by statistical analysis as described in later sections.

### Cell-attached recordings

Cell-attached recordings were performed as previously described (*23*), briefly: Spontaneous neuronal activity from organoid neuronal populations was recorded using blind patch-clamp techniques. Glass electrodes (approximately 7 MOhm resistance) were fabricated from filamented, thin-walled borosilicate glass (1.5 mm outer diameter, 0.86 mm inner diameter; Hilgenberg GmbH, Malsfeld, Germany) using a vertical two-stage puller (PC-12, Narishige, East Meadow, NY). The electrodes were filled with an internal solution comprising (in mM): 140 K-gluconate, 10 KCl, 10 HEPES, 10 Na2-Phosphocreatine, 0.5 EGTA, adjusted to pH 7.25 with KOH.

Electrodes were inserted at a 45° angle relative to the organoid surface. During recordings, organoids were maintained in Cerebral Maturation Medium (CMM) without Matrigel at 35°C. An increase in pipette resistance to 10-200 MOhm typically coincided with the appearance of spikes, which served as the criterion to initiate recording. All recordings were acquired using an intracellular amplifier in current-clamp mode (Multiclamp 700B, Molecular Devices), sampled at 10 kHz (CED Micro 1401-3, Cambridge Electronic Design Limited), and filtered with a high-pass filter to isolate neuronal spikes from field potentials. Analysis of cell-attached recordings was performed using custom-written MATLAB code. Spikes were extracted from raw voltage traces by applying a threshold set well above the background noise level. The average firing rate for each recorded cell was calculated over a 4-minute recording period.

### Ca^2+^ Imaging

Calcium imaging was performed for neural organoids at 16 weeks post-differentiation. In brief, Calcium transients were observed by incubating cerebral organoids for an hour with 8 µM Fluo-5 AM (Abcam; ab241083) in the ACSF, i.e., recording solution containing (in mM) 139 NaCl, 10 HEPES, 4 KCl, 2 CaCl2, 10 D-glucose, and 1 MgCl2 (pH 7.5, osmolarity adjusted to 310 mOsm). After one hour of incubation, the organoids were placed in a recording chamber filled with clean ACSF that had been pre-warmed to 37 °C, and imaging of calcium transients was performed on a SliceScope Pro 3000 microscope (Scientifica) with a CCD digital camera (Scientifica, SciCam Pro). The images and data were captured and processed using a custom-written GUI in MATLAB 9.8 (R2021a, MathWorks). Recording of single cells were performed at an acquisition rate of 10 Hz. Time series data of calcium imaging recordings was analyzed using a custom-written Python script. The intensity of shifts in fluorescence levels was calculated by measuring the changes between the values of two adjacent image frames. The fluorescence ratio (Δf/f0) was calculated by dividing the signal’s maximum amplitude by the baseline fluorescence which was calculated as the average signal when the cell was inactive.

### Immunoblot analysis

Organoids homogenized in lysis buffer containing 50 mM Tris (pH 7.5),150 mM NaCl, 10% glycerol, and 0.5% Nonidet P-40 (NP-40) that was supplemented with protease and phosphatase inhibitors and incubated on ice for 15 minutes. The samples were than centrifuged at (14,000 rpm) for 15 minutes at 4 degrees. The liquid phase was collected, and protein concentrations were measured using the Bradford protein assay (Bio-Rad, #5000006). Samples were prepared using 30-50 ug of protein in 1:4 ratio with sample buffer. Western blotting was performed under standard conditions at a constant current of 400 mA, 100-110 V, for 2-3 hours at room temperature. Gels were transferred using Bio-rad’s semi-dry transfer system for 15 minutes. Blots were repeated and quantified 2–3 times per experiment in Bio-Rad’s Image Lab software. Representative images of those repeated experiments are shown.

### .RNA extraction, reverse transcription-PCR, and qPCR

Total RNA was isolated using Bio-Tri reagent (Biolab; 9010233100) and cleaned using the Zymo Direct-zol RNA MiniPrep kit (Zymo; R2050) as described by the manufacturer. 0.25-1 µg of RNA was used to synthesize cDNA using a qScript cDNA Synthesis kit (QuantaBio; 95047). qRT-PCR was performed using Power SYBR Green PCR Master Mix (Applied Biosystems; AB4367659). All measurements were performed in triplicate and were standardized to the levels of either HPRT or UBC. All primer sequences used are noted in Table S2.

### Library preparation and Single-cell RNA-seq (scRNA-seq) (10x Chromium)

Neural organoids were dissociated into single cells papain-based enzymatic digestion method, based on a previously published protocol (*36*), with the following variations: 3-4 organoids from each hPSCs line were transferred to a 60 mm culture dish containing 4 mL of pre-warmed Earle’s Balanced Salt Solution (EBSS) (Sartorius; 02-010-1A). The organoids were manually dissected into small pieces and transferred into a new 60 mm dish containing 2.5 mL of Enzymatic dissociation solution [composed of 27.3 U/mL papain solution (Worthington; LS003126) supplemented with 200 U/mL of DNase I solution (Sigma; 10104159001) and 10 μM ROCKi in Hanks’ Balanced Salt Solution (HBSS) (Sartorius; 02-015-1A)], and incubated on an orbital shaker (70 rpm) in a humidified incubator for 30 minutes at 37°C. Following digestion, the tissue was collected in a 15 mL tube, and 5 mL of warm EBSS was added. The mixture was triturated 20 times using a 10 mL pipette, after which it was allowed to settle until all pieces sank. The resulting supernatant containing dissociated cell suspension was carefully transferred to a new 15 mL tube, avoiding any undigested tissue pieces. To the cell suspension, 5 mL of EBSS, 3 mL of Enzyme inhibiting solution [containing 10 mg/mL Trypsin inhibitor solution (Sigma; T9253) and 10 mg/mL BSA in EBSS) were added and gently mixed. Cells were pelleted by centrifugation at 300 g for 7 minutes at room temperature. The liquid was aspirated, and 5 ml of warm EBSS was added, the cells were centrifuged again and resuspended in cold 0.1% BSA solution + 10 μM ROCKi in PBS. The cells were passed through a 40 μm strainer. Cell viability and concentration were determined using trypan blue staining. Finally, cells were diluted in ice-cold DPBS + 0.1% BSA + ROCKi solution to a concentration of 100 cells/μL and kept on ice until droplet collection.

Library preparation and sequencing was done by the Hebrew University’s Core facility following a standard protocol. Briefly, single-cell suspensions, prepared as described in the previous section, were processed for RNA sequencing using the 10x Genomics platform. Libraries were generated using the Chromium Next GEM Single Cell 3ʹ GEM, Library & Gel Bead Kit v3.1 (10× Genomics, Pleasanton, CA, USA), following the manufacturer’s protocol. Libraries that had passed quality control were then sequenced on an Illumina NovaSeq 6000 platform (Illumina, San Diego, CA, USA). Sequencing parameters were set to generate paired-end reads with the following specifications: 28 base pairs (bp) for Read 1 and 90 bp for Read 2. Read 1 contained cell-identifying barcodes and unique molecular identifiers (UMIs), while Read 2 corresponded to mRNA transcript sequences. The sequencing depth was targeted to achieve approximately 50,000 reads per cell, overall obtaining about 1B reads, which were evenly distributed between the 5 libraries.

### Single-cell data preprocessing

Organoids from five different conditions (WT = 6,792; WWOX-KO = 4,124; SCAR12 = 4,454; WOREE = 7,687; WOREE-WWOX = 5,151) were analyzed jointly, resulting in a total of 28,208 cells. After removing undetected genes and cells with high mitochondrial content or total count numbers, we retained 26,352 high-quality cells.

Normalization, batch correction, clustering, dimensionality reduction, and visualization steps were performed using the computational tool ACTIONet (*108*), release version available at (https://github.com/shmohammadi86/ACTIONet/tree/R-release), using the functions *normalize.ace*, *reduce.and.batch.correct.ace.Harmony*, *runACTIONet*, *clusterCells*, and *plot.ACTIONet* for visualization. Batch effects were corrected using Harmony accounting for the condition of the organoids (WT, WWOX-KO, SCAR12, WOREE, WOREE-WWOX). Clusters were identified using the Leiden algorithm with default parameters implemented in ACTIONet. 12 cell clusters were annotated based on manually curated marker genes and transcriptional signature comparisons with a human primary single-cell data covering early (*40*), mid (*37*, *39*), and late (*38*) fetal cortical development. Cluster profiles were generated as the average expression across cells within each cluster. Signatures were calculated using the total pairwise fold-change sum of a given cluster profile relative to others. Statistical overrepresentation analyses of marker genes manually curated from the literature on these signatures were used to interpret and annotate the clusters. Correlation of cluster signatures with cell type signatures from single-cell data from the primary samples was used to further support cluster annotations. Clusters that could not be confidently mapped to specific cell types and/or presented offset differentiation or stress signatures were flagged and not considered for downstream analyses. A total of 18,007 cells annotated as radial glia, neuronal progenitors, or neurons (WT = 5,649; WWOX-KO = 3,020; SCAR12 = 3,422; WOREE = 5,916; WOREE_WWOX_ = 3,763) passed all quality controls.

### Differential Gene Expression

Differential gene expression (DGE) analysis was performed using nonparametric Wilcoxon tests as implemented in the R package *presto* (*109*). Genes with a log fold change of ±0.5 and a p-value < 0.05 were considered differential. DGE analysis was performed separately for radial glia cells and for neuronal progenitor and neuronal cells.

### WWOX and MYC expression in-vivo

Human single-cell data at post-conceptional week 16 (Trevino *et al*, 2021b) and tissue-level cell sorted data from week 13 to 20 were used to asses WWOX gene expression in vivo. Expression at single-cell level was assessed by projecting WWOX expression onto a 2D manifold of the data using the uniform manifold approximation method (UMAP) as implemented in ACTIONet. The same approach was applied to visualize radial glia (VIM, SOX2), and neuronal (TBR1, SATB2, NEUROD6) markers for comparison. Expression values at single-cell level were further compared and correlated with annotated cell types. Expression in FACS-sorted data was assessed by comparing relative expression values of WWOX across fetal cell types. Expression in tissue-level data was assessed across either developmental time or anatomical zones. Temporal expression of WWOX and MYC was assessed in human forebrain developmental data from embryonic stages to adulthood (Cardoso-Moreira *et al*, 2019b) and in reference time-series of cortical organoid differentiation (Gordon *et al*, 2021). Observed trends are based on smoothed normalized expression estimated using Tukey’s methods as implemented in the *smooth* function from the stats package in R. Anatomical zone expression was assessed in laser microdissected human fetal data (Miller *et al*). Relative expression values are based on mean-subtracted values across all cortical zones. Log-transformed TMM normalization as implemented in the R package *edgeR* was used to account for library size variation. Replicates were averaged only for samples between days 25 and 463 of differentiation. All heatmap visualizations were performed using the R package *Complex Heatmap*.

### Cell cycle phase and potency state inference

To estimate the cell cycle phase of radial glia cells from single-cell transcriptomic data, the *tricycle* R package was used with default parameters. The cell phase discretization was defined based on the cell cycle timepoint of each single cell as follows: G1/G0 0π - 0.5π and > 2π, S 0.5π - 1π, G2M 1π - 1.5π, M 1.5π - 2π. The cell cycle proportions between WWOX-KO and WT were compared by calculating the proportion of cells in each phase relative to the total number of cells in each condition and by computing the log2 fold change between WWOX-KO and WT conditions. To assess the prevalence of MYC expression across cell cycle phases, gene expression levels were correlated to the cell cycle phase estimations and the contribution for each condition was estimated. The potency potential os single cells was estimated from transcriptomic data using the CytoTrace2 package as described in (Kang *et al*, 2024). Cells were categorized into three potency levels: Multipotent (high potency), Oligopotent/Unipotent (medium potency), and Differentiated (low potency).

### Overrepresentation analysis

Statistical overrepresentation estimations of conditions and cell cycle phases across potency states in radial glia cells are based on binomial tests on the 3 potency categories.

Enrichment analysis of pathways of interest obtained from the *EnrichR* database (https://maayanlab.cloud/Enrichr/#libraries), KEGG, and REACTOME databases, was assessed on ranked transcriptional signatures of potency states by using the assess.geneset.enrichment.from.scores function of the ACTIONet package. MYC positive and negative regulon genesets were obtained from an integrative database for the estimation of human transcription factor activities (*VIPER*). The p-values were adjusted using Bonferroni correction, and all scores calculated as -log10 (p-value) were capped to a maximum value of 10 for visualization purposes. Gene Set Enrichment Analysis (GSEA) was performed over ranked scores using the *fgsea* R package . The pathway database *SynGO* was used to estimate enrichment over log2 fold changes between experimental conditions in neuronal cells.

### Neuron Analysis

Neurons and intermediate progenitors from all conditions were isolated and re-analyzed following the ACTIONet standard pipeline described above. To interpret neuronal subtypes, reference neuronal cell type-specific signatures were computed from (*58*) and correlated to the organoid cluster signatures. Neuron clusters were interpreted and annotated based on reference correlations and preferential expression of marker genes. For clusters expressing markers associated with both excitatory and inhibitory neurons and showing ambiguous mapping in the correlation matrix, a “mixed neurons” category was assigned. The resulting 7 subtypes were divided into early excitatory neurons (EN-IT Immature, EN Newborn, and IPC-EN) and late excitatory neurons (EN-L5-ET, EN-L5-CT, EN-L6-CT, and Mixed Neurons).

### Pseudobulk analysis

All pseudobulk profiled were estimated from the single-cell data as the sum of raw counts over cells of a given condition and normalized as log-transformed counts per million (cpm).

### Dorso-ventral gene expression

To investigate the regional patterning of our organoids, we evaluated the expression of dorsal genes (PAX6 and EOMES) and ventral genes (GSX2, DLX2, SLC32A1 -VGAT-) reported in (Renner *et al*, 2017) in the pseudobulk transcriptional profiles of each condition (WT, WWOX-KO, SCAR12, and WOREE). The dorsal / ventral genes ratio was computed using the average expression of the same genes.

### MYC and WWOX interactome analysis

To build the interactome network of WWOX and MYC, we downloaded the high-confidence (e.g., A and B categories) MYC interactions from the R package dorothea (https://saezlab.github.io/dorothea/). The resulting list was filtered for differentially expressed genes from comparing WT and WWOX-KO radial glia. Both upstream and downstream transcription factors were selected. For the WWOX genes we selected those upstream, since WWOX is not a transcription factor and does not regulate other genes.

Statistical overrepresentation estimations of conditions and cell cycle phases across potency states in radial glia cells are based on binomial tests on the 3 potency categories.

Enrichment analysis of pathways of interest obtained from the *EnrichR* database , KEGG, and REACTOME databases, was assessed on ranked transcriptional signatures of potency states by using the assess.geneset.enrichment.from.scores function of the ACTIONet package. MYC positive and negative regulon genesets were obtained from an integrative database for the estimation of human transcription factor activities (*VIPER*)human transcription factor activities (*VIPER*). The p-values were adjusted using Bonferroni correction, and all scores calculated as -log10 (p-value) were capped to a maximum value of 10 for visualization purposes. Gene Set Enrichment Analysis (GSEA) was performed over ranked scores using the *fgsea* R package. The pathway database *SynGO* was used to estimate enrichment over log2 fold changes between experimental conditions in neuronal cells.

### Chromatin immunoprecipitation sequencing (ChIP-seq)

ChIP-seq was performed as previously described [40]. Briefly, WiBR3 hESC-derived NSCs of wildtype and WWOX-KO (∼10^7^ cells) were crosslinked with 1% formaldehyde (methanol free, Thermo Scientific 28906) for 10 min at room temperature and quenched with glycine, 125 mM final concentration. Fixed cells were washed twice in PBS and incubated in lysis buffer (10 mM EDTA, 0.5% SDS, 50 mM Tris-HCl pH = 8, and protease and phosphatase inhibitors) for 30 min on ice. Cells were sonicated using bioruptor sonicator to produce chromatin fragments of ∼200–300 bp. The sheared chromatin was centrifuged 10 min at 20,000 × g. From the supernatant, 2.5% were saved as input DNA and the rest was diluted in dilution buffer (50 mM TRIS-HCl pH8, 0.01% SDS, 150 mM NaCl, and 1% Triton X-100). The chromatin was immunoprecipitated by incubation with 5 μl of anti-c-Myc antibody (rabbit polyclonal c-Myc (9402)-Cell Signaling,). Immune complexes were captured with protein G Dynabeads. Immunoprecipitates were washed once with low salt, twice with high salt buffer, and twice with LiCl buffer, and twice with TE buffer. The chromatin was eluted from the beads with 240 μl of elution buffer (100 mM sodium bicarbonate and 1% SDS) and incubated overnight at 65 °C to reverse the cross-linking. Samples were treated with proteinase K at 45 °C for 2 h. DNA was precipitated by phenol/chloroform/isoamylalcohol extraction. The ChIPed and the Input DNA were used to prepare libraries by Kappa Hyperprep kit and sequenced in Nextseq (Illumina).

### Chromatin immunoprecipitation sequencing (ChIP-seq) analysis

Raw paired-end reads were trimmed with Trim Galore! and aligned to the human reference genome (hg38) using HISAT2, retaining only uniquely mapped reads with mapping quality ≥10. BAM files were sorted, indexed, and filtered with SAMtools. Normalized coverage and ChIP/Input enrichment tracks were generated with deepTools (bamCoverage and bamCompare) using read count normalization. Promoter-level enrichment (±1 kb from TSS) was quantified from normalized bigWig files with bigWigAverageOverBed (UCSC tools). To assess the preferential binding of MYC on promoters of manually curated genes associated with immature excitatory neurons, the z-score of WT and KO was computed on the null distribution (WT pval = 2.03e-05). The MYC binding density of the top 20 immature excitatory neuron genes between WT and KO were shown in the heatmap.

### Statistical analysis

Results of the experiments were expressed as mean ± SEM. Statistical significance was determined, after confirming normal distribution using the Wilk-Shapiro test, by either using the one-way ANOVA test, correcting for the multiple comparisons with Tukey’s or Dunnett’s multiple comparisons tests, or by using multiple t-tests, correcting for multiple comparisons by controlling the False Discovery Rate (FDR) using the Benjamini–Hochberg procedure. The specific test used in each experiment is described in the corresponding figure legend. *P*-value cutoffs for statistically significant results were used as following: n.s (non-significant), *p ≤ 0.05, **p ≤ 0.01, ***p ≤ 0.001, ****p ≤ 0.0001. Statistical analysis and visual data presentation were performed using GraphPad Prism 9. No randomization or blinding was applied in this study. The experiments were performed on several biological replicates, with at least two clones of each hPSC lines used for each genotype (except for the WiBR3 WT and JH-iPS11 WT lines).

## Supporting information

Expanded View

## Data availability

The single-cell RNA sequencing data have been uploaded to ArrayExpress with accession number: E-MTAB-14792.

We used raw publicly available data from single-cell and bulk sequencing from these publications ((Polioudakis *et al*; Ramos *et al*; Trevino *et al*, 2021a; Bruggen *et al*; Cardoso-Moreira *et al*, 2019b; Song *et al*; Miller *et al*)

## Acknowledgments

We thank all members of the Aqeilan’s and Davila-Velderrain’s laboratories for technical help and fruitful discussion. We are grateful to Prof. Nissim Benvenisty and Dr. Tamar Golan-Lev from the Azrieli Center for Stem Cells and Genetic Research, and to Dr. Abed Nasereddin and Dr. Idit Shiff from the Genomic Core Facility for their help. Special thanks go to all WOREE and SCAR12 syndrome patients and families for their enduring support and dedication.

## Author contributions

Conceptualization: DS, AZ, DA, JDV, RA

Methodology: DS, AZ, DA, OH, IR, IK, KM, SS

Investigation: DS, AZ, DA, JDV, RA

Visualization: DS, AZ, DA, JDV

Supervision: JDV, RA

Writing—original draft: DS, AZ, DA, JDV, RA

## Expanded View figures

**Fig. EV1.**
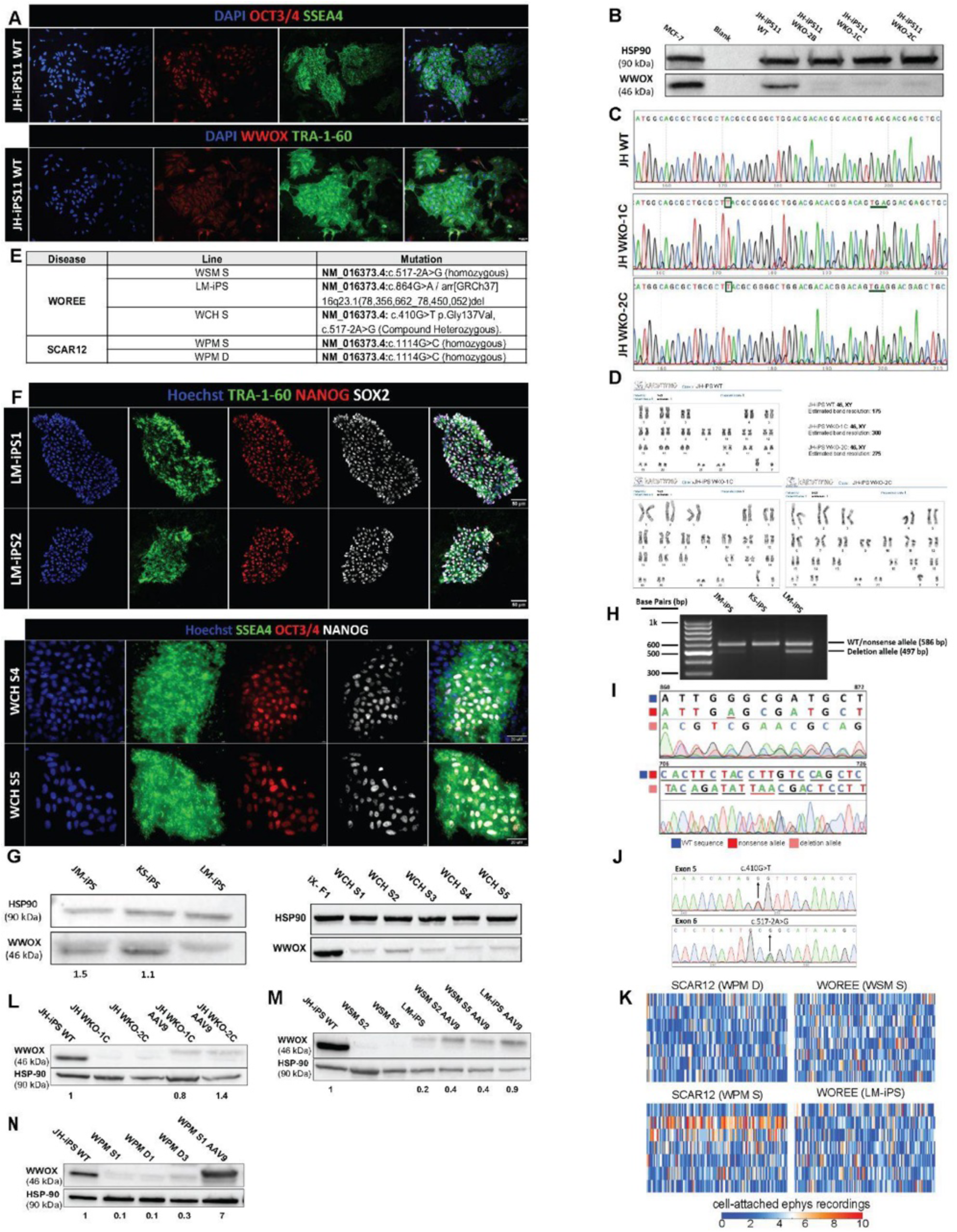
iPSCs generation and CRISPR/Cas9 editing. **(A)** Immunofluorescent staining of pluripotency markers in WT iPSC line established JH-iPS11, and for WWOX. **(B)** Immunoblot analysis confirming the successful knock out of the WWOX gene in JH-iPS11 cells. WKO; WWOX knockout, followed by a clone-specific label. **(C)** Sanger sequencing of JH-WT, JH WKO-1C and JH WKO-2C showing indels (boxed) introducing a premature stop codon (horizontal line). **(D)** G-band karyotyping reports of JH-WT, JH WKO-1C and JH WKO-2C indicating no detectable chromosomal abnormalities. **(E)** Summary of WOREE and SCAR12 patients included in the study and their exact mutations. **(F)** Immunofluorescent staining of pluripotency markers in a WOREE patient-derived iPSCs lines, LM-iPS and WCH, harboring different complex heterozygous mutation in the WWOX gene. **(G)** Immunoblot analysis confirming WWOX expression levels in iPSCs of the LM-iPS patient and the healthy parents (JM-iPS and KS-iPS) (left panel), and in WCH patient iPSC clones compared to a wildtype control**\*. (H)** Gel electrophoresis after PCR amplification of a cDNA fragment spanning parts of exons 5-8 of the WWOX gene, showcasing the absence of exon 6 in one parental allele. **(I)** Sanger sequencing of the cDNA fragments shown in (E). The top panel shows the maternal nonsense mutation (c.864G>A) compared to a reference genome, and the sequence shift in the paternal allele due to the deletion of 0.9 Mbp. The bottom panel shows the premature stop codon (underlined in red) resulting from the deletion and frameshift in the paternal allele. The numbers in the top corners denote the position in the reference WWOX coding sequence. **(J)** Sanger sequencing of WCH S4 clone showing the maternal missense mutation in exon 5 (c.410G>T), and the paternal splice site mutation in in exon 6 (c.517-2A>G). **(K)** Raster plots demonstrating selected neuronal firing from SCAR12 (WPM D and WPM S) and WOREE (WSM S and LM-iPS) week 7 organoids over 4 minutes of cell-attached recording, related to Figure 1C. **(L)** Immunoblot analysis of JH-WT, JH-WKO and JH-WKO AAV9-hSyn-hWWOX cerebral organoids from week 16, confirming knockout levels of WWOX and its rescue**. (M)** Immunoblot analysis of WWOX levels in WOREE lines and their AAV9 rescue included in the scRNA-seq experiment in week 16 cerebral organoids. **(N)** Immunoblot analysis of WWOX levels in SCAR12 lines included in the sc-RNA-seq experiment in week 16 cerebral organoids.

**Fig. EV2.**
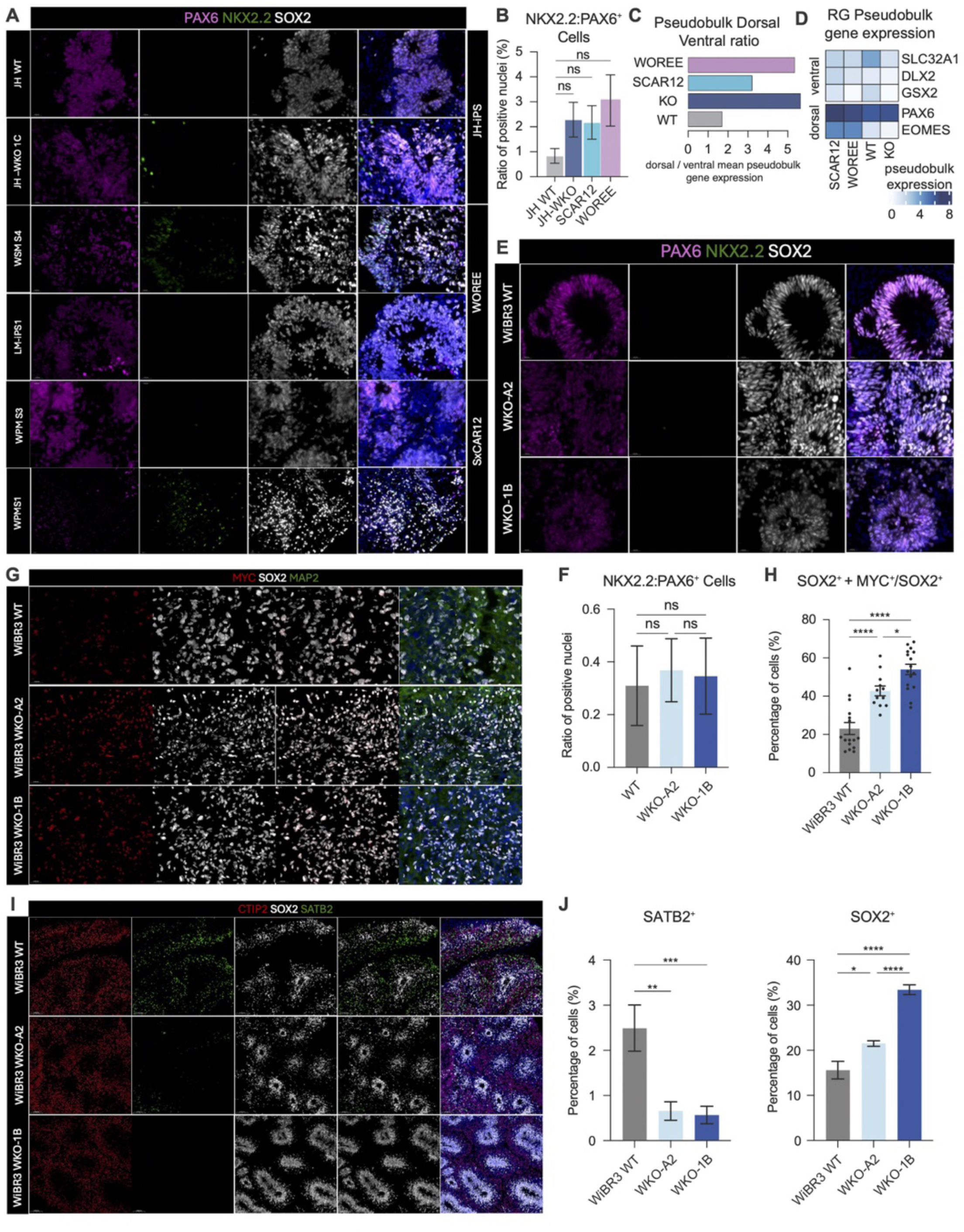
Characterization and validation of cerebral organoids. **(A)** Representative immunofluorescent staining of Pax6 (dorsal marker) and NKX2.2 (ventral marker) in all organoids included in the scRNA seq study at week 7 in vitro. **(B)** Quantification of (a) showing the ratio of NKX2.2: PAX6 positive cells revealing predominant dorsal origin with no significant differences in patterning among lines. **(C)** Analysis of cerebral organoids included in our scRNA-seq data showing increased proportion of dorsal relative to ventral markers expressed in all lines. **(D)** Pseudobulk analysis expression levels of known dorsal and ventral markers in all lines. **(E)** Representative immunofluorescent staining of Pax6 (dorsal marker) and NKX2.2 (ventral marker) of hESCs-derived cerebral organoids of WT and WWOX-KO at week 15 in vitro. **(F)** Quantification of (e) showing the ratio of NKX2.2: PAX6 positive cells revealing predominant dorsal origin with no significant differences in patterning among lines. **(G)** Immunostaining of week 15 hESCs-derived cerebral organoids at week 15 in vitro for SOX2 and MYC double positive populations. **(H)** Quantification of immunostaining in (g) showing expansion of MYC^+^ SOX2^+^ progenitor populations. **(I)** Immunostaining of cerebral organoids at week 15 for SOX2 marker for progenitors, and SATB2 and CTIP2 neuronal markers. **(J)** Quantification of SOX2+ and SATB2+ populations in (I) showing similar reduced differentiation capacities to our iPSC-derived cerebral organoids. *See Table EV1 for details on organoid numbers, sections, and batches. Data are presented as mean ± SEM. Statistical significance was determined using one-way ANOVA with Tukey’s multiple comparisons test. n.s (non-significant), *p ≤ 0.05, **p ≤ 0.01, ***p ≤ 0.001, ****p ≤ 0.0001*.

**Fig. EV3.**
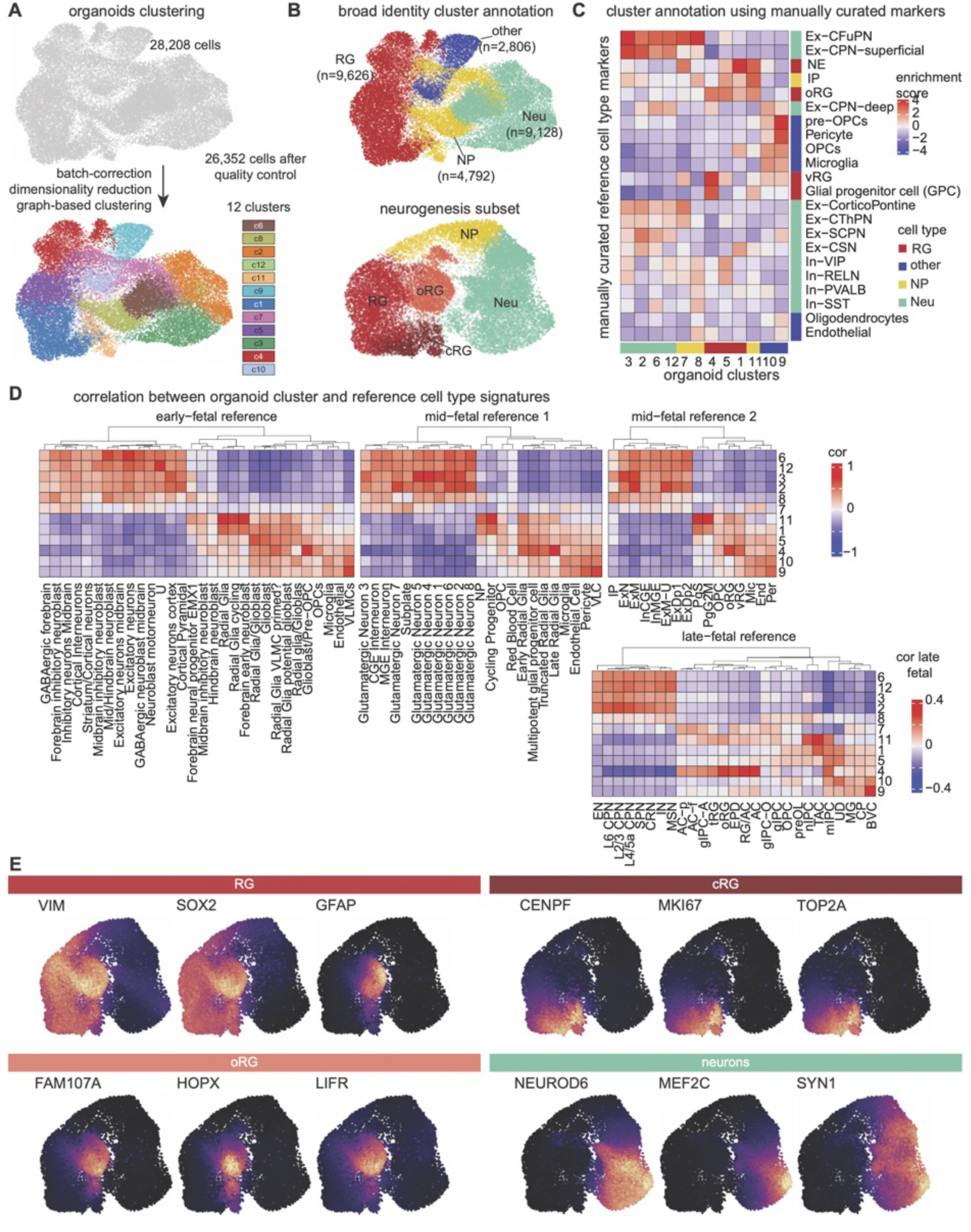
Cell type identification of scRNA-seq samples. **(A)** (Upper) UMAP of all organoids together after standard quality controls (removed 1856 low quality cells). (Lower) Clustering of cells recovering 12 groups. **(B)** Clusters colored by their general annotation, with RG accounting for 9,626 cells, NP 4,792 cells, Neu 9,128 cells and other 2,806 cells. **(C)** Heatmap of scaled enrichment score of manually curated marker genes (rows) in the 12 clusters (columns), all colored by their general cell type annotation. **(D)** Cluster signatures comparison to references fetal cortical development of different ages: early fetal (post conceptional week 08-10), mid-fetal 1 (w16-24), mid-fetal 2 (w15-16), late-fetal (w36). Correlation for early and mid-fetal from -1 to 1, correlation values for late-fetal from -0.4 to 0.4. **(E)** Feature plots in WT organoids for pan-RG, cRG, oRG and Neurons markers. References: early fetal (10.1016/j.devcel.2022.04.016), mid-fetal 1 (10.1016/j.cell.2021.07.039), mid-fetal 2 (10.1016/j.neuron.2019.06.011) and late-fetal (10.1038/s41467-022-34975-2, subject at w36). RG = Radial Glia, vRG = ventricular Radial Glia, oRG = outer Radial Glia, IP = Intermediate Progenitor, Ex-CFuPN = CorticoFugal Projection Neuron, Ex-CThPN = CorticoThalamic Projection Neuron, Ex-SCPN = SubCerebral Projection Neuron, Ex-CPN = Commisural (callosal) Projection Neuron, In = Interneuron, OPC = Oligodendrocyte Progenitor Cell, NE = Neuroepithelial cells, U = Unknown, VLMC = Vascular and Leptomeninges, CGE = Caudal Ganglionic Eminence, MGE = Medial Ganglionic Eminence, ExN = Excitatory Neuron, ExM = Migrating Excitatory neuron, ExM-U = Upper-Migrating Excitatory neuron, ExDp1 = Excitatory neuron Deep 1, ExDp2 = Excitatory neuron Deep 2, PgS = Progenitor in S cell cycle phase, PgG2M = Progenitor in G2M cell cycle phase, Mic = Microglia, End = Endothelial cell, Per = Pericyte, EN = Excitatory Neuron, L6 CPN = Layer 6 Cortical Projection Neurons, SPN = subplate neurons, CRN = Cajal-Retzius cells, IN = Interneurons, MSN = Medium Spiny Neurons, AC-f = fibrous Astrocyte, AC-p = protoplasmic Astrocyte, gIPC = glial Intermediate Progenitor Cell, tRG = truncated Radial Glia, EPD = Ependymal cell, RG/AC = Radial Glia / Astrocyte, AC = Astrocyte, gIPC-O = Oligodendrogenesis-biased gIPC, giIPC = glial IPC, preOL = pre Oligodendrocyte, nIPC = neuronal IPC, TAC = Transit-Amplifying Cell/cycling progenitor, mIPC = multipotent IPC, UD = Undefined, MG = Microglia, CP = prenatal Cortical Plate or choroid plexus, BVC = Blood Vessel Cell, cRG = cycling Radial Glia.

**Fig. EV4.**
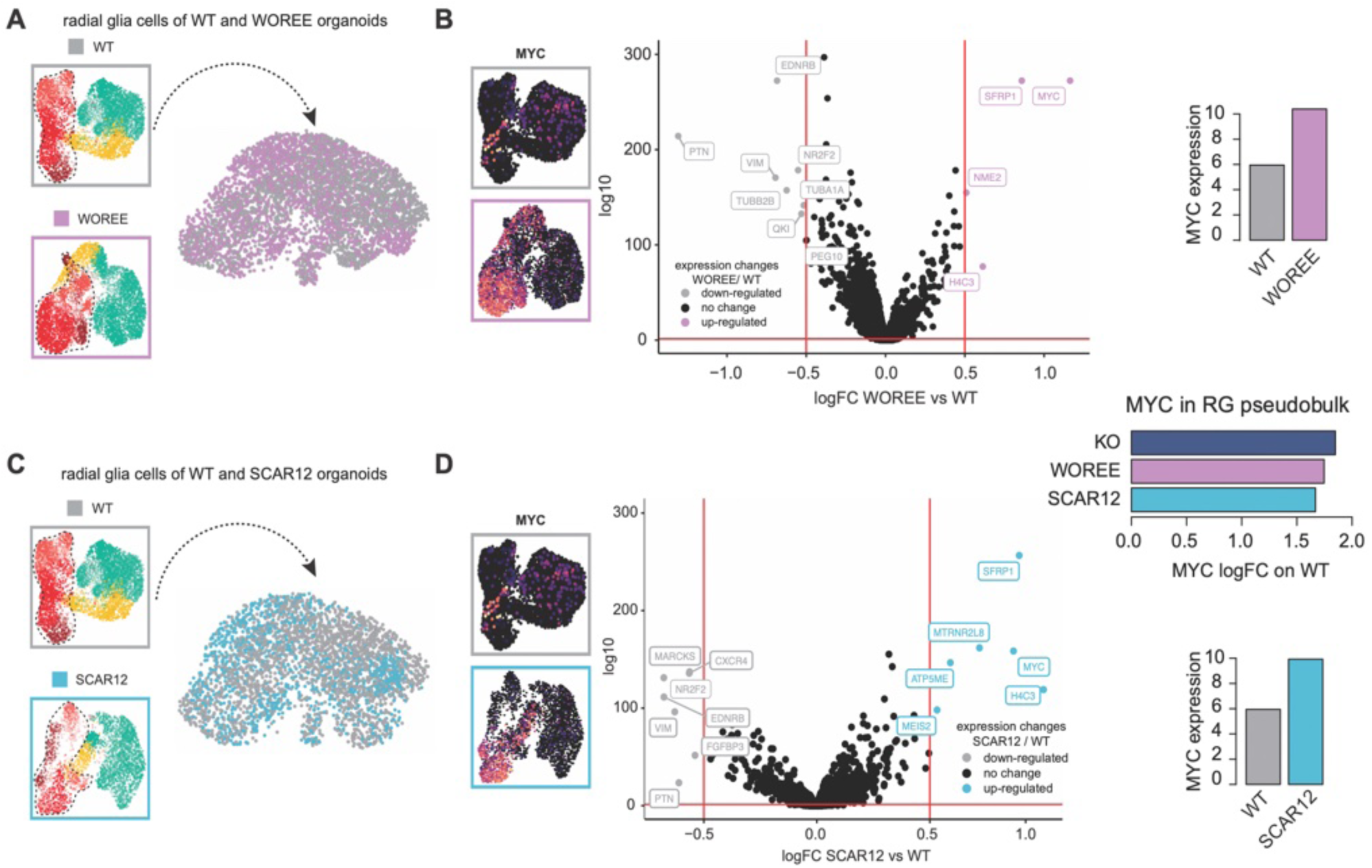
MYC is activated in radial glia of WWOX-deficient neurodevelopmental disorders patients. **(A)** Radial glia UMAP from WT and WOREE organoids (n = 4,712). **(B)** Differential gene expression between WOREE and WT radial glia cells, upregulated genes in WOREE are shown in lilla, downregulated genes in the WOREE in grey, not significant genes are in black. On the right MYC expression in WT and WOREE organoids, quantified in the bar graphs presented below. **(C)** Radial glia isolation from WT and SCAR12 organoids (n = 3,333). **(D)** Differential gene expression between SCAR12 and WT organoids. MYC expression in WT and SCAR12, shown on the right, quantified in the bar graphs presented below. Upregulated genes in SCAR12 are shown in light blue, downregulated genes in the SCAR12 in grey, not significant genes are in black. Red bars show threshold of significance, -log 10 p-value > 1.30 (p-value = 0.05) and log2 fold change of ± 0.5.

**Fig. EV5.**
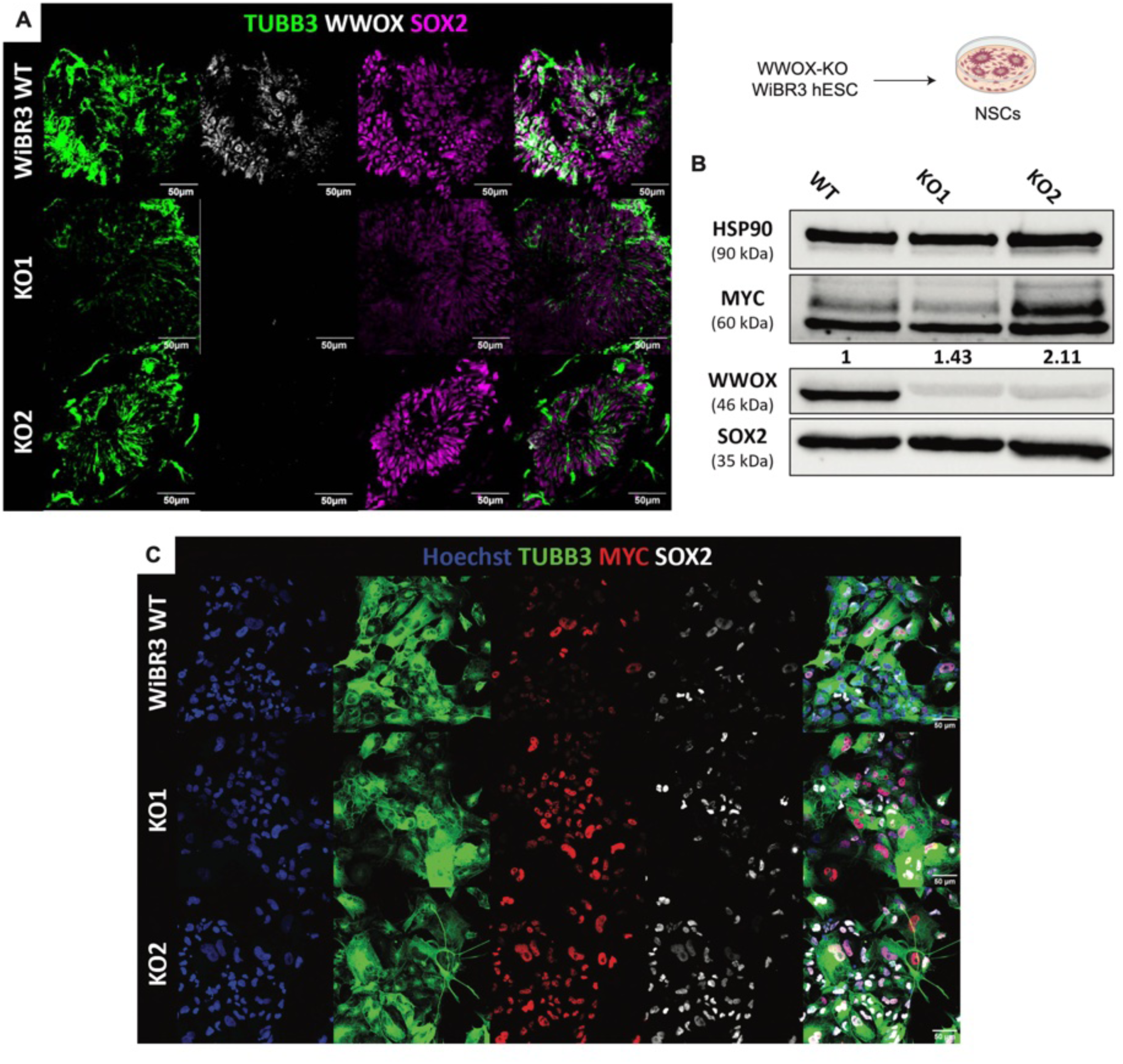
WWOX-KO leads to MYC activation in radial glia cells in-vitro. **(A)** Immunofluorescent staining of neural stem cells (NSCs) generated from the human embryonic stem cell line WiBR3. **(B)** Immunoblot analysis of lysates from 2D-cultured NSCs, comparing the MYC expression levels following WWOX-KO. The numbers indicate a quantification of the bands, normalized to the bands of HSP90 and presented as fold change. **(C)** Representative fields from immunostaining for MYC, SOX2 and TUBB3 in the NSCs culture.

**Fig. EV6.**
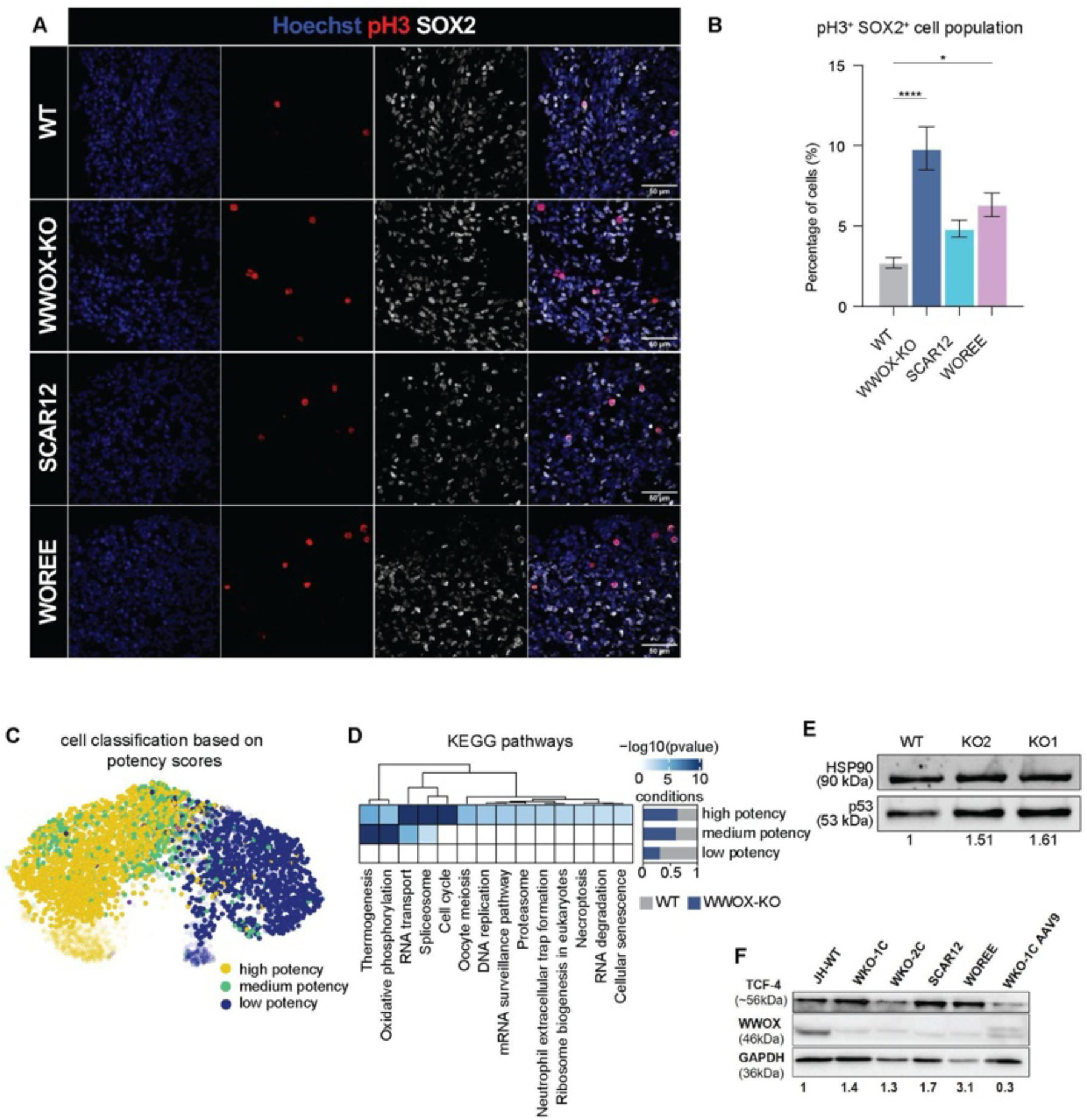
WWOX-KO leads to MYC activation in radial glia cells in-vitro. **(A)** Representative images of immunostained week 16 cerebral organoids for SOX2 and the G2/M marker pH3. **(B)** Quantification of images in (E) comparing WWOX-KO, SCAR12 and WOREE To wildtype levels. See Table EV1 for details on organoid numbers, sections, and batches. Data are presented as mean ± SEM. Statistical significance was determined using one-way ANOVA with Tukey’s multiple comparisons test. n.s (non-significant), *p ≤ 0.05, **p ≤ 0.01, ***p ≤ 0.001, ****p ≤ 0.0001. **©** UMAP representation of the potency states assigned to each single cell, assigned based on the score plotted in main Figure 4E. **(D)** Pathway analysis of pluripotency states. Shown top significant pathways after Bonferroni correction (p-value < 0.05). **(E)** Immunoblot analysis of WT and WWOX-KO NSCs showing p53 activation after loss of WWOX. **(F)** Immunoblot analysis of TCF4 protein levels in week 16 cerebral organoids comparing WWOX-KO, SCAR12, WOREEand WKO-AAV9 rescued lines to wildtype levels.

**Fig. EV7.**
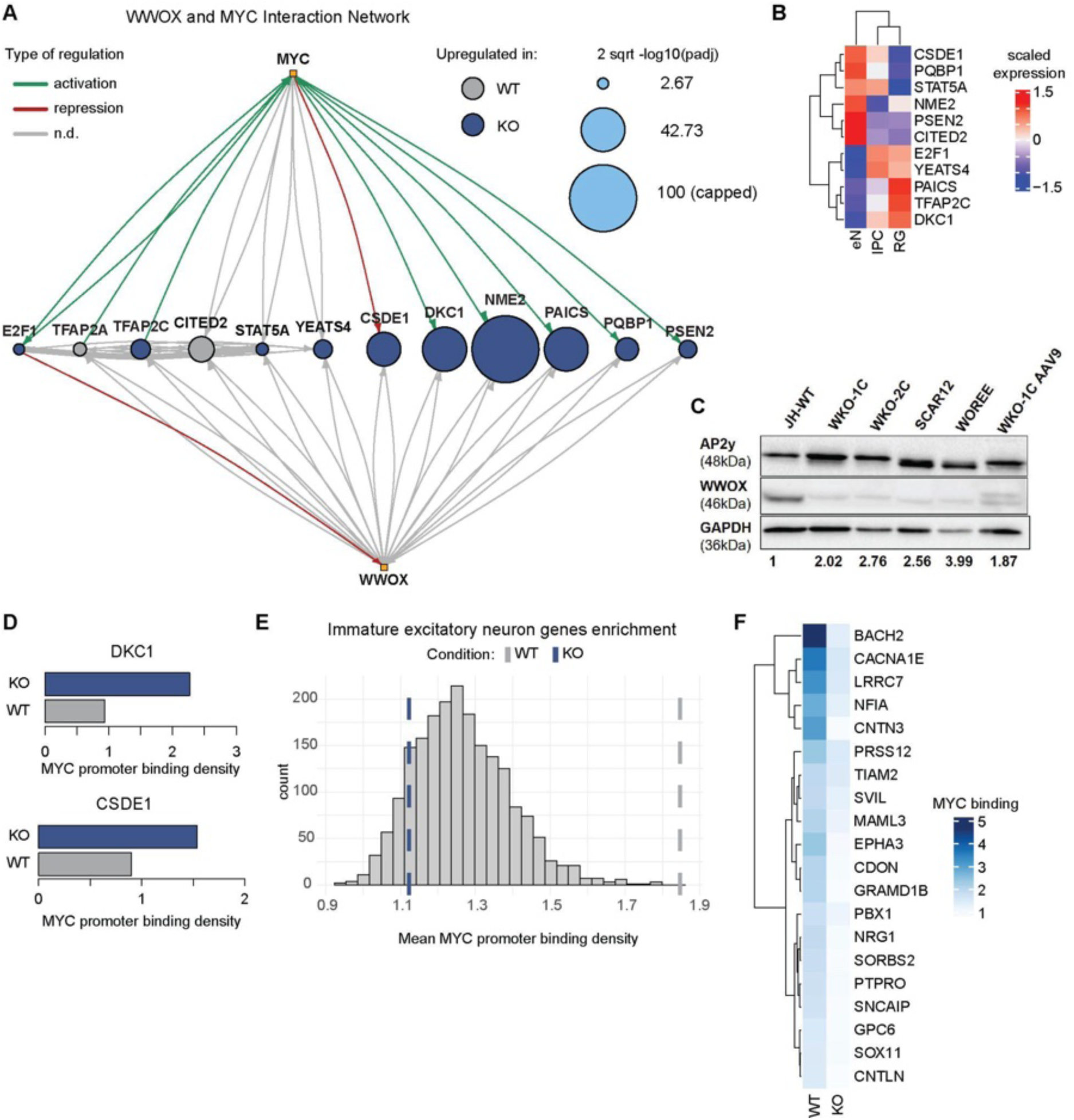
Shared MYC and WWOX interactome analysis. **(A)** MYC and WWOX interaction network, including only shared genes differentially expressed in RGs. The nodes represent single genes, whose colors depict the condition in which it is upregulated (grey = WT, blue = KO), and the size the square root of the -log10 p-value capped at 100 for visualization purposes. Edges describe the relationship of interactions: green activation, red inhibition, and grey not known. The arrows express the direction of interaction. MYC and WWOX are depicted by yellow squares. **(B)** Expression pattern of interactome-derived genes in FACS-sorted bulk RNA sequencing data. Data are scaled by row. **(C)** Immunoblot validation of upregulated pluripotency marker TFAP2C in week 16 cerebral organoids, comparing WWOX-KO, WOREE, SCAR12, and AAV9 rescued WWOX-KO to wildtype. **(D)** MYC binding density on promoters of DKC1 and CSDE1 in WT and KO conditions in ChIP sequencing data. **(E)** Enrichment analysis of manually curated immature neuron genes in WT and KO ChIP-seq data. Significance was determined by comparing observed enrichment values to those obtained from 1000 iterations with randomly selected gene sets of equal size (WT pval < 0.0001). MYC binding density is shown for each condition. **(F)** MYC binding among the top 20 immature neurons genes in WT condition.

**Fig. EV8.**
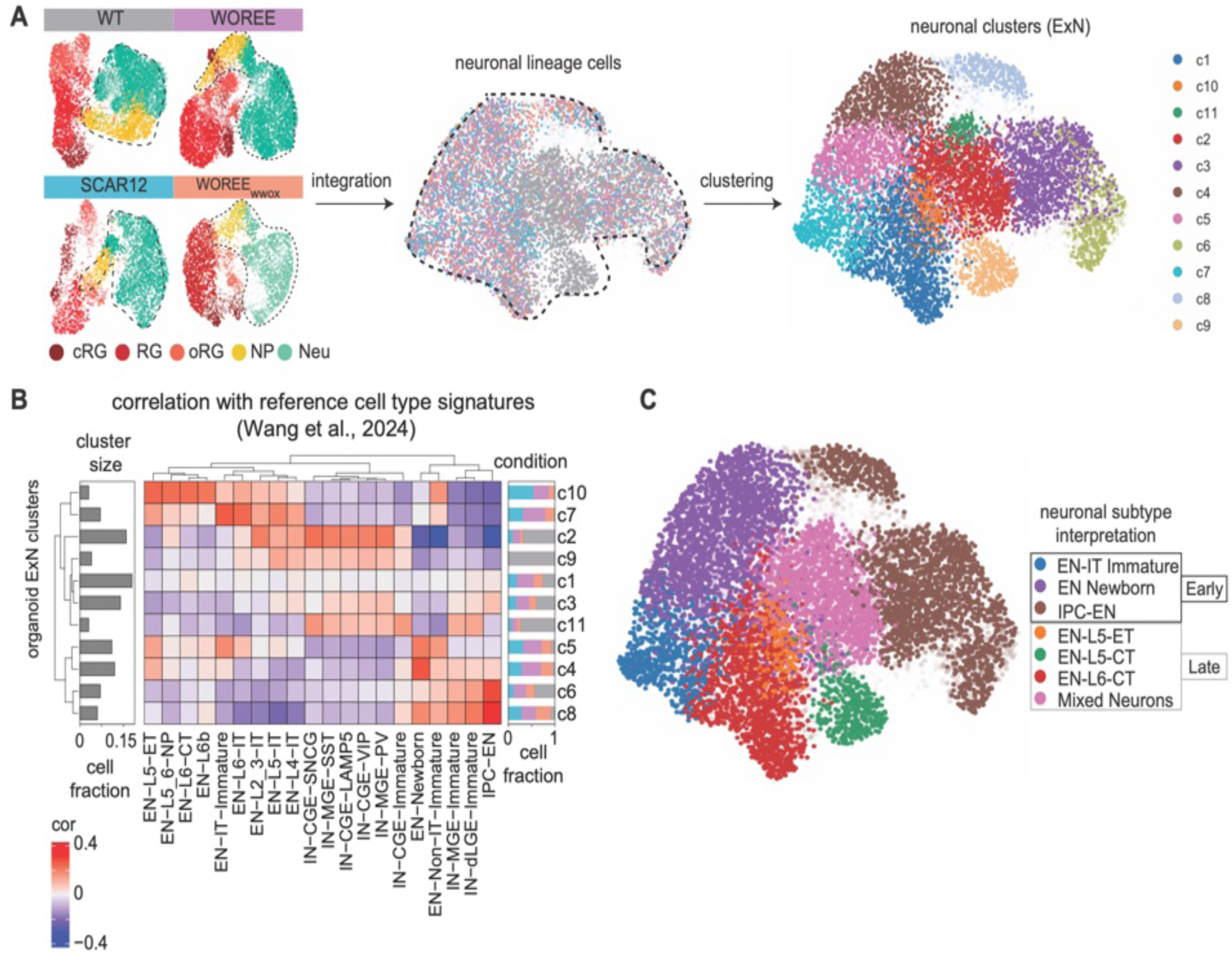
Neuronal cell types annotation. **(A)** (Left) Isolation of neurons from WT, WOREE, SCAR12 and WOREE-WWOX organoids. (Right) UMAP plot coloured by condition. **(B)** Clustering of neuronal cells recovering 11 clusters. **(C)** Heatmap of correlations of cluster signatures to reference neuronal cell types (10.1101/2024.01.16.575956). The barplot on the left shows the size of each cluster, while the stacked barplot on the right represents how many cells populate each cluster divided by condition. **(D)** UMAP visualization coloured by neuronal subtypes recovered from (C). The subtypes are assigned based on best correlation score with reference signatures, and divided in early and late classes based on the age these subtypes are found in the reference dataset. The mixed neuronal population is assigned depending on the multiple high scores with both interneuron and excitatory neuron cell types, thus presenting a more general neuronal signature.

**Fig. EV9.**
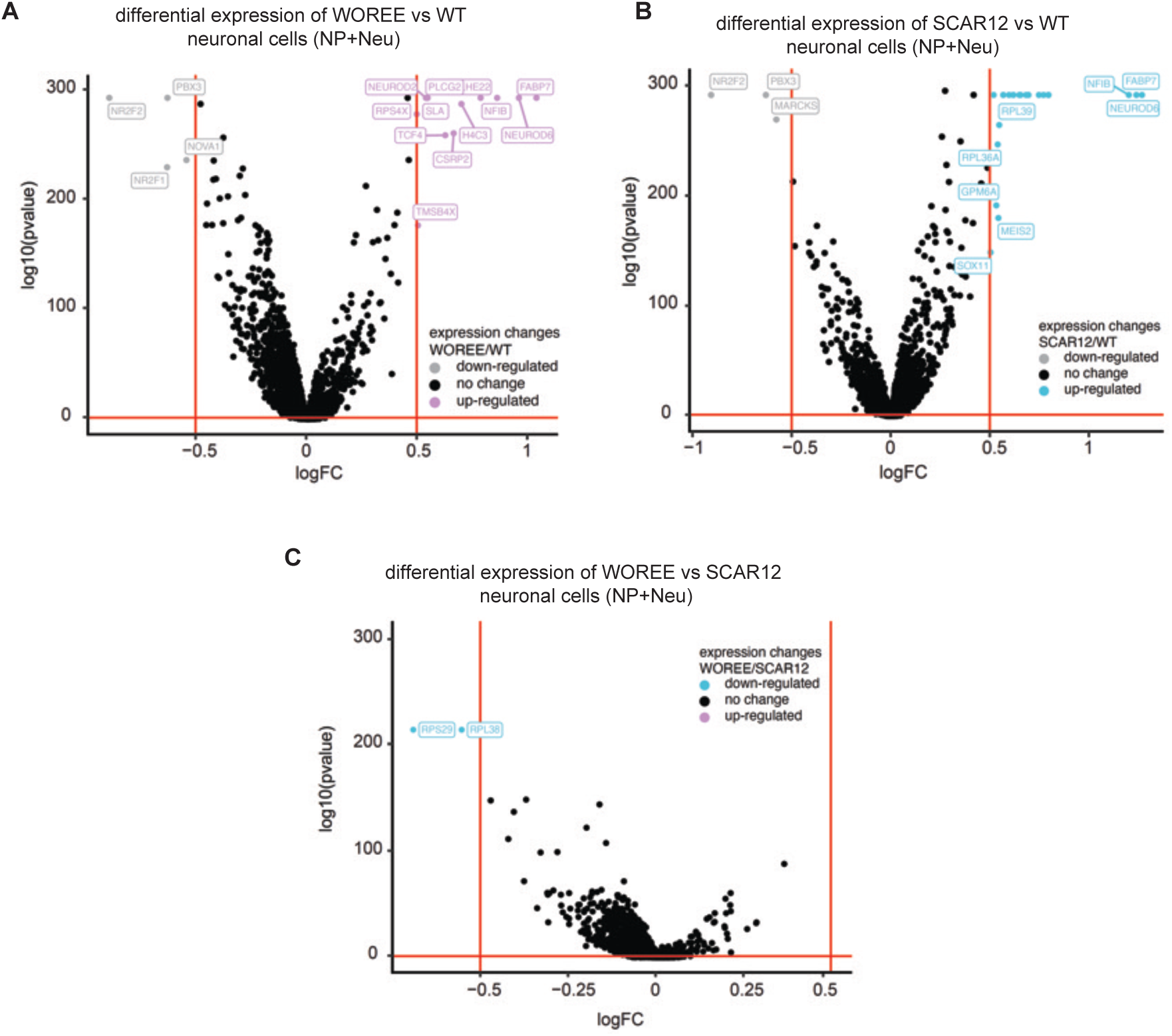
Differentially expressed genes in patient-derived neurons. **(A)** Volcano plot showing differential gene expression between WOREE and WT neurons cells, upregulated genes in WOREE are shown in lilla, downregulated genes in the WOREE in grey, not significant genes are in black. **(B)** Differential gene expression between SCAR12 and WT neurons. Upregulated genes in SCAR12 are shown in light blue, downregulated genes in the SCAR12 in grey, not significant genes are in black. **(C)** Differential gene expression between WOREE and SCAR12 neurons. Upregulated genes in WOREE are shown in lilla, Upregulated genes in SCAR12 are shown in light blue, not significant genes are in black.

**Fig. EV10.**
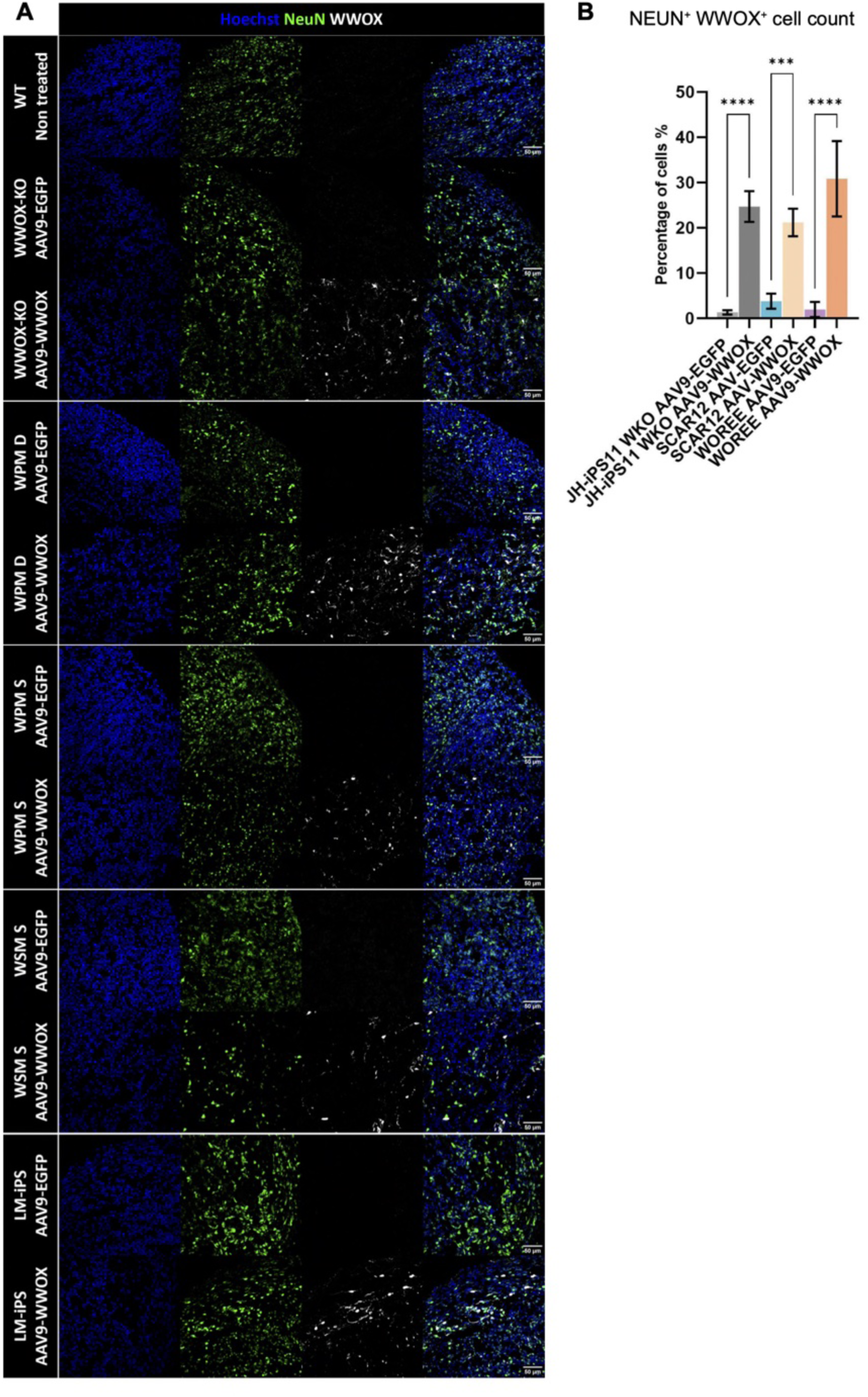
AAV9-WWOX infection in cerebral organoids restores neuronal WWOX expression. **(A)** Immunostaining of week 16 COs infected with AAV9-WWOX showing the expression of the neuronal marker NeuN and WWOX per cell line, validating the neuronal restoration of WWOX to patient-derived organoids through neuronal expression of WWOX. **(B)** Quantification of percentage of NEUN+ WWOX+ cells of images in (A) comparing AAV9-hSyn-hWWOX infected organoids to their respective AAV9-hSyn-eGFP infected mutants. (WT: n= 4 organoids; WPM S AAV9-EGFP: n=4; WPM S AAV9-WWOX: N=4; WPM D AAV9-EGFP: n=4; WPM D AAV9-WWOX: n=4; WSM S AAV9-EGFP: n=4; WSM S AAV9-WWOX: n=4; LM-iPS AAV9-EGFP: n=3; LM-iPS AAV9-WWOX: n=4). *See Table EV1 for details on organoid numbers, sections, and batches. Data are presented as mean ± SEM. Statistical significance was determined using one-way ANOVA with Sidak’s multiple comparisons test. n.s (non-significant), *p ≤ 0.05, **p ≤ 0.01, ***p ≤ 0.001, ****p ≤ 0.0001*.

**Fig. EV11.**
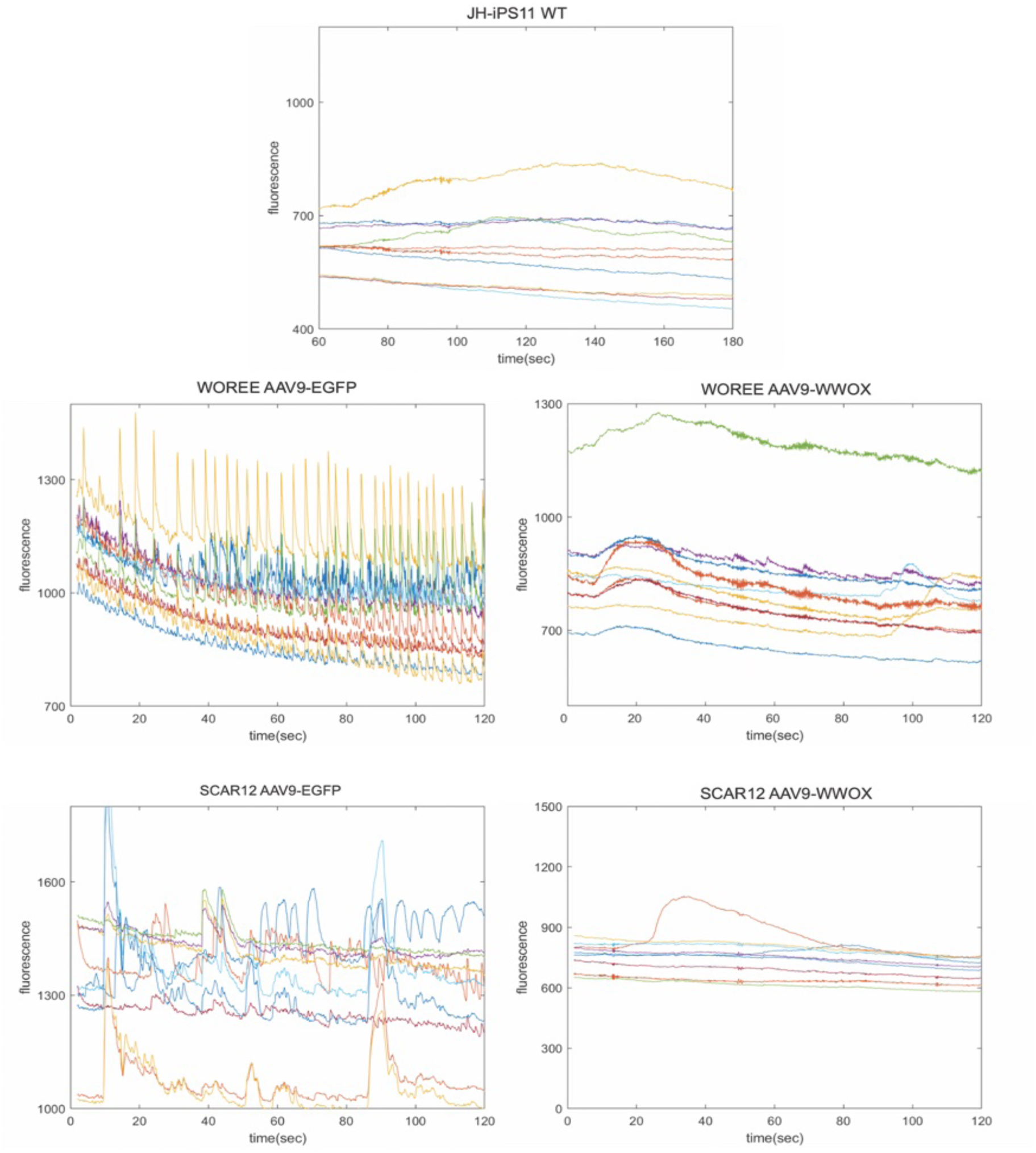
Neuronal restoration of WWOX suppressed neuronal hyperexcitability. Representative calcium transient in selected neurons recorded in week 16 COs infected either with AAV9-EGFP or AAV9-WWOX of each cell line.

**Fig. EV12.**
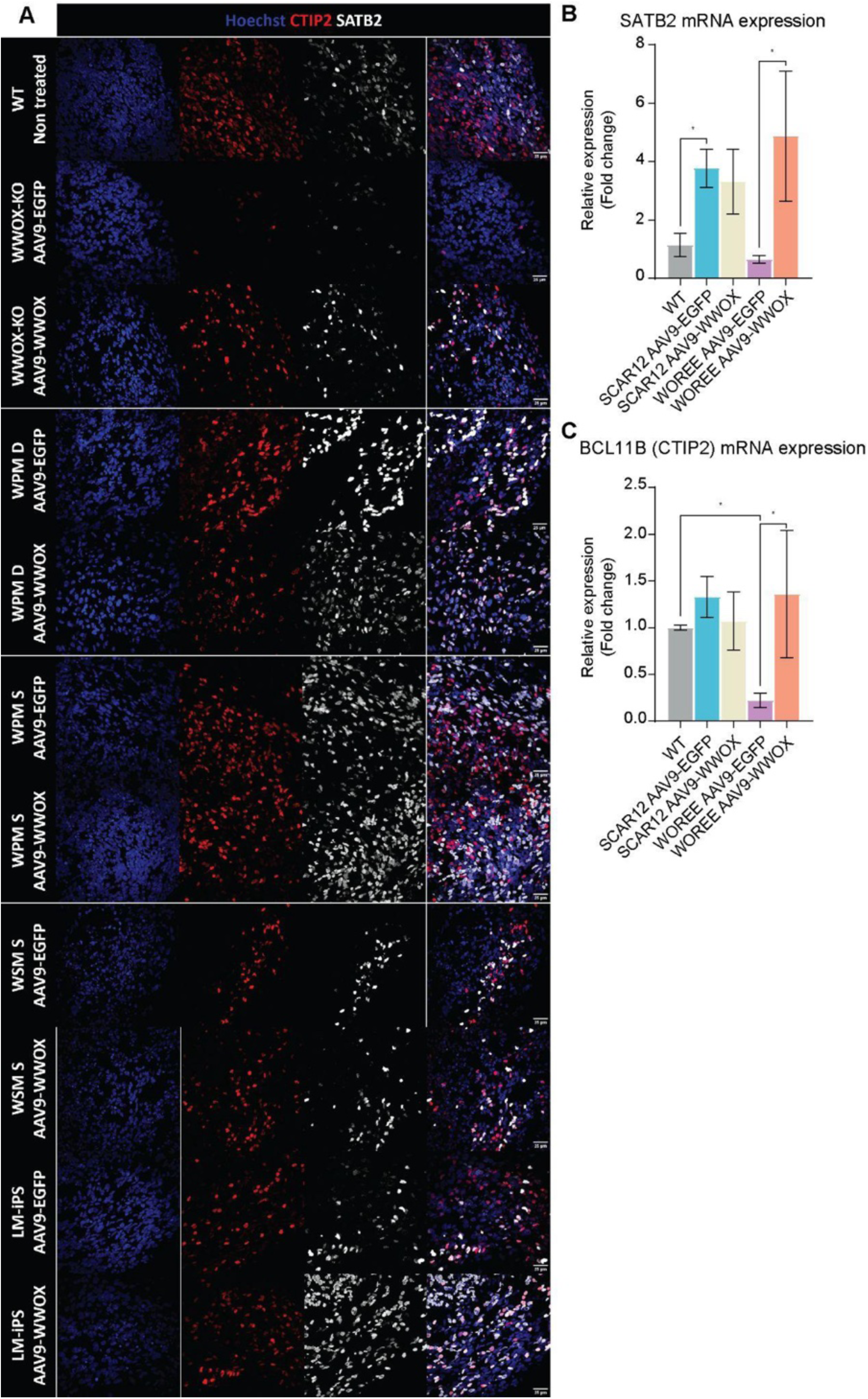
The effect of neuronal WWOX restoration on cortical layers markers. (**A)** Immunostaining of week 16 COs infected with either AAV9-EGFP or AAV9-WWOX showing the expression of the neuronal markers CTIP2 and SATB2, as an expansion of figure 6, showing all lines included (WT: n= 4 organoids; WPM S AAV9-EGFP: n=4; WPM S AAV9-WWOX: N=4; WPM D AAV9-EGFP: n=4; WPM D AAV9-WWOX: n=4; WSM S AAV9-EGFP: n=4; WSM S AAV9-WWOX: n=4; LM-iPS AAV9-EGFP: n=3; LM-iPS AAV9-WWOX: n=4). **(B)** qPCR analysis of the expression of the superficial layer neuron marker SATB2 in week 16 organoids (WT: n=4 organoids; SCAR12 AAV9-EGFP: n=8; SCAR12 AAV9-WWOX: n=8; WOREE AAV9-EGFP: n=6, WOREE AAV9-WWOX: n=6). Statistical significance was determined by a one-way ANOVA test, correcting for multiple comparisons by controlling the FDR using the Benjamini–Hochberg procedure. **(C)** qPCR analysis of the expression of the deep layer neuron marker CTIP2 (BCL11B) in week 16 organoids (WT: n=4 organoids; SCAR12 AAV9-EGFP: n=8; SCAR12 AAV9-WWOX: n=8; WOREE AAV9-EGFP: n=6, WOREE AAV9-WWOX: n=6). Statistical significance was determined by a one-way ANOVA test, correcting for multiple comparisons by controlling the FDR using the Benjamini–Hochberg procedure.

**Fig. EV13.**
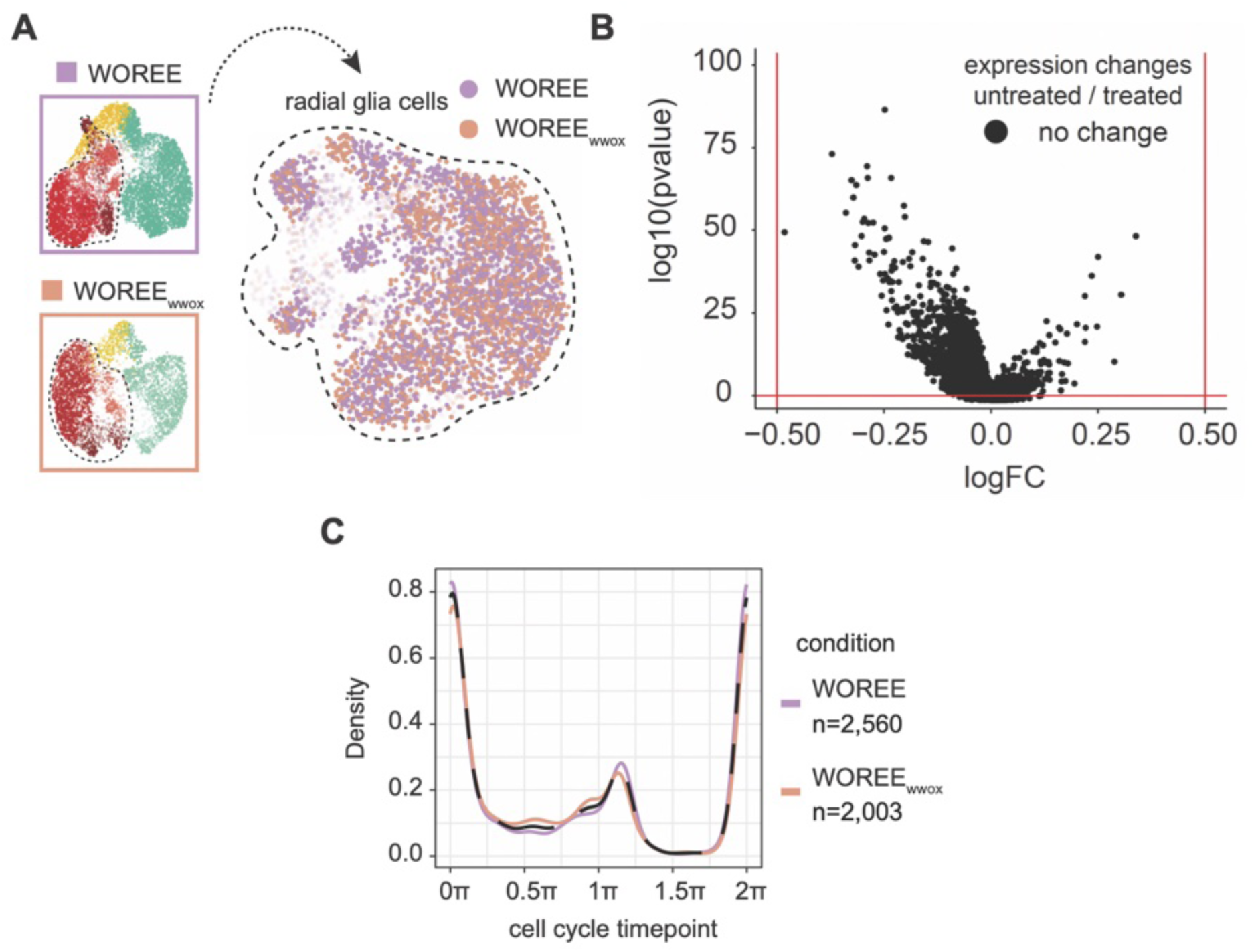
Neuron-targeted gene therapy did not affect the transcriptome and cell-cycle phase of radial glia cells. **(A)** Isolation of radial glia cells from untreated (WOREE) and treated (WOREE-WWOX) organoids. On the right, UMAP cell visualization colored by condition. **(B)** Differential gene expression between radial glia in treated vs untreated organoid shows no significant differentially expressed gene between WOREE-WWOX and WOREE radial glia cells. **(C)** Cell cycle inference in radial glia cells showing overlap of WOREE-WWOX and WOREE conditions.

