## Supplementary material for "WWOX deficiency impairs neurogenesis and neuronal function in human organoids": Expanded View

Expanded View figures for

Daniel J. Steinberg *et al*

*Corresponding authors:

Rami I. Aqeilan, Jose Davila-Velderrain

**This PDF file includes:**

Figs. EV 1 to EV13

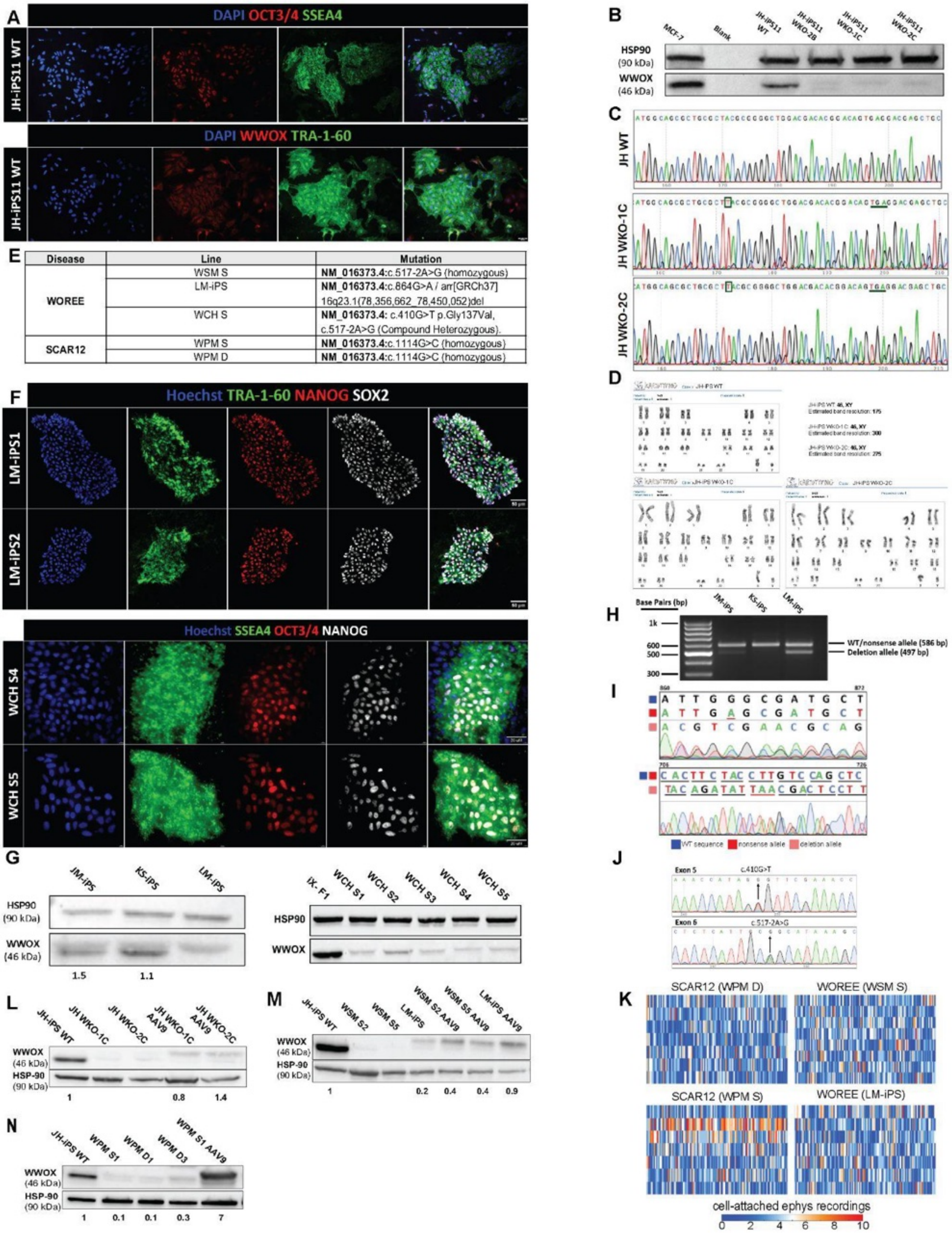
A
DAPI OCT3/4 SSEA4
JH-iPS11 WT
DAPI WWOX TRA-1-60
JH-iPS11 WT
B
MCF-7
Blank
JH-iPS11 WT
JH-iPS11 WKO-2B
JH-iPS11 WKO-1C
JH-iPS11 WKO-2C
HSP90 (90 kDa)
WWOX (46 kDa)
C
JH WT
JH WKO-1C
JH WKO-2C
D
JH-iPS WT 46, XY
Estimated band resolution: 175
JH-iPS WKO-1C: 46, XY
Estimated band resolution: 300
JH-iPS WKO-2C: 46, XY
Estimated band resolution: 275
E
Disease | Line | Mutation
WOREE | WSM S | NM_016373.4:c.517-2A>G (homozygous)
WOREE | LM-iPS | NM_016373.4:c.864G>A / arr[GRCh37] 16q23.1(78,356,662_78,450,052)del
WOREE | WCH S | NM_016373.4: c.410G>T p.Gly137Val, c.517-2A>G (Compound Heterozygous).
SCAR12 | WPM S | NM_016373.4:c.1114G>C (homozygous)
SCAR12 | WPM D | NM_016373.4:c.1114G>C (homozygous)
F
Hoechst TRA-1-60 NANOG SOX2
LM-iPS1
LM-iPS2
Hoechst SSEA4 OCT3/4 NANOG
WCH S4
WCH S5
G
JM-iPS
KS-iPS
LM-iPS
HSP90 (90 kDa)
WWOX (46 kDa)
1.5
1.1
IX-F1
WCH S1
WCH S2
WCH S3
WCH S4
WCH S5
HSP90
WWOX
H
Base Pairs (bp)
JM-iPS
KS-iPS
LM-iPS
1k
600
500
300
WT/nonsense allele (586 bp)
Deletion allele (497 bp)
I
WT sequence
nonsense allele
deletion allele
J
Exon 5
c.410G>T
Exon 6
c.517-2A>G
K
SCAR12 (WPM D)
WOREE (WSM S)
SCAR12 (WPM S)
WOREE (LM-iPS)
cell-attached ephys recordings
0 2 4 6 8 10
L
JH-iPS WT
JH WKO-1C
JH WKO-2C
JH WKO-1C AAV9
JH WKO-2C AAV9
WWOX (46 kDa)
HSP-90 (90 kDa)
1
0.8
1.4
M
JH-iPS WT
WSM S2
WSM S5
LM-iPS
WSM S2 AAV9
WSM S5 AAV9
LM-iPS AAV9
WWOX (46 kDa)
HSP-90 (90 kDa)
1
0.2
0.4
0.4
0.9
N
JH-iPS WT
WPM S1
WPM D1
WPM D3
WPM S1 AAV9
WWOX (46 kDa)
HSP-90 (90 kDa)
1
0.1
0.1
0.3
7

**Fig. EV1. iPSCs generation and CRISPR/Cas9 editing. (A)** Immunofluorescent staining of pluripotency markers in WT iPSC line established JH-iPS11, and for WWOX. **(B)** Immunoblot analysis confirming the successful knock out of the WWOX gene in JH-iPS11 cells. WKO; WWOX knockout, followed by a clone-specific label. **(C)** Sanger sequencing of JH-WT, JH WKO-1C and JH WKO-2C showing indels (boxed) introducing a premature stop codon (horizontal line). **(D)** G-band karyotyping reports of JH-WT, JH WKO-1C and JH WKO-2C indicating no detectable chromosomal abnormalities. **(E)** Summary of WOREE and SCAR12 patients included in the study and their exact mutations. **(F)** Immunofluorescent staining of pluripotency markers in a WOREE patient-derived iPSCs lines, LM-iPS and WCH, harboring different complex heterozygous mutation in the WWOX gene. **(G)** Immunoblot analysis confirming WWOX expression levels in iPSCs of the LM-iPS patient and the healthy parents (JM-iPS and KS-iPS) (left panel), and in WCH patient iPSC clones compared to a wildtype control***. (H)** Gel electrophoresis after PCR amplification of a cDNA fragment spanning parts of exons 5-8 of the WWOX gene, showcasing the absence of exon 6 in one parental allele. **(I)** Sanger sequencing of the cDNA fragments shown in (E). The top panel shows the maternal nonsense mutation (c.864G>A) compared to a reference genome, and the sequence shift in the paternal allele due to the deletion of 0.9 Mbp. The bottom panel shows the premature stop codon (underlined in red) resulting from the deletion and frameshift in the paternal allele. The numbers in the top corners denote the position in the reference WWOX coding sequence. **(J)** Sanger sequencing of WCH S4 clone showing the maternal missense mutation in exon 5 (c.410G>T), and the paternal splice site mutation in in exon 6 (c.517-2A>G). **(K)** Raster plots demonstrating selected neuronal firing from SCAR12 (WPM D and WPM S) and WOREE (WSM S and LM-iPS) week 7 organoids over 4 minutes of cell-attached recording, related to Figure 1C. **(L)** Immunoblot analysis of JH-WT, JH-WKO and JH-WKO AAV9-hSyn-hWWOX cerebral organoids from week 16, confirming knockout levels of WWOX and its rescue. **(M)** Immunoblot analysis of WWOX levels in WOREE lines and their AAV9 rescue included in the scRNA-seq experiment in week 16 cerebral organoids. **(N)** Immunoblot analysis of WWOX levels in SCAR12 lines included in the sc-RNA-seq experiment in week 16 cerebral organoids.

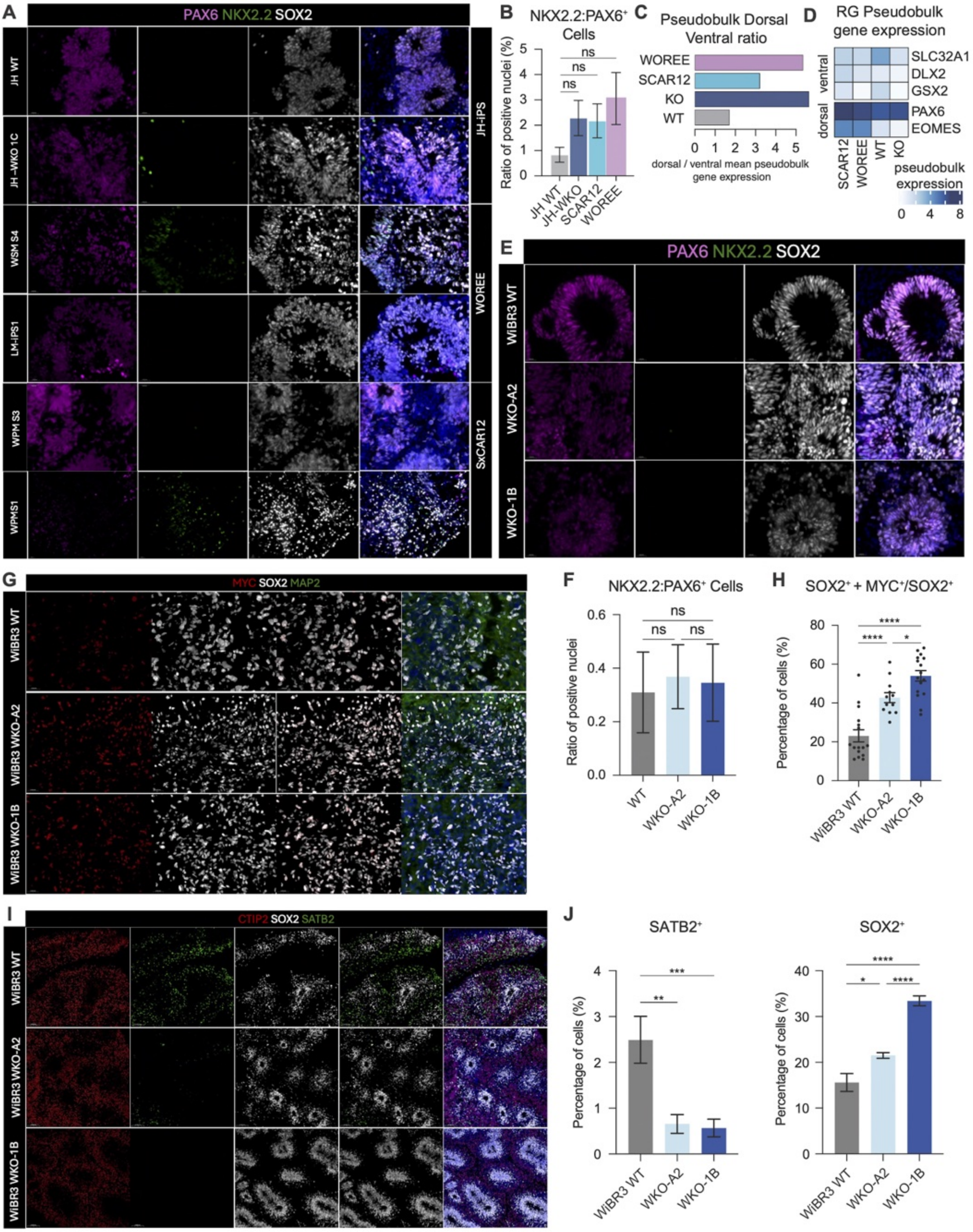
A
PAX6 NKX2.2 SOX2
JH WT
JH -WKO 1C
WSM S4
LM-iPS1
WPM S3
WPMS1
JH-iPS
WOREE
SxCAR12
B
NKX2.2:PAX6+ Cells
Ratio of positive nuclei (%)
ns
JH WT
JH-WKO
SCAR12
WOREE
C
Pseudobulk Dorsal Ventral ratio
WOREE
SCAR12
KO
WT
dorsal / ventral mean pseudobulk gene expression
D
RG Pseudobulk gene expression
SLC32A1
DLX2
GSX2
PAX6
EOMES
ventral
dorsal
SCAR12
WOREE
WT
KO
pseudobulk expression
E
PAX6 NKX2.2 SOX2
WiBR3 WT
WKO-A2
WKO-1B
G
MYC SOX2 MAP2
WiBR3 WT
WiBR3 WKO-A2
WiBR3 WKO-1B
F
NKX2.2:PAX6+ Cells
Ratio of positive nuclei
ns
WT
WKO-A2
WKO-1B
H
SOX2+ + MYC+/SOX2+
Percentage of cells (%)
****
*
WiBR3 WT
WKO-A2
WKO-1B
I
CTIP2 SOX2 SATB2
WiBR3 WT
WiBR3 WKO-A2
WiBR3 WKO-1B
J
SATB2+
SOX2+
Percentage of cells (%)
***
**
****
*
WiBR3 WT
WKO-A2
WKO-1B

**Fig. EV2. Characterization and validation of cerebral organoids. (A)** Representative immunofluorescent staining of Pax6 (dorsal marker) and NKX2.2 (ventral marker) in all organoids included in the scRNA seq study at week 7 in vitro. **(B)** Quantification of (a) showing the ratio of NKX2.2: PAX6 positive cells revealing predominant dorsal origin with no significant differences in patterning among lines. **(C)** Analysis of cerebral organoids included in our scRNA-seq data showing increased proportion of dorsal relative to ventral markers expressed in all lines. **(D)** Pseudobulk analysis expression levels of known dorsal and ventral markers in all lines. **(E)** Representative immunofluorescent staining of Pax6 (dorsal marker) and NKX2.2 (ventral marker) of hESCs-derived cerebral organoids of WT and WWOX-KO at week 15 in vitro. **(F)** Quantification of (e) showing the ratio of NKX2.2: PAX6 positive cells revealing predominant dorsal origin with no significant differences in patterning among lines. **(G)** Immunostaining of week 15 hESCs-derived cerebral organoids at week 15 in vitro for SOX2 and MYC double positive populations. **(H)** Quantification of immunostaining in (g) showing expansion of $MYC^{+}$ $SOX2^{+}$ progenitor populations. **(I)** Immunostaining of cerebral organoids at week 15 for SOX2 marker for progenitors, and SATB2 and CTIP2 neuronal markers. **(J)** Quantification of SOX2+ and SATB2+ populations in (I) showing similar reduced differentiation capacities to our iPSC-derived cerebral organoids.

*See Table EV1 for details on organoid numbers, sections, and batches. Data are presented as mean ± SEM. Statistical significance was determined using one-way ANOVA with Tukey's multiple comparisons test. n.s (non-significant), *p ≤ 0.05, **p ≤ 0.01, ***p ≤ 0.001, ****p ≤ 0.0001.*

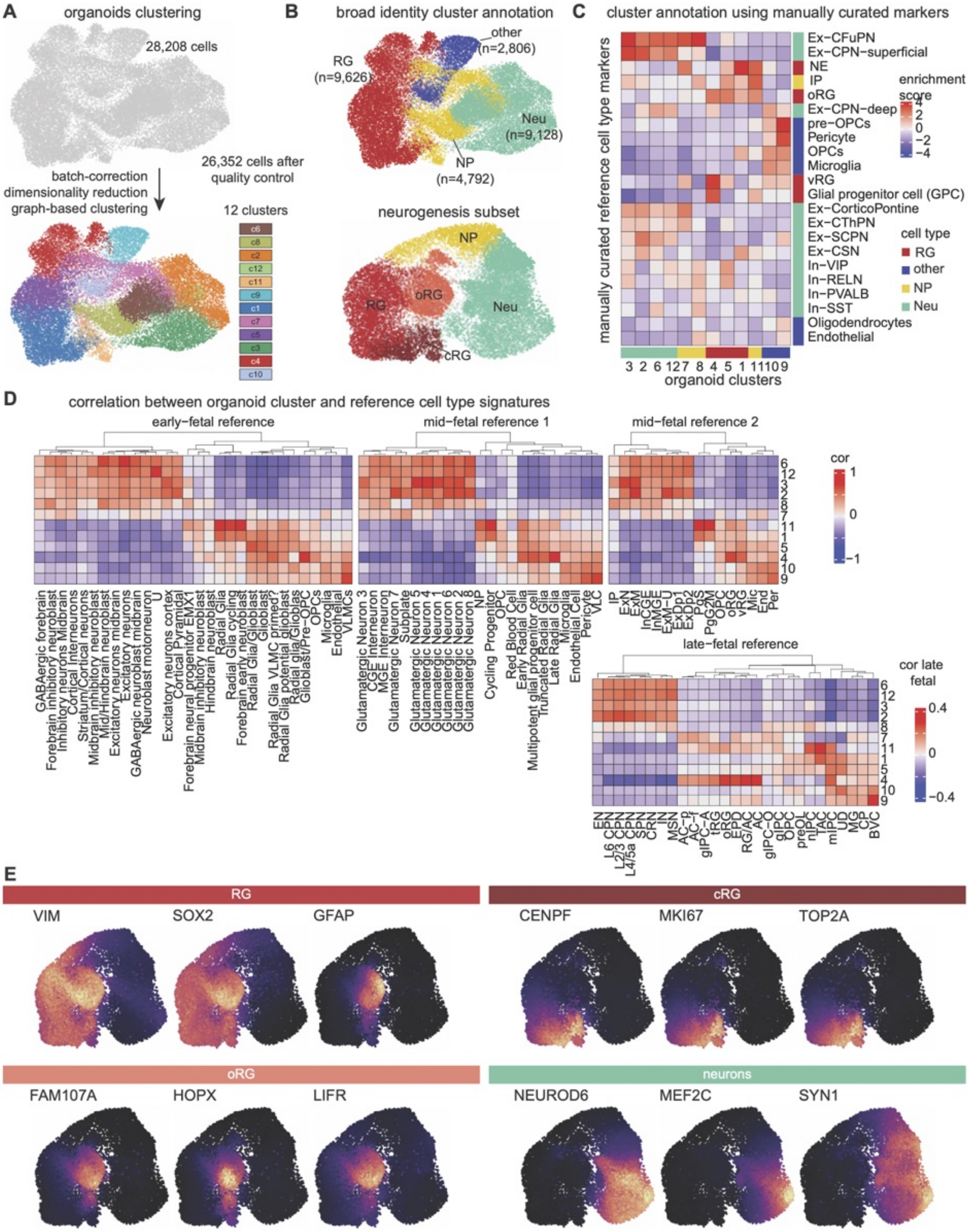

A
organoids clustering
28,208 cells
26,352 cells after quality control
batch-correction
dimensionality reduction
graph-based clustering
12 clusters
B
broad identity cluster annotation
RG (n=9,626)
other (n=2,806)
Neu (n=9,128)
NP (n=4,792)
neurogenesis subset
NP
RG
oRG
Neu
cRG
C
cluster annotation using manually curated markers
manually curated reference cell type markers
Ex-CFuPN
Ex-CPN-superficial
NE
IP
oRG
Ex-CPN-deep
pre-OPCs
Pericyte
OPCs
Microglia
vRG
Glial progenitor cell (GPC)
Ex-CorticoPontine
Ex-CThPN
Ex-SCPN
Ex-CSN
In-VIP
In-RELN
In-PVALB
In-SST
Oligodendrocytes
Endothelial
enrichment score
cell type
RG
other
NP
Neu
organoid clusters
D
correlation between organoid cluster and reference cell type signatures
early-fetal reference
mid-fetal reference 1
mid-fetal reference 2
late-fetal reference
cor
cor late fetal
E
RG
VIM
SOX2
GFAP
cRG
CENPF
MKI67
TOP2A
oRG
FAM107A
HOPX
LIFR
neurons
NEUROD6
MEF2C
SYN1

**Fig. EV3. Cell type identification of scRNA-seq samples. (A)** (Upper) UMAP of all organoids together after standard quality controls (removed 1856 low quality cells). (Lower) Clustering of cells recovering 12 groups. **(B)** Clusters colored by their general annotation, with RG accounting for 9,626 cells, NP 4,792 cells, Neu 9,128 cells and other 2,806 cells. **(C)** Heatmap of scaled enrichment score of manually curated marker genes (rows) in the 12 clusters (columns), all colored by their general cell type annotation. **(D)** Cluster signatures comparison to references fetal cortical development of different ages: early fetal (post conceptional week 08-10), mid-fetal 1 (w16-24), mid-fetal 2 (w15-16), late-fetal (w36). Correlation for early and mid-fetal from -1 to 1, correlation values for late-fetal from -0.4 to 0.4. **(E)** Feature plots in WT organoids for pan-RG, cRG, oRG and Neurons markers. References: early fetal (10.1016/j.devcel.2022.04.016), mid-fetal 1 (10.1016/j.cell.2021.07.039), mid-fetal 2 (10.1016/j.neuron.2019.06.011) and late-fetal (10.1038/s41467-022-34975-2, subject at w36). RG = Radial Glia, vRG = ventricular Radial Glia, oRG = outer Radial Glia, IP = Intermediate Progenitor, Ex-CFuPN = CorticoFugal Projection Neuron, Ex-CThPN = CorticoThalamic Projection Neuron, Ex-SCPN = SubCerebral Projection Neuron, Ex-CPN = Commisural (callosal) Projection Neuron, In = Interneuron, OPC = Oligodendrocyte Progenitor Cell, NE = Neuroepithelial cells, U = Unknown, VLMC = Vascular and Leptomeninges, CGE = Caudal Ganglionic Eminence, MGE = Medial Ganglionic Eminence, ExN = Excitatory Neuron, ExM = Migrating Excitatory neuron, ExM-U = Upper-Migrating Excitatory neuron, ExDp1 = Excitatory neuron Deep 1, ExDp2 = Excitatory neuron Deep 2, PgS = Progenitor in S cell cycle phase, PgG2M = Progenitor in G2M cell cycle phase, Mic = Microglia, End = Endothelial cell, Per = Pericyte, EN = Excitatory Neuron, L6 CPN = Layer 6 Cortical Projection Neurons, SPN = subplate neurons, CRN = Cajal-Retzius cells, IN = Interneurons, MSN = Medium Spiny Neurons, AC-f = fibrous Astrocyte, AC-p = protoplasmic Astrocyte, gIPC = glial Intermediate Progenitor Cell, tRG = truncated Radial Glia, EPD = Ependymal cell, RG/AC = Radial Glia / Astrocyte, AC = Astrocyte, gIPC-O = Oligodendrogenesis-biased gIPC, giIPC = glial IPC, preOL = pre Oligodendrocyte, nIPC = neuronal IPC, TAC = Transit-Amplifying Cell/cycling progenitor, mIPC = multipotent IPC, UD = Undefined, MG = Microglia, CP = prenatal Cortical Plate or choroid plexus, BVC = Blood Vessel Cell, cRG = cycling Radial Glia.

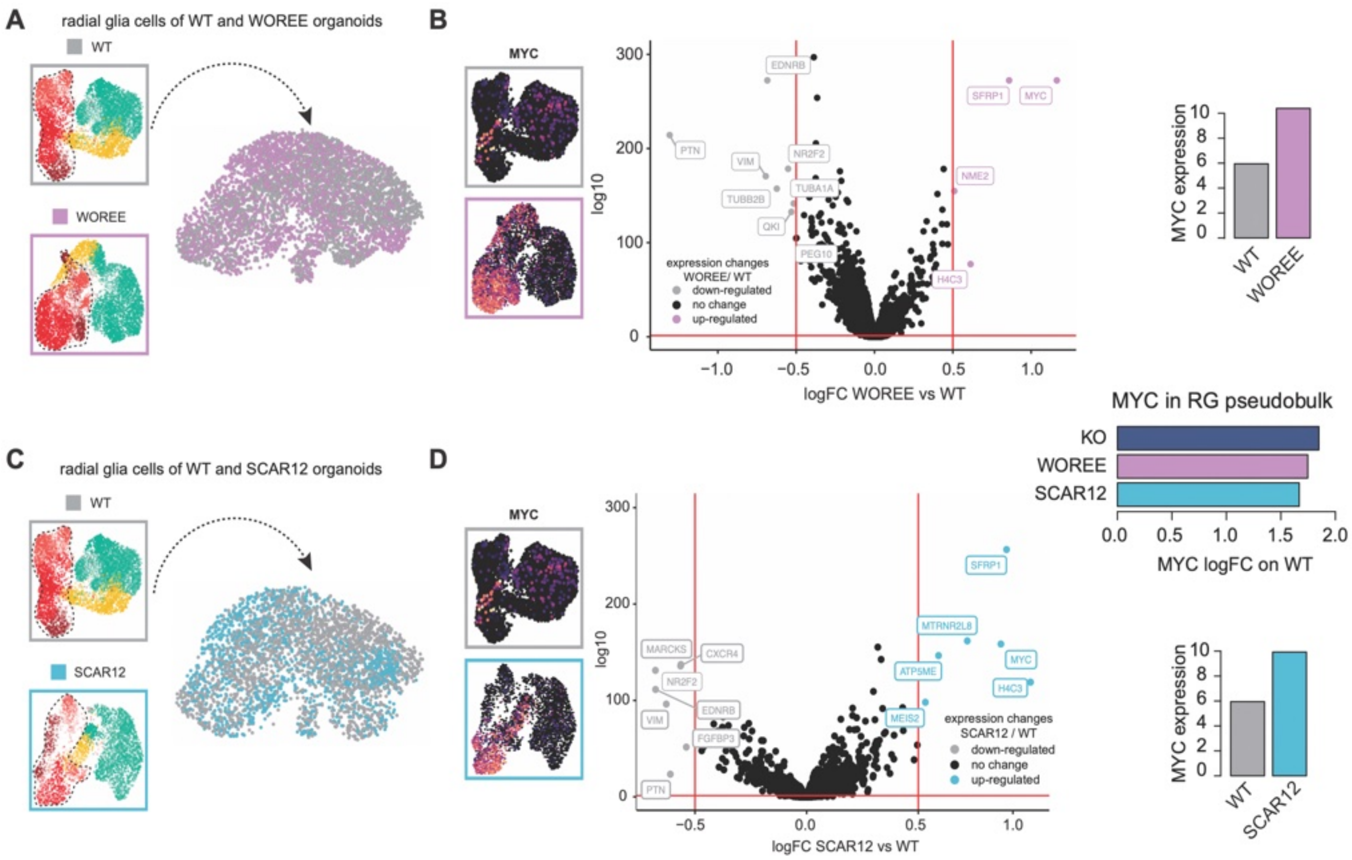


Differential gene expression between WOREE and WT radial glia cells, upregulated genes in WOREE are shown in lilla, downregulated genes in the WOREE in grey, not significant genes are in black. On the right MYC expression in WT and WOREE organoids, quantified in the bar graphs presented below. **(C)** Radial glia isolation from WT and SCAR12 organoids (n = 3,333). **(D)** Differential gene expression between SCAR12 and WT organoids. MYC expression in WT and SCAR12, shown on the right, quantified in the bar graphs presented below. Upregulated genes in SCAR12 are shown in light blue, downregulated genes in the SCAR12 in grey, not significant genes are in black. Red bars show threshold of significance, -log 10 p-value > 1.30 (p-value = 0.05) and log2 fold change of ± 0.5.

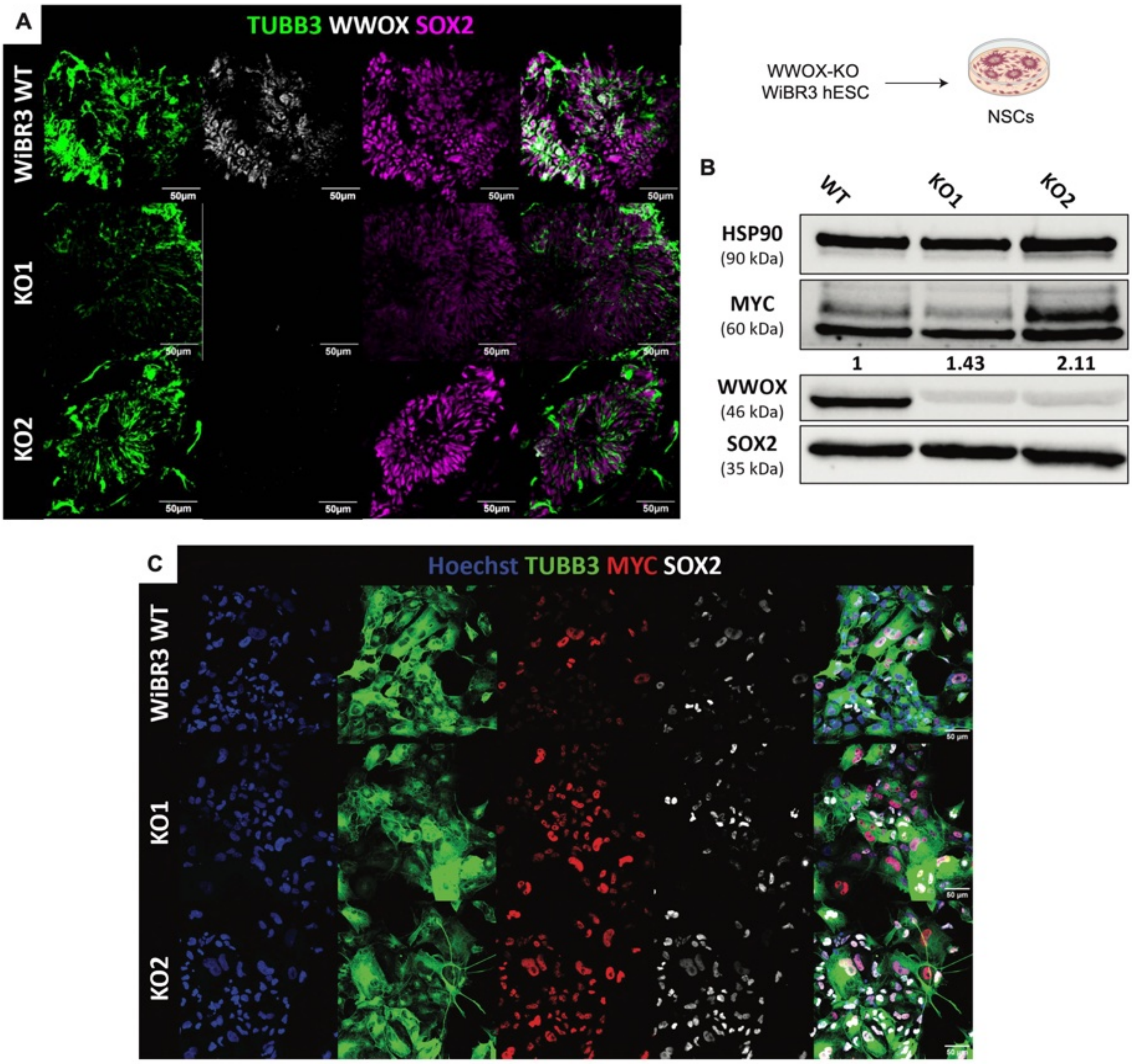


**Fig. EV5. WWOX-KO leads to MYC activation in radial glia cells in-vitro. (A)** Immunofluorescent staining of neural stem cells (NSCs) generated from the human embryonic stem cell line WiBR3. **(B)** Immunoblot analysis of lysates from 2D-cultured NSCs, comparing the MYC expression levels following WWOX-KO. The numbers indicate a quantification of the bands, normalized to the bands of HSP90 and presented as fold change. **(C)** Representative fields from immunostaining for MYC, SOX2 and TUBB3 in the NSCs culture.

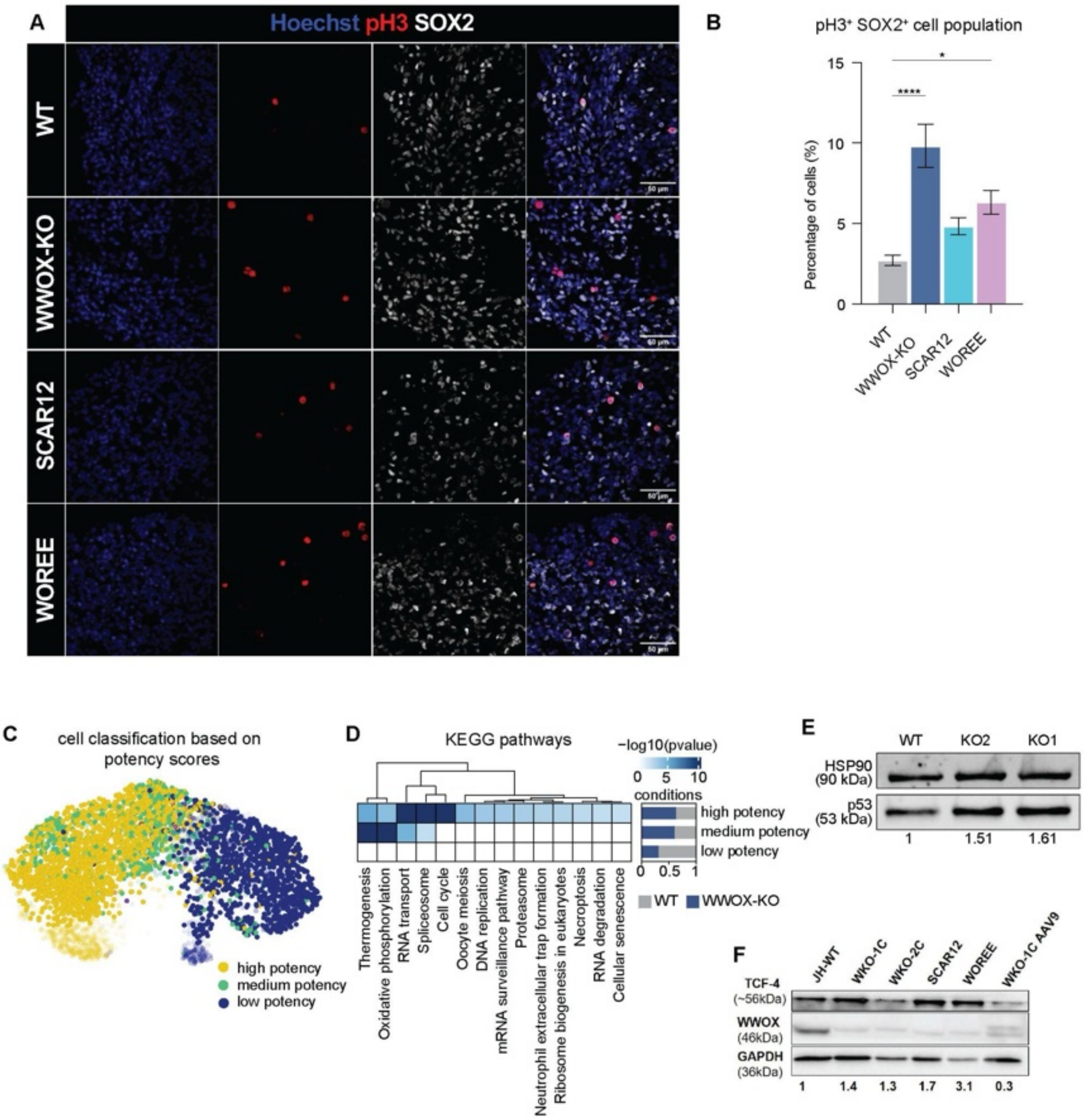


**Fig. EV6. WWOX-KO leads to MYC activation in radial glia cells in-vitro. (A)** Representative images of immunostained week 16 cerebral organoids for SOX2 and the G2/M marker pH3. **(B)** Quantification of images in (E) comparing WWOX-KO, SCAR12 and WOREE To wildtype levels. See Table EV1 for details on organoid numbers, sections, and batches. Data are presented as mean ± SEM. Statistical significance was determined using one-way ANOVA with Tukey's multiple comparisons test. n.s (non-significant), *p ≤ 0.05, **p ≤ 0.01, ***p ≤ 0.001, ****p ≤ 0.0001. © UMAP representation of the potency states assigned to each single cell, assigned based on the score plotted in main Figure 4E. **(D)** Pathway analysis of pluripotency states. Shown top significant pathways after Bonferroni correction (p-value < 0.05). **(E)** Immunoblot analysis of WT and WWOX-KO NSCs showing p53 activation after loss of WWOX. **(F)** Immunoblot analysis of TCF4 protein levels in week 16 cerebral organoids comparing WWOX-KO, SCAR12, WOREEand WKO-AAV9 rescued lines to wildtype levels.

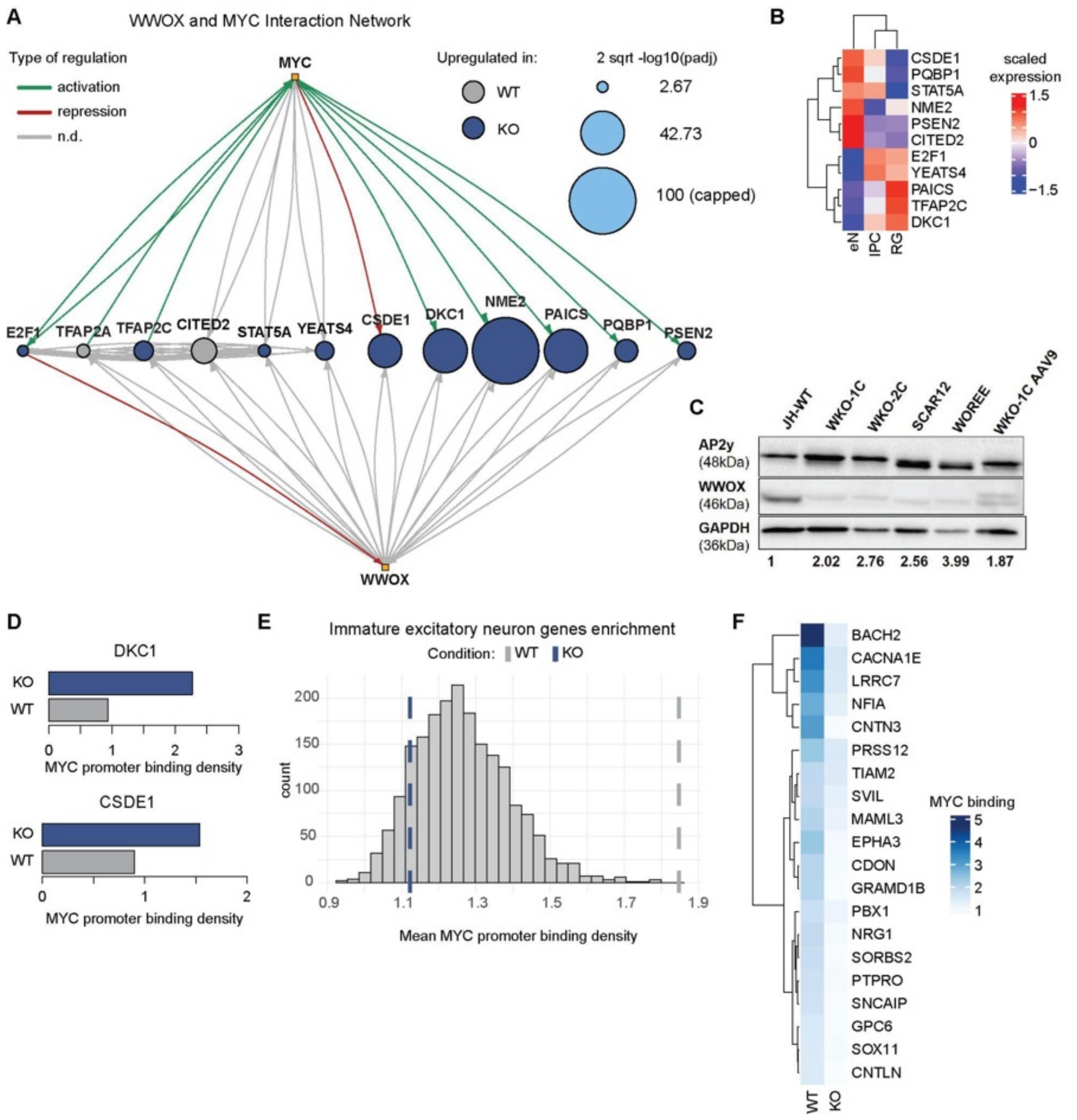


**Fig. EV7. Shared MYC and WWOX interactome analysis. (A)** MYC and WWOX interaction network, including only shared genes differentially expressed in RGs. The nodes represent single genes, whose colors depict the condition in which it is upregulated (grey = WT, blue = KO), and the size the square root of the -log10 p-value capped at 100 for visualization purposes. Edges describe the relationship of interactions: green activation, red inhibition, and grey not known. The arrows express the direction of interaction. MYC and WWOX are depicted by yellow squares. **(B)** Expression pattern of interactome-derived genes in FACS-sorted bulk RNA sequencing data. Data are scaled by row. **(C)** Immunoblot validation of upregulated pluripotency marker TFAP2C in week 16 cerebral organoids, comparing WWOX-KO, WOREE, SCAR12, and AAV9 rescued WWOX-KO to wildtype. **(D)** MYC binding density on promoters of DKC1 and CSDE1 in WT and KO conditions in ChIP sequencing data. **(E)** Enrichment analysis of manually curated immature neuron genes in WT and KO ChIP-seq data. Significance was determined by comparing observed enrichment values to those obtained from 1000 iterations with randomly selected gene sets of equal size (WT pval < 0.0001). MYC binding density is shown for each condition. **(F)** MYC binding among the top 20 immature neurons genes in WT condition.

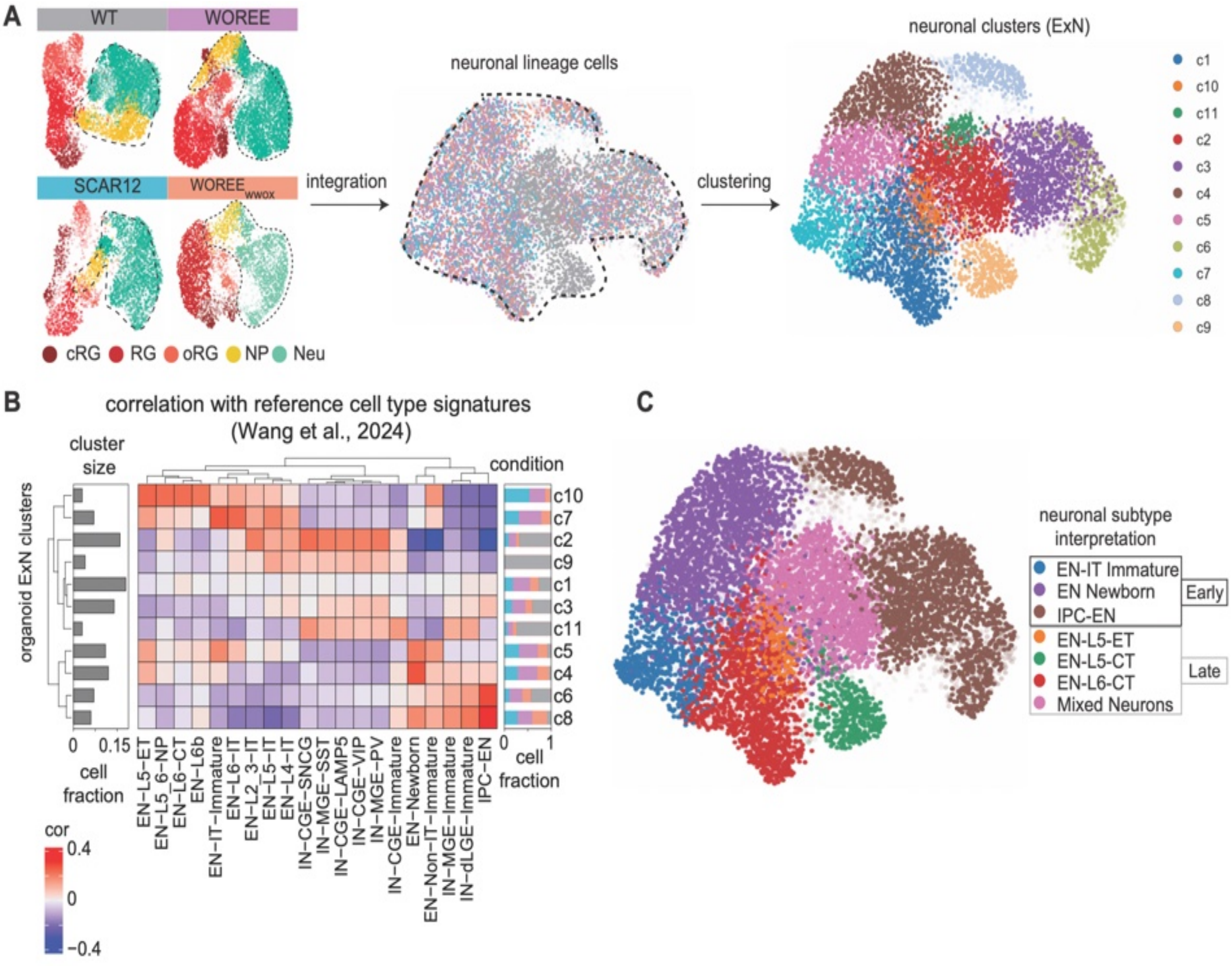


**Fig. EV8. Neuronal cell types annotation. (A)** (Left) Isolation of neurons from WT, WOREE, SCAR12 and WOREE-WWOX organoids. (Right) UMAP plot coloured by condition. **(B)** Clustering of neuronal cells recovering 11 clusters. **(C)** Heatmap of correlations of cluster signatures to reference neuronal cell types (10.1101/2024.01.16.575956). The barplot on the left shows the size of each cluster, while the stacked barplot on the right represents how many cells populate each cluster divided by condition. **(D)** UMAP visualization coloured by neuronal subtypes recovered from (C). The subtypes are assigned based on best correlation score with reference signatures, and divided in early and late classes based on the age these subtypes are found in the reference dataset. The mixed neuronal population is assigned depending on the multiple high scores with both interneuron and excitatory neuron cell types, thus presenting a more general neuronal signature.

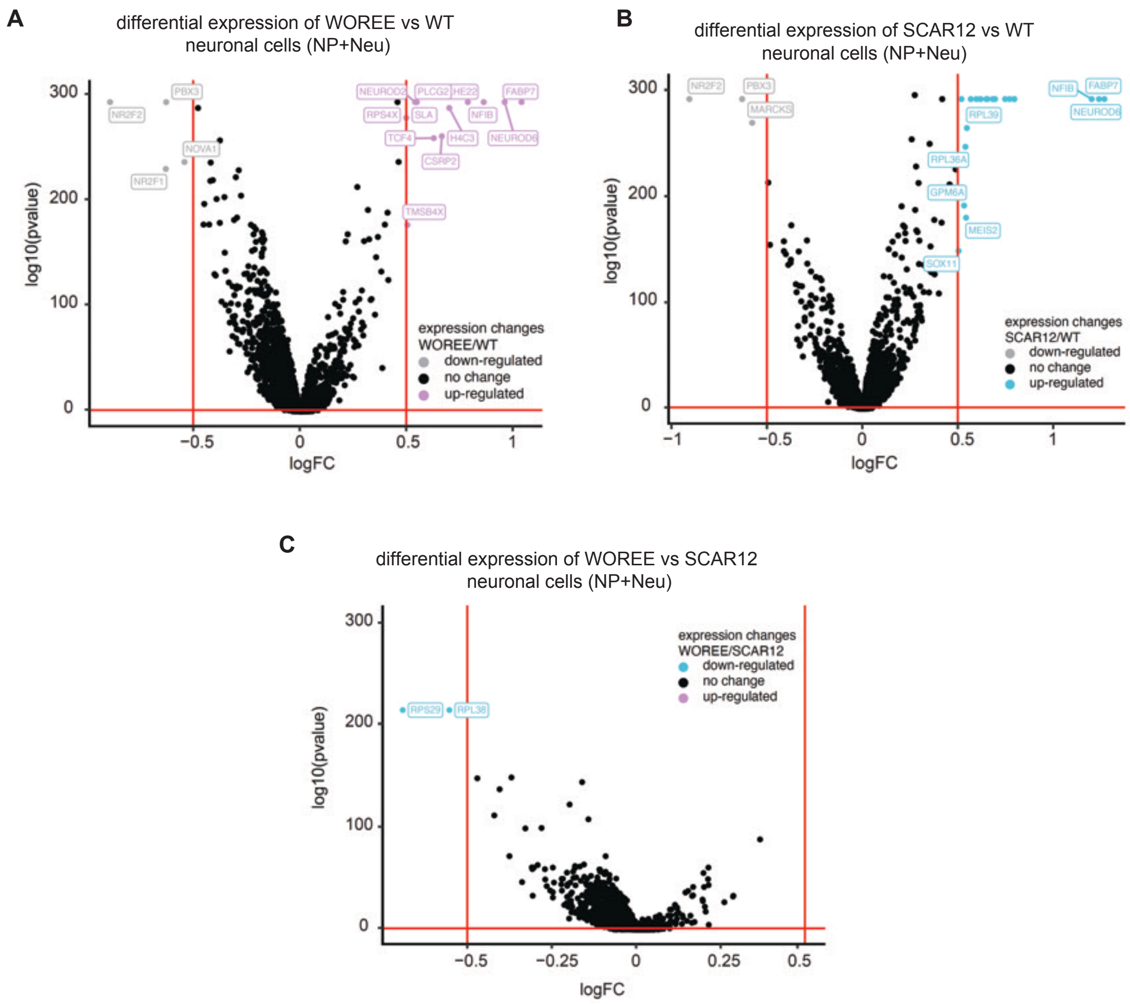


**Fig. EV9. Differentially expressed genes in patient-derived neurons. (A)** Volcano plot showing differential gene expression between WOREE and WT neurons cells, upregulated genes in WOREE are shown in lilla, downregulated genes in the WOREE in grey, not significant genes are in black. **(B)** Differential gene expression between SCAR12 and WT neurons. Upregulated genes in SCAR12 are shown in light blue, downregulated genes in the SCAR12 in grey, not significant genes are in black. **(C)** Differential gene expression between WOREE and SCAR12 neurons. Upregulated genes in WOREE are shown in lilla, Upregulated genes in SCAR12 are shown in light blue, not significant genes are in black.

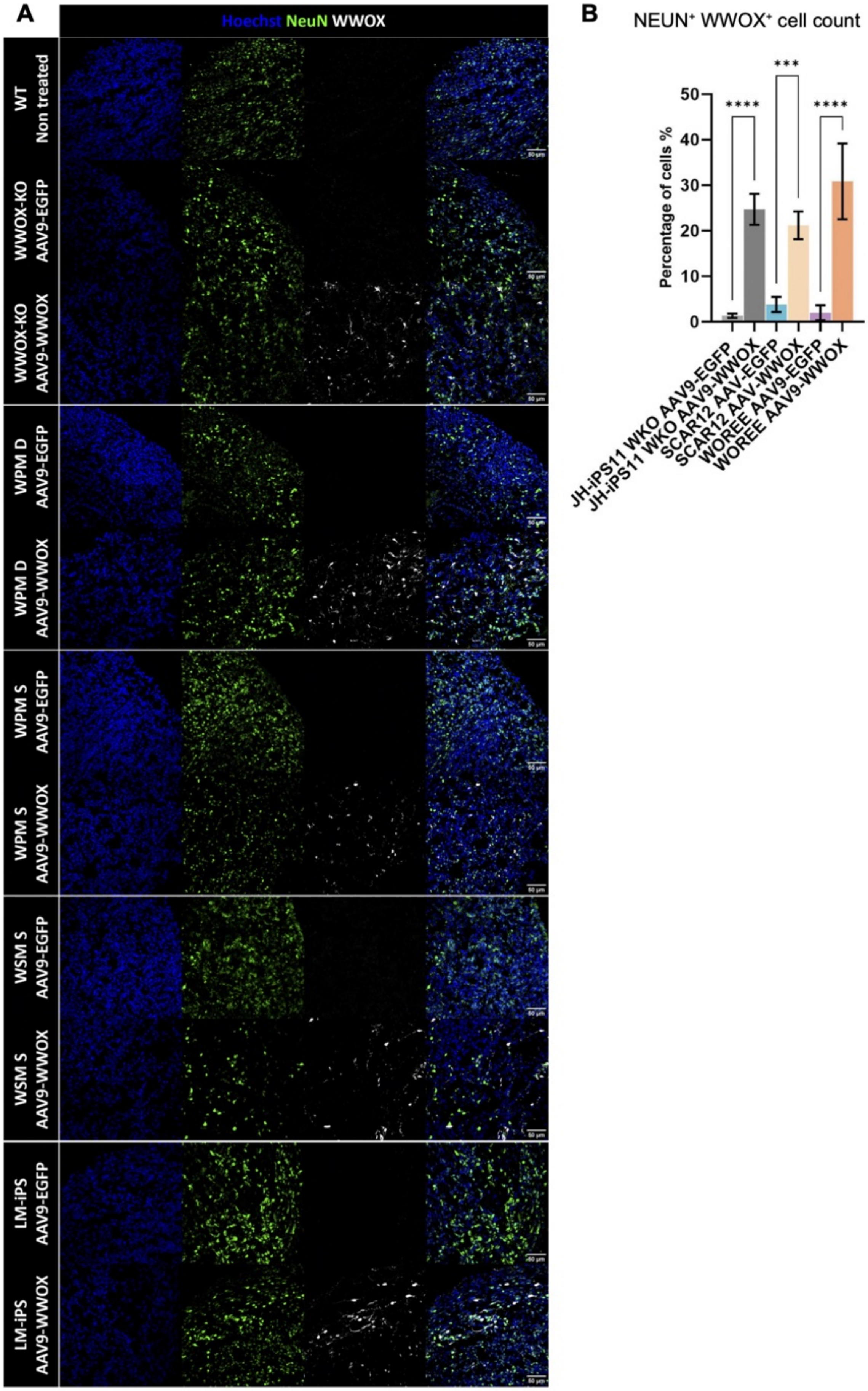
A
Hoechst NeuN WWOX
WT
Non treated
WWOX-KO
AAV9-EGFP
WWOX-KO
AAV9-WWOX
WPM D
AAV9-EGFP
WPM D
AAV9-WWOX
WPM S
AAV9-EGFP
WPM S
AAV9-WWOX
WSM S
AAV9-EGFP
WSM S
AAV9-WWOX
LM-iPS
AAV9-EGFP
LM-iPS
AAV9-WWOX
B
NEUN⁺ WWOX⁺ cell count
Percentage of cells %
0
10
20
30
40
50
****
***
****
JH-iPS11 WKO AAV9-EGFP
JH-iPS11 WKO AAV9-WWOX
SCAR12 AAV-EGFP
SCAR12 AAV-WWOX
WOREE AAV9-EGFP
WOREE AAV9-WWOX

**Fig. EV10. AAV9-WWOX infection in cerebral organoids restores neuronal WWOX expression. (A)** Immunostaining of week 16 COs infected with AAV9-WWOX showing the expression of the neuronal marker NeuN and WWOX per cell line, validating the neuronal restoration of WWOX to patient-derived organoids through neuronal expression of WWOX. **(B)** Quantification of percentage of NEUN+ WWOX+ cells of images in (A) comparing AAV9-hSyn-hWWOX infected organoids to their respective AAV9-hSyn-eGFP infected mutants. (WT: n= 4 organoids; WPM S AAV9-EGFP: n=4; WPM S AAV9-WWOX: N=4; WPM D AAV9-EGFP: n=4; WPM D AAV9-WWOX: n=4; WSM S AAV9-EGFP: n=4; WSM S AAV9-WWOX: n=4; LM-iPS AAV9-EGFP: n=3; LM-iPS AAV9-WWOX: n=4). *See Table EV1 for details on organoid numbers, sections, and batches. Data are presented as mean ± SEM. Statistical significance was determined using one-way ANOVA with Sidak's multiple comparisons test. n.s (non-significant), *p ≤ 0.05, **p ≤ 0.01, ***p ≤ 0.001, ****p ≤ 0.0001.*

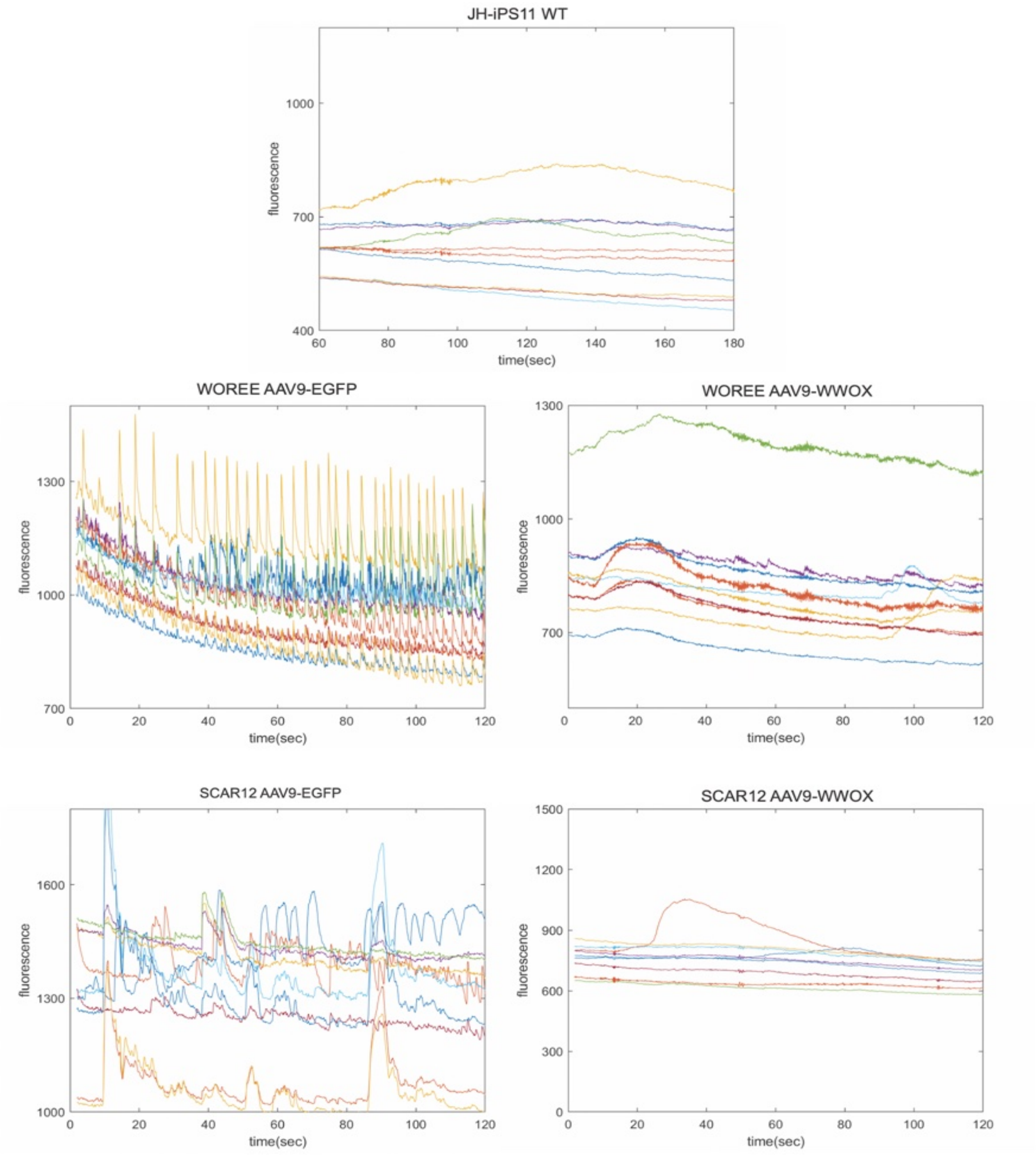


**Fig. EV11. Neuronal restoration of WWOX suppressed neuronal hyperexcitability.** Representative calcium transient in selected neurons recorded in week 16 COs infected either with AAV9-EGFP or AAV9-WWOX of each cell line.

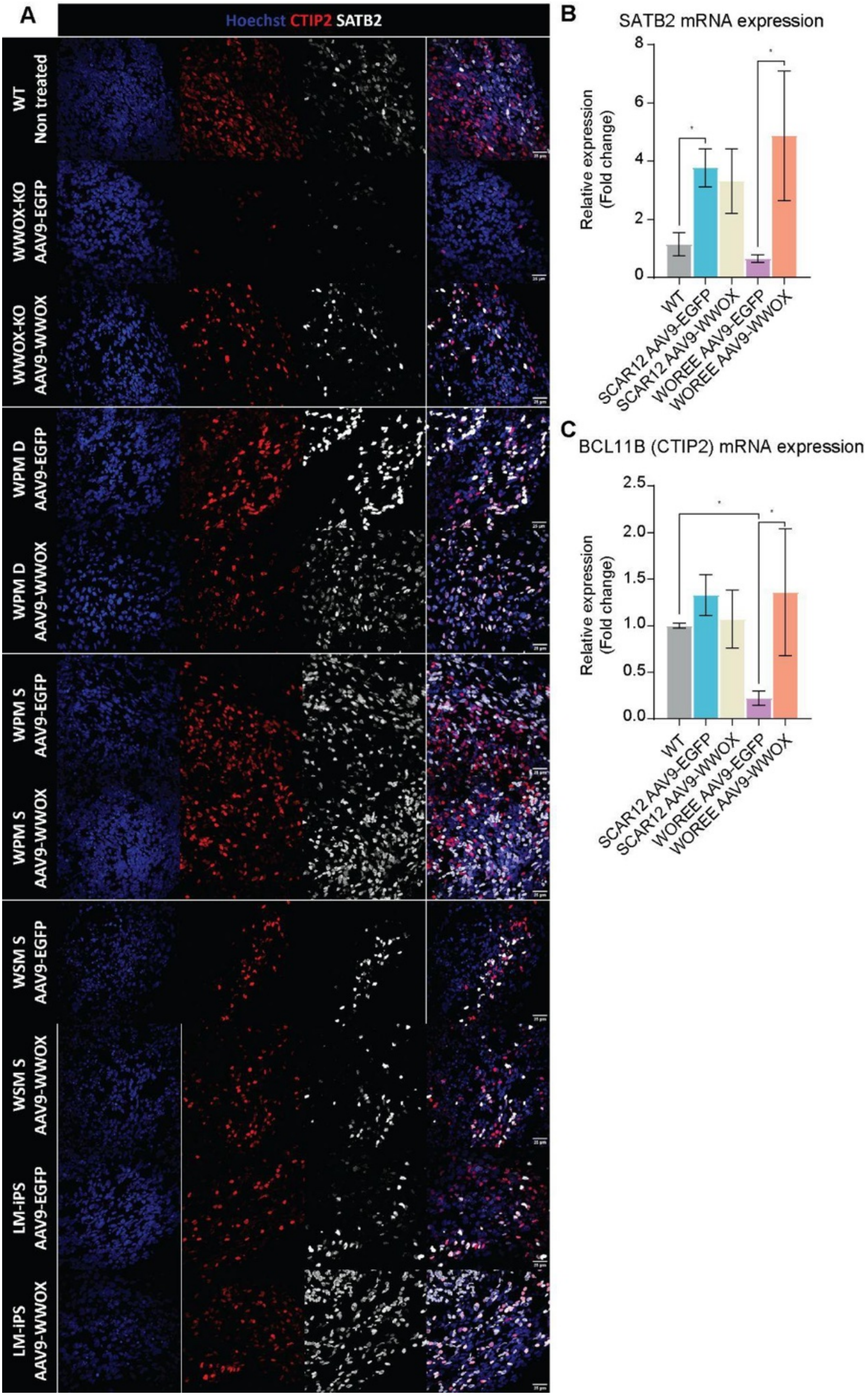
A
Hoechst CTIP2 SATB2
WT Non treated
WWOX-KO AAV9-EGFP
WWOX-KO AAV9-WWOX
WPM D AAV9-EGFP
WPM D AAV9-WWOX
WPM S AAV9-EGFP
WPM S AAV9-WWOX
WSM S AAV9-EGFP
WSM S AAV9-WWOX
LM-iPS AAV9-EGFP
LM-iPS AAV9-WWOX
B
SATB2 mRNA expression
Relative expression (Fold change)
WT
SCAR12 AAV9-EGFP
SCAR12 AAV9-WWOX
WOREE AAV9-EGFP
WOREE AAV9-WWOX
C
BCL11B (CTIP2) mRNA expression
Relative expression (Fold change)
WT
SCAR12 AAV9-EGFP
SCAR12 AAV9-WWOX
WOREE AAV9-EGFP
WOREE AAV9-WWOX

**Fig. EV12. The effect of neuronal WWOX restoration on cortical layers markers.** (**A)** Immunostaining of week 16 COs infected with either AAV9-EGFP or AAV9-WWOX showing the expression of the neuronal markers CTIP2 and SATB2, as an expansion of figure 6, showing all lines included (WT: n= 4 organoids; WPM S AAV9-EGFP: n=4; WPM S AAV9-WWOX: N=4; WPM D AAV9-EGFP: n=4; WPM D AAV9-WWOX: n=4; WSM S AAV9-EGFP: n=4; WSM S AAV9-WWOX: n=4; LM-iPS AAV9-EGFP: n=3; LM-iPS AAV9-WWOX: n=4). **(B)** qPCR analysis of the expression of the superficial layer neuron marker SATB2 in week 16 organoids (WT: n=4 organoids; SCAR12 AAV9-EGFP: n=8; SCAR12 AAV9-WWOX: n=8; WOREE AAV9-EGFP: n=6, WOREE AAV9-WWOX: n=6). Statistical significance was determined by a one-way ANOVA test, correcting for multiple comparisons by controlling the FDR using the Benjamini–Hochberg procedure. **(C)** qPCR analysis of the expression of the deep layer neuron marker CTIP2 (BCL11B) in week 16 organoids (WT: n=4 organoids; SCAR12 AAV9-EGFP: n=8; SCAR12 AAV9-WWOX: n=8; WOREE AAV9-EGFP: n=6, WOREE AAV9-WWOX: n=6). Statistical significance was determined by a one-way ANOVA test, correcting for multiple comparisons by controlling the FDR using the Benjamini–Hochberg procedure.

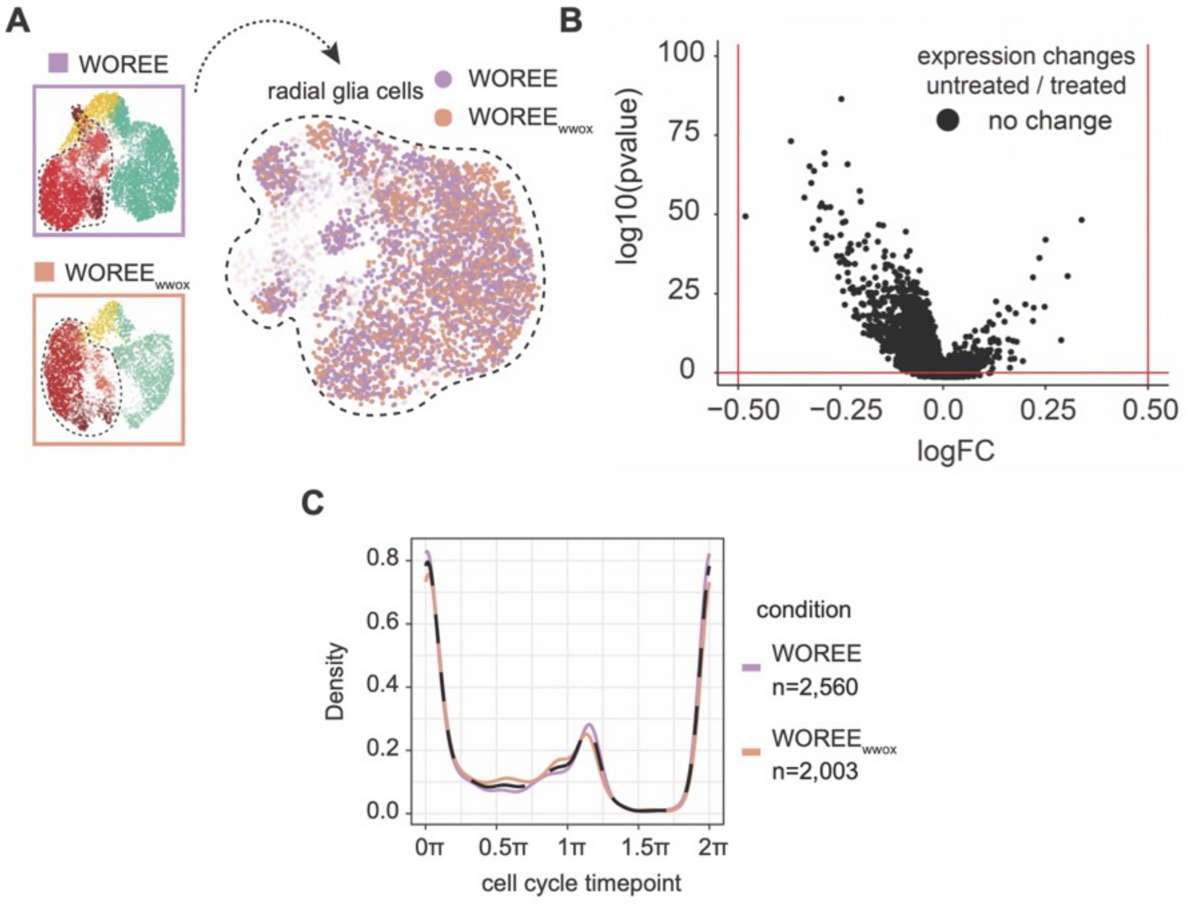


**Fig. EV13. Neuron-targeted gene therapy did not affect the transcriptome and cell-cycle phase of radial glia cells. (A)** Isolation of radial glia cells from untreated (WOREE) and treated (WOREE-WWOX) organoids. On the right, UMAP cell visualization colored by condition. **(B)** Differential gene expression between radial glia in treated vs untreated organoid shows no significant differentially expressed gene between WOREE-WWOX and WOREE radial glia cells. **(C)** Cell cycle inference in radial glia cells showing overlap of WOREE-WWOX and WOREE conditions.
